# Simultaneous in vitro expression of minimal 21 transfer RNAs by tRNA array method

**DOI:** 10.1101/2025.02.15.638384

**Authors:** Ryota Miyachi, Yoshihiro Shimizu, Norikazu Ichihashi

## Abstract

Transfer RNA (tRNA) plays a central role in translation. The simultaneous *in vitro* synthesis of minimal yet sufficient tRNA species (at least 21) poses a challenge for constructing a self-reproducible artificial cell. A key obstacle is the processing of the 5’ and 3’ ends, which requires a multi-step reaction in natural cells. In this study, we developed a simplified processing method that allows simultaneous expression of all 21 tRNAs in a reconstituted transcription/translation system (PURE system). We tested three available methods (leader, 5’-G variants, and HDVR attachment methods) and one new method (direct tRNA linkage method). Using these methods, we succeeded in simultaneous expression of six non-G-start tRNA from monocistronic six DNA templates in the PURE system. Furthermore, we developed a method that combines the direct tRNA linkage and HDVR attachment methods (termed tRNA array method). Using this method, we succeeded in simultaneous expression of all 21 tRNAs from a single polycistronic DNA template in the PURE system. The tRNA mixture produced by the tRNA array method supported a similar level of translation to the individually synthesized tRNA mixture. Additionally, we demonstrated that the minimal tRNA sets prepared by the tRNA array method can be used for genetic code engineering. This study represents a step toward the realization of self-reproducible artificial cells and also provides an easy method for preparing all tRNAs useful for genetic code engineering.

## Introduction

In the field of bottom-up synthetic biology, researchers have focused on creating artificial molecular systems *in vitro* to deepen our understanding of living systems (1–16). One of the major goals is to construct artificial systems with self-reproductivity, a universal ability of living organisms, using minimal components. To date, many efforts have been made to reconstitute molecular systems with functions that contribute to self-reproduction, such as DNA replication utilizing natural (17–20) or artificial schemes (21–25), membrane synthesis (8, 26), metabolic reactions (27–29), and membrane division (30, 31). However, the construction of a transcription/translation (TxTL) system capable of self-regeneration remains a significant challenge (32). To realize such systems, it is necessary to establish a TxTL system that regenerates its components from the DNA encoding all the components. A well-studied reconstituted TxTL system is the PURE system (33), which consists of the *E. coli* translation machinery. In the PURE system, translation proteins (7, 34–36), ribosomal proteins (37, 38), and rRNA (39, 40) are partially synthesized.

tRNA is an essential component of the translation system (41), but its *in vitro* expression scheme is still in progress. In the last few years, the *in vitro* synthesis of unmodified tRNAs has been demonstrated, enabling translation with a reconstituted genetic code (42–47). At least 21 types of tRNAs are required for the translation of arbitrary proteins (20 for each amino acid and one for formyl methionine). In previous studies, each of the 21 tRNA was chemically synthesized or transcribed *in vitro* and purified individually before use. The next challenge toward a self-reproducible translation system is to simultaneously express all 21 tRNA in their functional forms in the PURE system.

A major hurdle for the simultaneous expression of the 21 tRNA in the PURE system is 5’- and 3’- end processing. In *E. coli*, many tRNAs are transcribed as polycistronic premature RNA that also encode either identical tRNAs, unrelated tRNAs, proteins or rRNA. Premature RNA undergoes complicated processing by various RNases, including RNase E, III, T, PH, II, D, P, and PNPase, to produce mature tRNAs with the correct 5′ and 3′ ends (41, 48). This processing reaction has not been fully reconstituted *in vitro*, and the necessity for such a complicated process remains unclear. We think that simpler processing may be possible, at least *in vitro*.

In our previous study, we reported the expression of 15 of 21 tRNAs from monocistronic DNA in a PURE system (49). In the experiment, 5’-end processing was not needed because these 15 tRNAs possess 5’-G, which can be directly transcribed from the 5’ end with T7 RNA polymerase. The 3’- end processing is also not needed because the 3’-end of tRNA was produced by run-off transcription by using template DNA that possesses the 3’-end matching with the 3’-end of the tRNA gene (42). However, the 5’-end method is not applicable to the other six non-G-start tRNAs. In addition, the run-off method for the 3’-end poses a limitation to the 3’-terminal sequence of the template DNA. This 3’-end limitation should be removed to achieve a self-reproducible artificial system because with this limitation, the DNA template is difficult to replicate by currently reconstituted DNA replication schemes, which require circular DNA or linear DNA that contains specific sequences at the termini (21, 22, 50). In addition, for efficient DNA replication, all 21 tRNA should be encoded in a single DNA in a polycistronic manner. Consequently, the next critical challenge in tRNA synthesis for self-reproducing systems is to develop a method for producing all 21 tRNAs with correct 5’- and 3’-ends from a single template DNA with an arbitrary 3’-end sequence in the PURE system.

In this study, we tested some methods for the expression of 21 tRNAs with correct ends using a simpler scheme than that used in *E. coli*. For the six non-G-start tRNAs (Glu, Pro, Ile, Asn, Trp, and fMet), we employed two methods: attachment of a leader sequence, which was removed with RNase P, and mutating the 5’ nucleotide to G to generate 5’G variants. Using these methods, we succeeded in simultaneously expressing all six non-G-start tRNAs coupled with translation from six monocistronic DNA templates in the PURE system. For the 3′-end, we employed two methods: using a self-cleaving ribozyme (HDVR) and direct linkage of tRNAs separated by RNase P. We then found that the combination of these two 3’-end methods (named tRNA array method) also solves the problem of 5’-end processing. Using this tRNA array method, we succeeded in simultaneously expressing all 21 tRNAs from a single polycistronic DNA template in the PURE system. We also demonstrated that tRNA prepared using the tRNA array method is useful for genetic code engineering.

## Results

### Preparation of tRNA-free PURE system (tfPURE system)

In this study, we aimed to express tRNAs from DNA and use them for translation in the PURE system (Fig. 1A). For this purpose, a tRNA-free PURE system (tfPURE system) is required. We previously found that two components of the PURE system, EF-Tu and ribosomes, contain non-negligible levels of tRNA (49). To remove residual tRNA, we re-purified EF-Tu and ribosomes. However, the repurification process for ribosome, the separation of 50S and 30S subunits through sucrose gradient ultracentrifugation, was both laborious and low in yield. To solve this problem, we used a new and simpler method using a size-separation spin column, allowing the preparation of tRNA-free ribosomes at a higher yield (Supplementary Fig. S1).

**Fig. 1.**
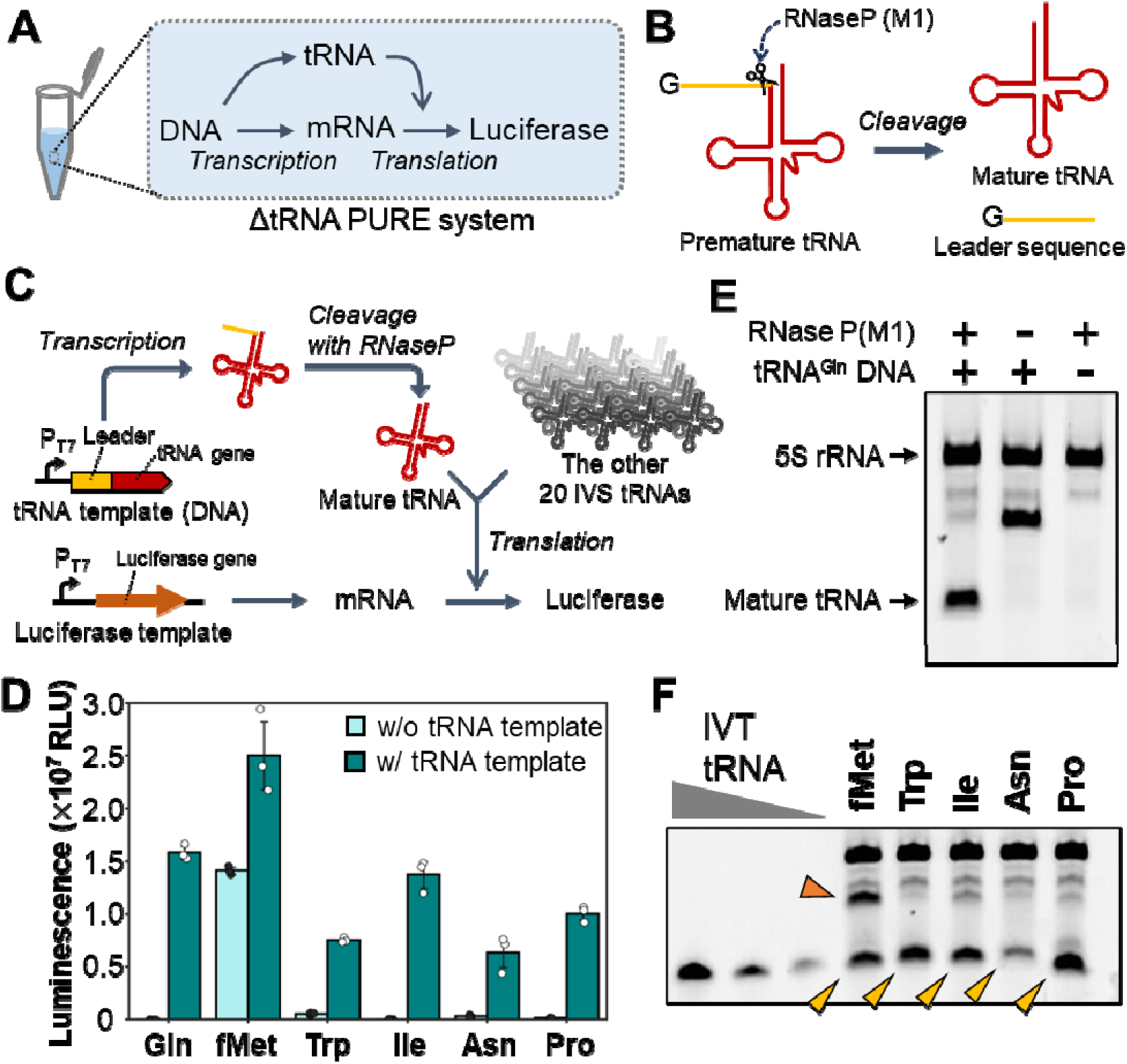
Leader method for 5’-end processing. (A) General scheme of tRNA synthesis coupled with translation in the tfPURE system. The reaction mixture contains tRNA-encoding DNA template (tRNA template) and luciferase-encoding DNA template (luciferase template). The luciferase is translated depending on the synthesized tRNA. (B) Scheme of 5’-end cleavage by RNase P, using the leader method. A premature tRNA is synthesized with a leader sequence at the 5’end. The leader sequence is removed by RNase P to produce a mature tRNA. M1 RNA, the catalytic RNA component of RNase P, was used for this experiment. (C) Specific scheme for non-G-start tRNA processing in the tfPURE system. One of the six non-G-start tRNAs was expressed from a tRNA template and cleaved with RNase P. The resultant mature tRNA is used for the translation of luciferase with the other 20 IVS tRNAs. The 3’ end of tRNA was prepared by run-off transcription. (D) Luciferase activity in the translation-coupled reaction of expressed tRNA. The reaction mixture contained each of six tRNA templates (50 nM), luciferase template (1 nM), RNase P (M1 RNA, 500 nM), T7 RNA polymerase (1.7 U/μL), 20 IVS tRNAs (excluding the tRNA to be expressed), and the tfPURE system (composition A, Supplementary Table 6). After incubation at 30°C for 16 h, luminescence was measured. Each data point represents three independent experiments, and error bars indicate standard deviations. (E) PAGE analysis after the translation-coupled reaction of tRNA^Gln^. The reaction was conducted as shown in (D), except for the omission of 20 IVS tRNAs. RNA was stained with SYBR Green II. (F) PAGE analysis of the other five tRNAs using the same method as described in (E), except for the omission of 20 IVS tRNAs. The expected bands for mature tRNA are indicated by yellow arrowheads.

### Leader method for 5’-end processing: attachment of leader sequence processed with RNase P

To enable the expression of six non-G-start tRNAs that cannot be directly synthesized using T7 RNA polymerase, we first evaluated the method of attaching a 5’-G leader sequence to the 5’-end and cleaving it with RNase P (51)(Fig.1B). Although this method has been previously used for *in vitro* transcription of tRNA (42, 46), it has not yet been applied to tRNA synthesis in the PURE system. To examine whether this method works for the translation-coupled reaction in the PURE system, we performed the reaction in the tfPURE system containing a tRNA template DNA encoding one of the six non-G start tRNAs, the other 20 *in vitro* synthesized (IVS) tRNA, RNase P, and a luciferase DNA template at 30°C for 16 h. The 3’-terminus of the tRNA template matched the 3’-end of each tRNA for the run-off transcription. If this method works as expected, premature tRNA with the leader sequence is synthesized from the tRNA template and cleaved by RNase P to produce a mature tRNA with the correct 5’-end. The mature tRNA works with the other 20 tRNAs to translate luciferase, which is used as a reporter for translation activity (Fig. 1C). Native RNase P consists of M1 RNA and C5 protein, but we used only M1 RNA component in this experiment because M1 RNA is sufficient for RNase P activity *in vitro* (52, 53).

First, we conducted this reaction using a tRNA template encoding tRNA^Gln^ in the tfPURE system that contains 20 other IVS tRNAs. The concentrations of NTP, Mg(OAc)_2_, and spermidine were optimized before the experiment (Supplementary Fig. S2). Luciferase activity increased to 10^7^, depending on the tRNA template (Fig. 1D). We then performed experiments with the remaining five tRNAs (tRNA^Pro^, tRNA^Ile^, tRNA^Asn^, tRNA^Trp^, and tRNA^fMet^) and detected similar levels of luciferase activity in the presence of the tRNA template. These results indicate that functional tRNAs were produced by this method. We also observed higher background luciferase activity for tRNA^fMet^ in the absence of the tRNA template. This might be attributed to undetectable amounts of tRNA^fMet^ remaining in the tfPURE system (49) or other elongator tRNA (mMet or other) involved in translation initiation as previously reported (54).

Next, we directly checked the cleavage of the leader sequence in the tfPURE system without adding other 20 IVS tRNAs, using electrophoresis. For tRNA^Gln^, we detected a band corresponding to mature tRNA depending on RNase P (Fig. 1E). Similar bands were detected for the other five tRNAs (Fig. 1F, yellow arrowheads). Quantification based on band intensity showed that synthesized tRNAs ranged from 16 to 35 ng/µL (Supplementary Fig. S3), similar to or greater than the average tRNA concentration used for IVT tRNA (11.7 ng/µL) in a previous study (49). For tRNA^fMet^, a relatively large amount of uncleaved premature tRNA was detected (Fig. 1F, orange arrowheads), indicating that cleavage efficiency varies depending on the tRNA sequence.

### 5’ variant method: introducing G mutations to the 5’-end

As a second method to prepare tRNA with a correct 5’-end, we substituted the non-G nucleotide at the 5’-end with G to generate 5’-G variants (Fig. 2A). We also designed 5’-A variants for tRNA^fMet^ because T7 RNA polymerase can initiate RNA synthesis from A when using the class II T7 promoter (55). Since the structure of the acceptor stem is known to be critical for tRNA functionality, we also introduced mutations into the complementary base sequences (Fig. 2A, Supplementary Table 4). Some of these variants (Trp: G|U and G|C, Gln: G|A and G|C, Ile: G|C, Pro: G|C and G|G, fMet: G|A and G|U) have been analyzed in previous studies on aminoacylation activity (56–59) or formylation activity (60). Furthermore, tRNA^Asn^(G|C), tRNA^Ile^(G|C), and tRNA^fMet^(G|A) have been used for translation in reconstituted systems (43, 46, 47). To evaluate the translation activities of these tRNAs in our system, we first added each purified IVS tRNA variant to the tfPURE system containing the other 20 IVS tRNAs and a luciferase template. After incubation at 30°C for 16 h, luciferase activity was measured. Most tRNA variants exhibited similar or higher luciferase activity than the wild-type tRNA (WT) (Fig. 2B), with the exception of one variant, tRNA^Pro^ (G|C). The lack of activity for this variant is consistent with the previous aminoacylation result (59). Next, we selected the tRNA variant with the highest activity for each amino acid (Fig. 2B, red triangles), and conducted tRNA synthesis coupled with luciferase translation in the tfPURE system (Fig. 2C). Luciferase activity was detected for all the variants tested, depending on the tRNA template (Fig. 2D). For tRNA^fMet^, a higher background luciferase activity was detected without the tRNA template. The luciferase activity of tRNA^Asn^ was much lower than those of the other tRNAs.

**Fig. 2.**
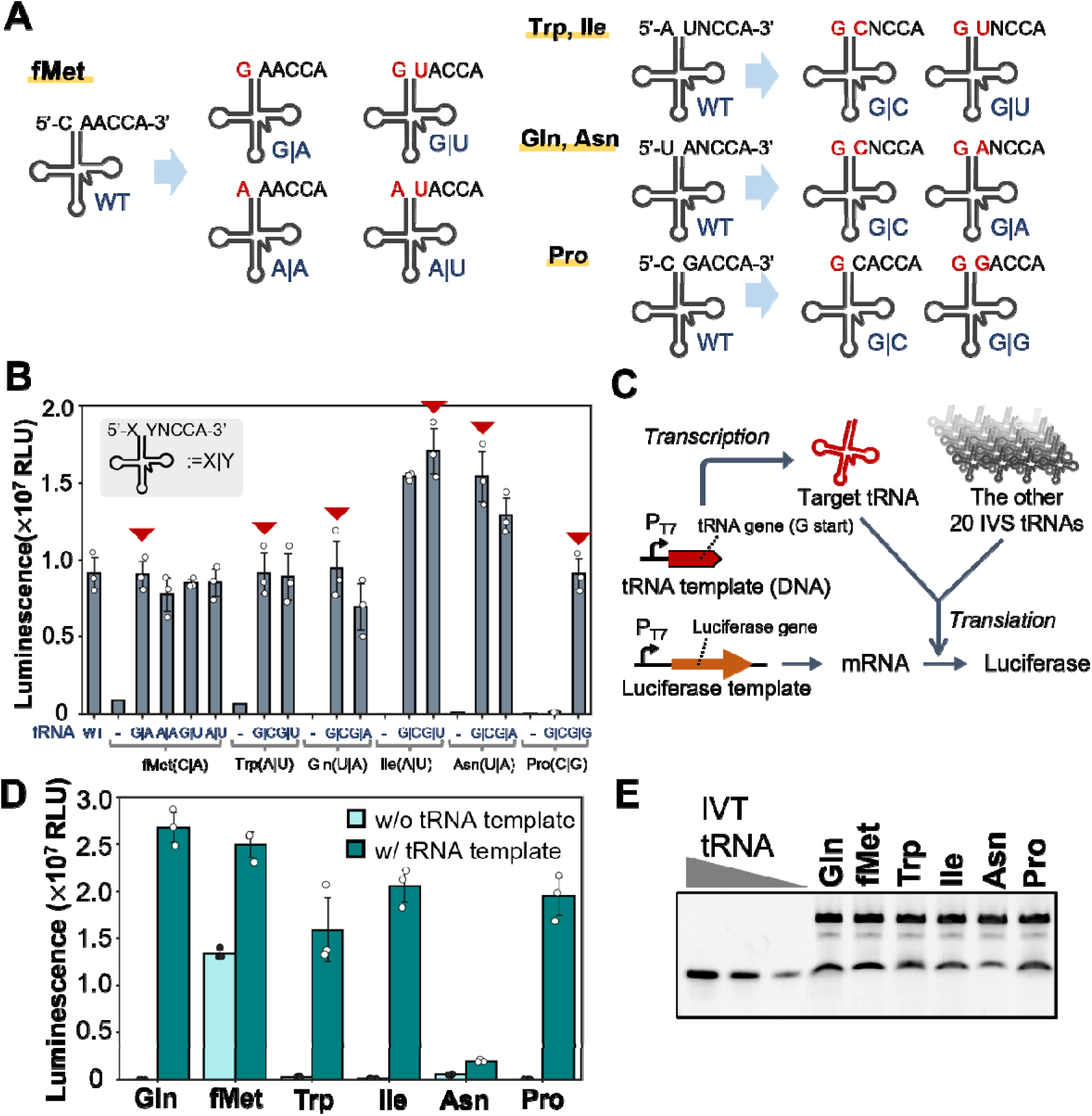
5’-G variant method for 5’-end of tRNA. (A) Design of 5’-G or A variants. These tRNA variants were designed by substituting bases at the 5’-end with their paired bases in the acceptor stem. The variants were named using the base-pairing combination after substitution (e.g., G|A). (B) Luciferase activity after translation with purified IVS tRNA variants. Each tRNA variant synthesized in vitro and purified was added to the tfPURE system (composition A, Supplementary Table 6), together with the other 20 IVS tRNAs, and their translation activity was assessed using luciferase expression. Original wild-type tRNAs (WT) and a reaction without tRNA variants (−) were used as controls. (C) Scheme for the expression of 5’-G variants coupled with translation in the tfPURE system. One of the six tRNA variants was expressed from a tRNA template and was used for the translation of luciferase with the other IVS 20 tRNAs. The 3’ end of the tRNA was prepared by run-off transcription. (D) Luciferase activity in the translation-coupled reaction of the expressed tRNA. The reaction mixture contained a tRNA template for one of the 5’-G variants indicated by the red arrows in (B) (50 nM), the luciferase template (1 nM), T7 RNA polymerase (1.7 U/μL), 20 IVS tRNAs (excluding the tRNA to be expressed), and the tfPURE system(composition C, Supplementary Table 6). After incubation at 30°C for 16 h, luminescence was measured. (E) PAGE analysis. The reaction was conducted as shown in (D), except for omitting 20 IVS tRNAs, and subjected to PAGE analysis. RNA was stained with SYBR Green II. Each data point in (B) and (D) represents three independent experiments, and error bars indicate standard deviations.

The synthesized tRNAs were analyzed by electrophoresis. We detected a band of the expected size for all the tRNAs (Fig. 2E). Quantification based on band intensity showed that synthesized tRNAs, except for tRNA^Asn^, ranged from 22 to 35 ng/µL (Supplementary Fig. S4), which was comparable to the amount obtained using the previous method with the leader sequence. However, the yield of tRNA^Asn^ (6.6 ng/µL) was significantly lower than that of the other tRNAs for unknown reasons.

To determine which method to use in the next simultaneous expression experiment, we compared the translation activity of tRNAs prepared by the leader and 5’-G variant methods described above (Supplementary Fig. S5). These two methods were sufficiently useful for most tRNAs, except for tRNA^Asn^, for which the 5’G variant method produced a much lower expression level than the leader method. We also found that a relatively large amount of uncleaved premature tRNA^fMet^ remained when using the leader method. Based on these results, we decided to use the leader method for tRNA^Asn^, the 5’G variant method for tRNA^fMet^, and both methods for the other tRNAs in the subsequent simultaneous expression experiment.

At this point, we examined the effect of the 5’-end phosphate on translation. tRNAs synthesized using the 5’-G variant method are expected to contain a triphosphate group at the 5’-end. In contrast, tRNAs synthesized by the leader method are expected to contain 5’-monophosphate because of the RNase P treatment. To investigate whether this difference affects translation, we prepared 5’-triphosphate or monophosphate tRNA^His^ and tRNA^Ala^ using simple *in vitro* transcription or the leader method. Comparison of luciferase activity with these tRNAs revealed that the translation activities of 5’-monophosphates were only slightly lower than those of 5’-triphosphates for both tRNA^His^ and tRNA^Ala^ (Supplementary Fig. S6).

### Simultaneous six tRNA synthesis from monocistronic DNAs

We simultaneously synthesized all six non-G-start tRNAs from the six tRNA templates in the tfPURE system and evaluated their functionality in luciferase translation (Fig. 3A). The leader and 5’-G variant methods were adopted for tRNA^Asn^ and tRNA^fMet^, respectively, and both methods were compared for the remaining four tRNAs (Fig. 3B). The luminescence with tRNA templates for both methods were significantly higher than that without tRNA templates, reaching more than 0.6×10^7^. This value indicated that all the five tRNAs (for Gln, Ile, Pro, Trp, and Asn) were sufficiently expressed. Notably, using the 5’-G variant method for the four tRNAs (Gln, Ile, Pro, and Trp) yielded higher luciferase activity (1.3×10^7^) than the leader method, indicating that the 5’-G variant method is more useful for maximizing tRNA expression from monocistronic DNA templates in the PURE system.

**Fig. 3.**
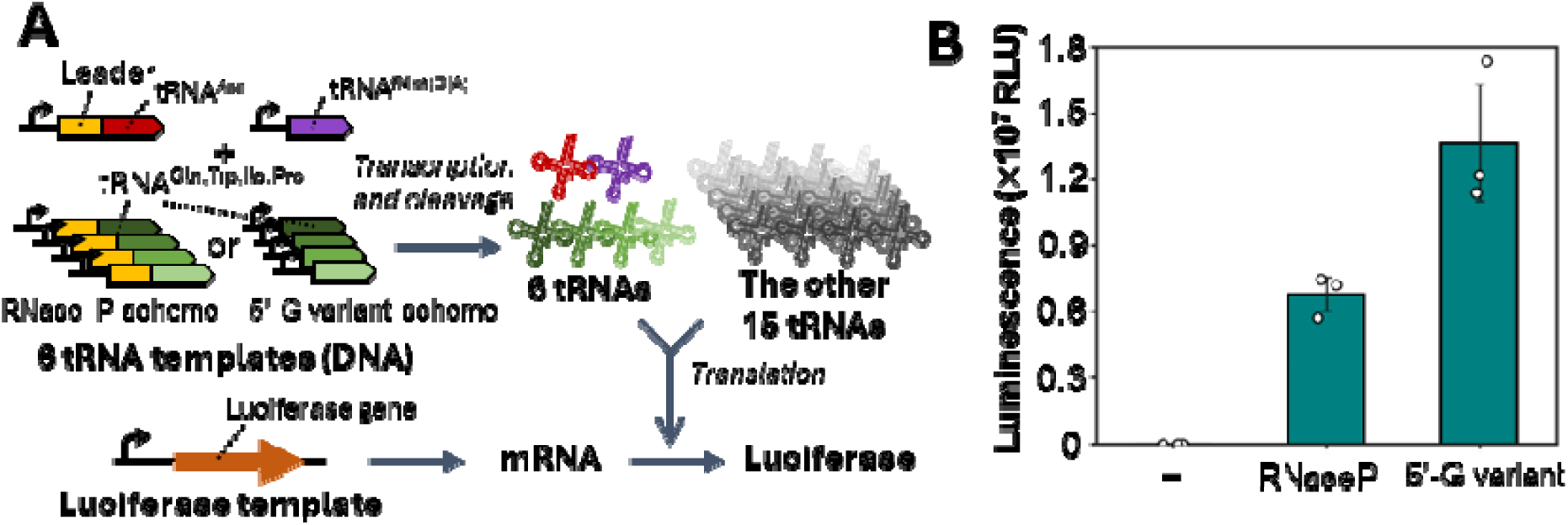
Simultaneous expression of tRNAs from monocistronic tRNA templates. (A) Scheme of the simultaneous expression of six non-G-start tRNAs coupled with translation. Six tRNAs were synthesized from the corresponding tRNA templates in the tfPURE system. The synthesized tRNAs were used for luciferase translation with other IVS 15 tRNAs. The leader and 5’-G variant methods were adopted for tRNA^Asn^ and tRNA^fMet^, respectively. Both methods were tested using four other tRNAs (tRNA^Gln, Ile, Pro, and Trp^). (B) Luciferase activity in the translation-coupled reaction for six tRNA co-expression. The reaction mixture contained six tRNA templates (15 nM each), luciferase template (1 nM), RNase P (M1 RNA, 500 nM), T7 RNA polymerase (1.7 U/μL), the other 15 IVS tRNAs, and tfPURE system (composition B, Table S6). The 3’ ends of the tRNAs were prepared by run-off transcription. The reaction mixture was incubated at 30°C for 16 h, and luminescence was measured. The methods used for the four tRNAs (tRNA^Gln, Ile, Pro, and Trp^) are shown.

### HDVR method for 3’-end processing: introducing self-cleavage ribozyme (HDVR)

In the previous experiment for simultaneous 21 tRNA synthesis, the 3’-terminus of the RNA template must match the 3’-end of the tRNA gene for run-off transcription. Accordingly, the tRNA template must be monocistronic (i.e., one DNA template encodes only one tRNA gene). To overcome this limitation and allow tRNA synthesis from polycistronic DNA, as in the cell, we explored two 3’-end processing methods. The first method utilized the self-cleaving ribozyme HDVR, as previously reported (61). We placed the HDVR immediately downstream of the tRNA gene, followed by a strong T7 terminator, T7 hyb10 (62) (Fig. 4A). During transcription, the premature tRNA connected with HDVR and terminator is synthesized and then undergoes self-cleavage of HDVR, generating a tRNA with a 2’, 3’ cyclic phosphate at its 3’-end. Since this cyclic phosphate cannot be aminoacylated by aminoacyl-tRNA transferases, it is dephosphorylated by T4 polynucleotide kinase (PNK) (63).

**Fig. 4.**
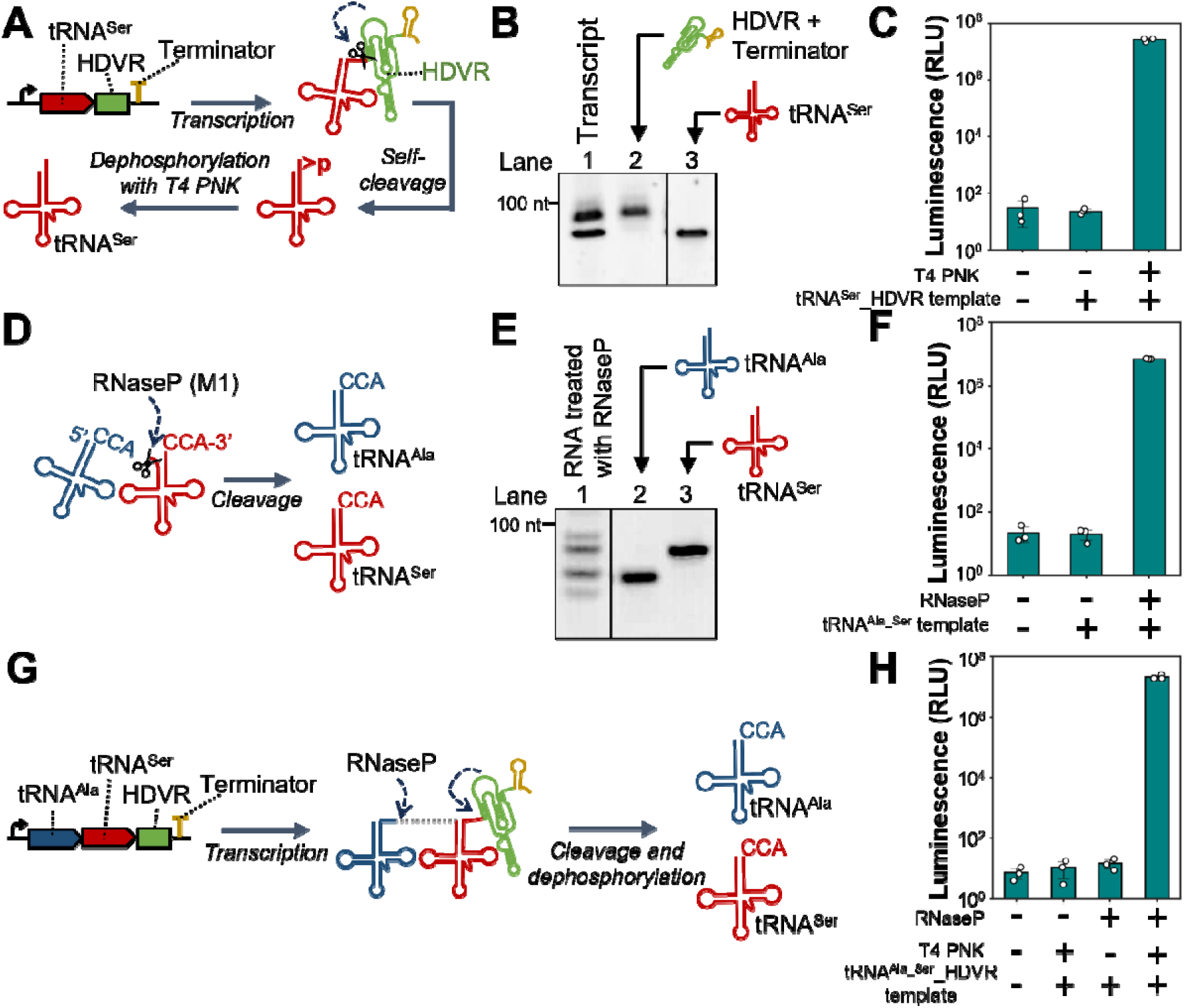
HDVR and linked tRNA methods for 3’-end processing. (A) Scheme of the HDVR method. HDVR ribozyme is placed directly downstream of the tRNA gene with a terminator (T7 hyb10). The resulting 2’,3’-cyclic phosphate is removed by T4 PNK, yielding mature tRNAs. (B) PAGE analysis after reaction shown in (A). An aliquot of the *in vitro* transcription reaction with template DNA and T4 PNK was analyzed (lane 1, Transcript). Products synthesized from a template containing only HDVR and terminator (lane 2) and tRNA^Ser^ (lane 3) were used as controls. (C) Luciferase activity in translation-coupled reactions by using the HDVR method. The reaction mixture contained the tRNA template with HDVR (50 nM), luciferase template (1 nM), T7 RNA polymerase (1.7 U/μL), T4 PNK (0.094 U/μL), 20 other IVS tRNAs (excluding tRNA^Ser^), and the tfPURE system (composition D). The reaction mixture was incubated at 30°C for 16 h, and luminescence was measured. (D) Scheme of the linked-tRNA method. RNase P separates the directly linked tRNAs and simultaneously processes the 3’- and 5’-ends of the separated tRNAs. (E) PAGE analysis of the linked tRNA treated with RNase P. The linked tRNAs prepared by *in vitro* transcription (250 nM) and RNase P (M1 RNA) (500 nM) were incubated at 37°C for 16 h in buffer R (50 mM Tris HCl(pH7.6), 60 mM NH_4_Cl, 10 mM Mg(OAc)_2_, and 5 mM spermidine), and an aliquot (1 µL) was applied (lane 1). Purified IVS tRNA^Ala^ (lane 2) and tRNA^Ser^ (lane 3) were used as controls. The entire image is shown in Supplementary Fig. S7. (F) Luciferase activity in a translation-coupled reaction using the linked tRNA method. The reaction mixture contained the DNA template encoding the linked tRNA^Ala-Ser^, luciferase template (1 nM), T7 RNA polymerase (1.7 U/μL), 500 nM RNase P (M1 RNA), other 19 IVS tRNAs (excluding tRNA^Ala^ and tRNA^Ser^), and the tfPURE system (composition B). The 3’-end was determined by run-off transcription. The reaction was conducted at 30°C for 16 h, and the luminescence was measured. (G) Scheme of the combination of HDVR and linked tRNA method. Premature linked tRNA^Ala-Ser^ -HDVR expressed from DNA is subjected to a series of processes: self-cleavage of HDVR, dephosphorylation by T4 PNK, and digestion with RNase P to produce two mature tRNAs. (H) Luciferase activity in a translation-coupled reaction using the combined method. The reaction mixture contained the DNA template encoding tRNA^Ala-Ser^-HDVR (50 nM), luciferase template (1 nM), T7 RNA polymerase (1.7 U/μL), T4 PNK (0.094 U/μL), RNase P (M1 RNA) (500 nM), other 19 IVS tRNAs, and the tfPURE system (composition E). The reaction was incubated at 30°C for 16 h, and luminescence was measured. Each data point in (C), (F), and (H) represents three independent experiments, and error bars indicate standard deviations.

To test the self-cleavage ability of premature tRNA^Ser^-HDVR, we first performed *in vitro* transcription in standard buffer (see Methods) and subjected the transcript to PAGE analysis. The transcript exhibits two main bands (Fig. 4B, lane 1), each of which corresponds to the cleaved-HDVR (lane2) and mature tRNA^Ser^ (lane 3), respectively, supporting successful self-cleavage. Next, we investigated whether the tRNA^Ser^ generated through self-cleavage is functional for translation in the tfPURE system. We conducted the reaction shown in Fig. 4A in the tfPURE system containing the tRNA^Ser^-HDVR DNA template, the other 20 IVS tRNAs, and a luciferase template at 30°C for 16 h, and luciferase activity was measured. The luminescence increased to ∼10^7^ depending on both the tRNA template and T4 PNK (Fig. 4C), supporting the functionality of tRNA^Ser^ generated through the self-cleavage of HDVR and dephosphorylation with T4 PNK.

### Linked tRNA method for 3’- and 5’-ends processing

As the second method for 3’-end processing, we devised another strategy utilizing RNase P for the simultaneous processing of the 3’-end of a tRNA and the 5’-end of another tRNA. RNase P recognizes the structure of a tRNA (64) and cleaves the specific site between the 5’-leader sequence and subsequent tRNA. Based on the weak sequence requirement for the 5’-leader sequence (65) (66), we hypothesized that if the 5’-leader RNA was replaced with another tRNA, RNase P could cut between the two tRNAs, allowing simultaneous processing of the 3’-end of the former tRNA and the 5’-end of the latter tRNA (Fig. 4D). To test this hypothesis, we prepared a substrate RNA (tRNAs^Ala-Ser^) composed of directly linked tRNA^Ala^ (76 nt) and tRNA^Ser^ (88nt) and then treated it with RNase P (M1) in standard buffer for RNase P (52)(see legend). The RNA product (Fig. 4E, Supplementary Fig. S7) exhibited two major bands (lane 1), corresponding to either mature tRNA^Ala^ (lane 2) or tRNA^Ser^ (lane 3), supporting the successful cleavage of tRNAs.

Encouraged by this result, we expressed linked tRNAs^Ala-Ser^ from the DNA template in the tfPURE system and processed it in the same reaction mixture. The reaction mixture contained a DNA template that encoded tRNAs^Ala-Ser^, the other 19 tRNAs, RNase P (M1), a luciferase template, and the tfPURE system. In this experiment, the 3’-end of tRNAs^Ala-Ser^ was determined by run-off transcription. After incubation at 30°C for 16 h, luciferase activity was measured. The luminescence increased to ∼10^7^ depending on both the tRNA^Ala-Ser^ template and RNase P, supporting the production of functional tRNAs through digestion with RNase P (Fig. 4F). The reaction mixture was subjected to PAGE (Supplementary Fig. S8). The products exhibited two bands corresponding to each mature tRNAs, although a thick band was detected at the position of uncut tRNA (tRNAs^Ala-Ser^), indicating that there is still room for improvement in digestion efficiency, which was addressed later.

Next, we combined this linked tRNA method (Fig. 4D) with the HDVR method (Fig. 4A) for the simultaneous expression of the two tRNAs without run-off transcription (Fig. 4G). In this experiment, the premature linked tRNA attached to HDVR (tRNAs^Ala-Ser^-HDVR) was expressed from a DNA template and then processed to mature tRNA^Ala^ and tRNA^Ser^ through digestion by RNase P and HDVR self-cleavage followed by dephosphorylation by T4 PNK. We conducted this reaction in the tfPURE system containing the other 19 IVS tRNAs and the luciferase template at 30°C for 16 h and measured luciferase activity (Fig. 4H). The luminescence increased to ∼10^7^ depending on the tRNA template, RNase P (M1), and T4 PNK, indicating that functional tRNAs were produced according to the expected processes.

### Multicistronic tRNA expression using tRNA array methods

The combination of the linked tRNA and HDVR methods can be extended to a greater number of tRNA. We named this combined method for multiple tRNAs as “tRNA array method.” Next, we attempted to express all 21 tRNAs using this method. First, we divided the 21 tRNAs into 3-5 groups (Fig. 5A). The four tRNAs, Ile, Phe, Glu, and Asn (IPEN), are grouped as single operons because they are required at particularly high concentrations for efficient translation (42). The remaining 17 tRNAs were divided into four operons, each containing 3-5 tRNAs, arranged to maintain similar operon sizes. The order of each array was determined as follows. We presumed that the most upstream tRNA would affect transcription because it directly attaches to the promoter. To find tRNAs suitable for this position, we compared the expression levels of some tRNAs (Gly, Asp, Ser, Leu, Ala, and His) (Supplementary Fig. S9), which were expressed at higher levels in our previous study (49). Four tRNAs (Gly, Asp, Ser, and Leu) that exhibited higher transcription levels were assigned to the most upstream positions in each array. The remaining tRNAs were arranged for no specific reason, but mainly alphabetically.

**Fig. 5.**
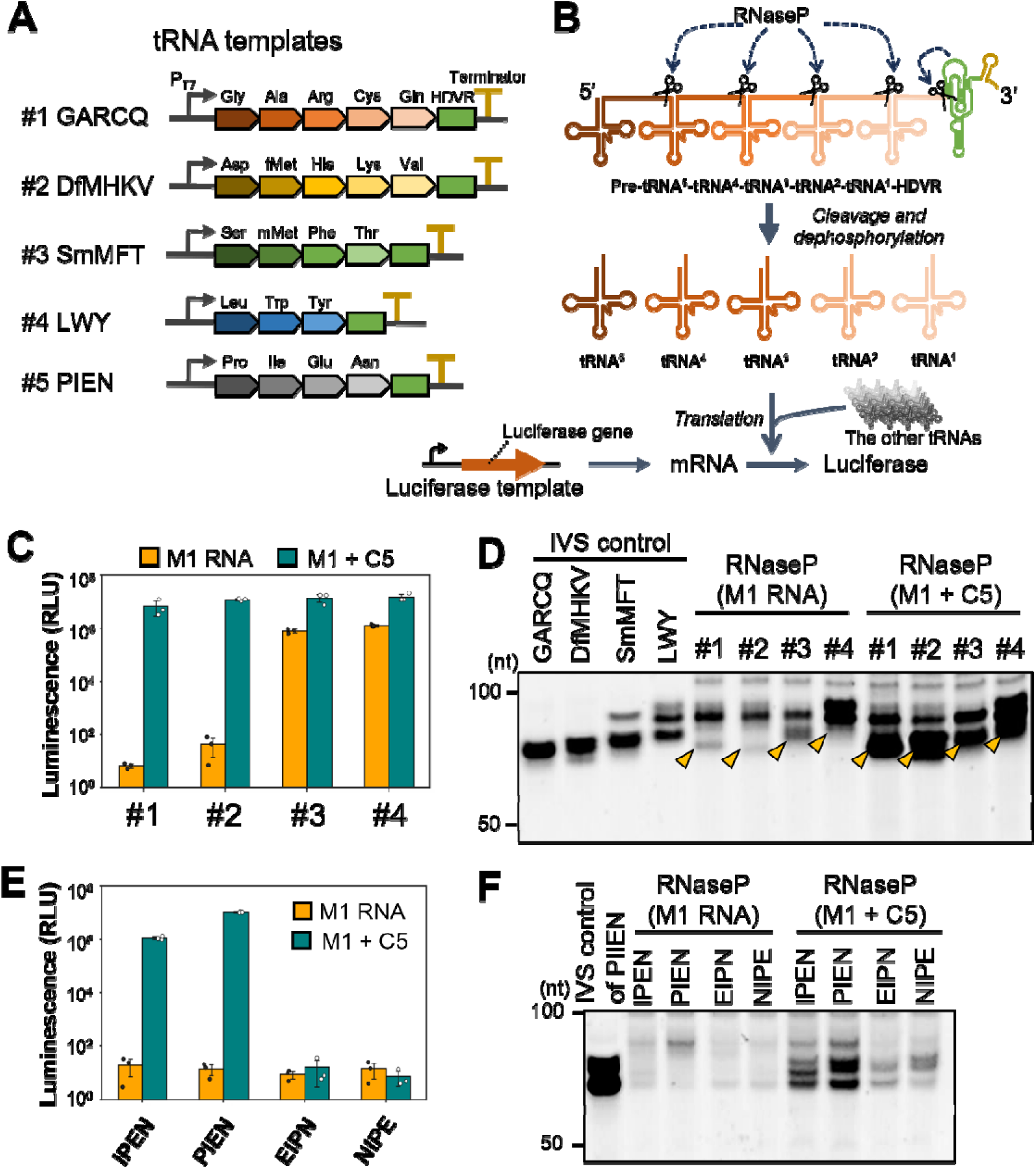
Simultaneous expression 3-5 tRNAs from multicistronic DNA template by the tRNA array method. (A) Design of the tRNA templates. Each template was named according to the corresponding amino acids. (B) Scheme of the tRNA array method and translation-coupled reaction. A premature tRNA consist of linked 3-5 tRNAs and HDVR undergoes a series of processes: self-cleavage of HDVR, dephosphorylation by T4 PNK, and digestion with RNase P to produce 3-5 mature tRNAs, which are used for translation of luciferase. (C) Luciferase activity in the translation-coupled reaction for each tRNA template (#1-4). The reaction mixture contained each tRNAs template (50 nM), luciferase template (1 nM), T7 RNA polymerase (1.7 U/μL), T4 PNK (0.094 U/μL), 16-18 purified tRNAs (excluding the target tRNAs for expression), and the tfPURE system (composition E). For “M1 RNA”, M1 RNA (1 μM) was added as RNase P. For “M1 + C5”, M1 RNA (1 μM) and C5 protein (1.5 μM) was added as RNase P. Incubation was performed at 30 °C for 24 h, and luminescence was measured. (D) PAGE analysis of the reaction mixture incubated in (C). As a size control, a mixture of purified IVS tRNA was used. (E) Luciferase activity in a translation-coupled assay for each tRNAs template encoding Ile, Phe, Glu, and Asn (IPEN), in different orders. This method is the same as that described in (C). (F) PAGE analysis of the reaction mixture incubated in (E). As a size control, a mixture of IVS tRNAs of IPEN was applied. Data points in (C) and (E) represent three independent experiments, and error bars indicate standard deviation.

To evaluate the functionality of tRNA expressed from these tRNA arrays, we performed a translation-coupled reaction in the tfPURE system. During the reaction, multicistronic DNA is transcribed to produce premature tRNA composed of connected 3-5 tRNAs and HDVR, which are then separated into mature tRNAs through digestion by RNase P and self-cleavage of HDVR followed by dephosphorylation by T4 PNK (Fig. 5B). The resultant tRNAs are used for luciferase translation.

We conducted this translation-coupled reaction with each one of the four groups (#1 GARCQ, #2 DfMHKV, #3 SmMFT, and #4 LWY) in the tfPURE system containing RNase P (M1 RNA) at 30 °C for 24 h. The results are shown in Fig. 5C (yellow bars indicated as “M1 RNA”). The luminescence for #3 and #4 reached approximately 10^6^, whereas those for #1 and #2 were less than 10^2^. PAGE analysis of the synthesized tRNAs revealed that the cleaved products corresponding to the expected tRNA sizes were faint for templates #1 and #2 (Fig. 5D, “RNase P (M1)”). These low yields of tRNAs can be caused by insufficient digestion by RNase P of M1 RNA only. To improve this, we supplemented the reaction system with C5 protein, another subunit of E. coli RNase P. Although C5 protein itself does not catalyze the cleavage reaction, it facilitates the proper binding to its substrate (65). As expected, C5 addition increased both luminescence (Fig. 5C, “M1 + C5”) and the expected band intensities (Fig. 5D, “RNase P (M1 + C5)”) for all templates.

Next, we evaluated the expression of the highly demanding IPEN tRNA from multicistronic template. To determine the optimal arrangement for transcription and cleavage, we prepared four DNA templates encoding the IPEN tRNA genes in different orders (IPEN, PIEN, EIPN, and NIPE). The translation-coupled assay for these templates showed that higher luminescence was detected for the IPEN (10^6^) and PIEN (10^7^) templates with RNase P (M1 + C5) (Fig. 5E). Consistently, PAGE analysis showed that clear bands were detected at the expected sizes for the IPEN and PIEN templates only in the presence of C5 protein (Fig. 5F). The PIEN template (#5 PIEN, Fig. 5A) was used for subsequent experiments.

### Polycistronic 21 tRNA expression using tRNA array method

To advance toward self-replicating artificial cells, it is ideal for all tRNAs to be encoded in a single genomic DNA. We then constructed a single polycistronic DNA containing all the five tRNA arrays shown in Fig. 5A and evaluated the function of the expressed tRNAs (Fig. 6A). In this experiment, the five transcripts were synthesized from five T7 promoters and underwent a series of processes: self-cleavage of HDVR, dephosphorylation by T4 PNK, and digestion by RNase P to synthesize 21 tRNAs, which are used for luciferase translation. After optimizing NTP and magnesium concentrations (Supplementary Fig. S10), we detected a luciferase activity of 4.7×10^6^, dependent on the single 21 tRNA template (Fig. 6B). This luminescence value was comparable to those of the 21 IVS tRNAs, which are 7.9×10^5^ under the same condition and 9.1×10^6^ under the conditions used in the above experiments (Fig. 2B, WT). These results demonstrate that a sufficient amount of tRNAs was produced in the PURE system using this method. Consistently, PAGE analysis of the reaction product revealed that the total amount of synthesized tRNA (0.82 µg/µl) was similar level to the amount of 21 IVS tRNA (0.6 µg/µl) usually included in the PURE system (49)(Fig. 6C). We further investigated the composition of the synthesized tRNAs by direct RNA sequencing using an Oxford Nanopore sequencer according to a previous study (67, 68). In this method, adapters were attached to RNAs with CCA termini, and direct RNA sequencing revealed the relative abundance of tRNAs of the correct size. We found that although the ratio of each tRNA varied, all 21 tRNAs were synthesized at least 1% of the total tRNA (corresponding to 7.9 ng/µl).

**Fig. 6.**
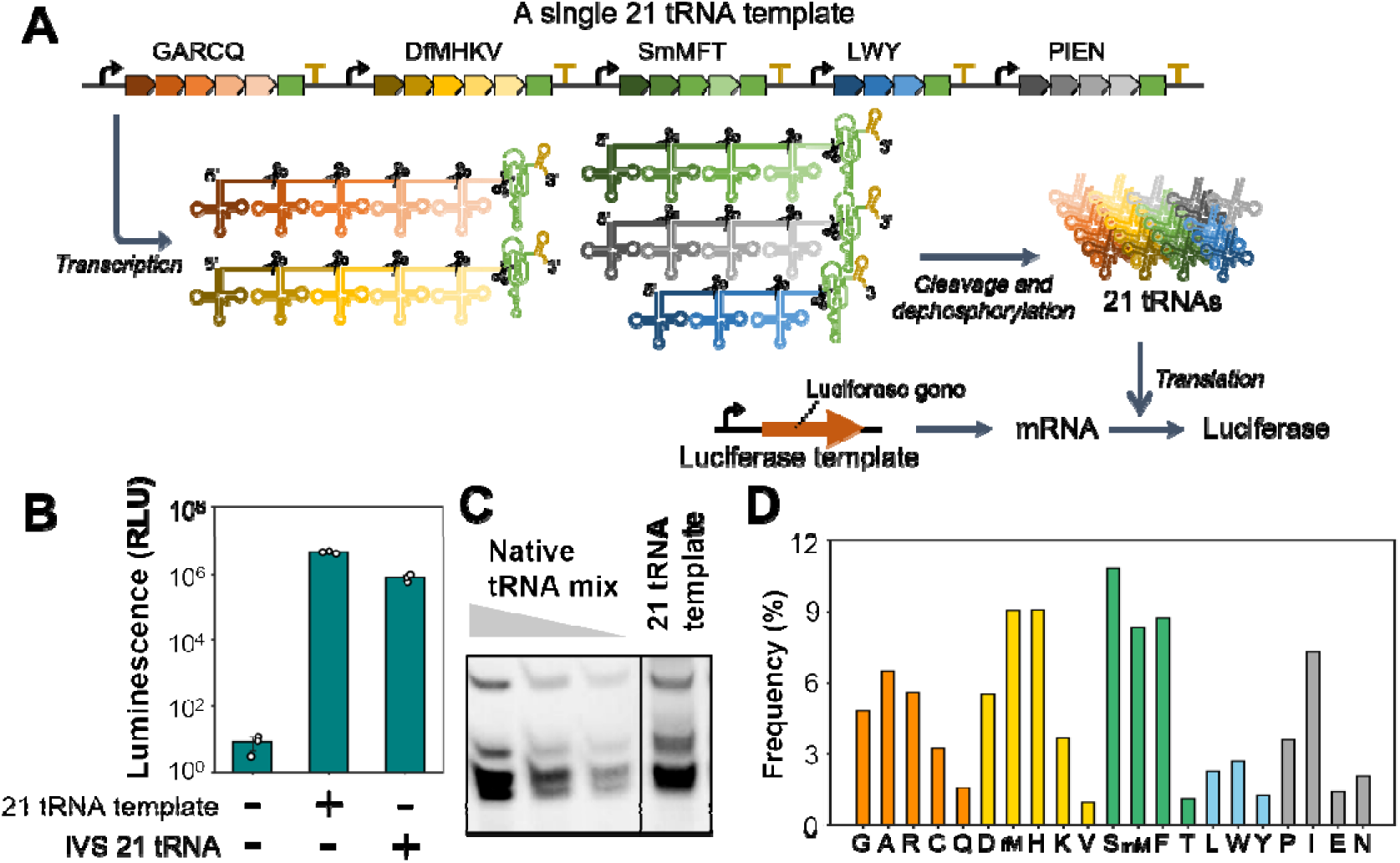
Simultaneous expression of 21 tRNAs from a single polycistronic DNA template. (A) Scheme of the simultaneous expression of 21 tRNAs coupled with translation using the tRNA array method. All tRNA arrays shown in Fig. 5A were encoded on a single DNA template, which was transcribed to produce five transcripts. The transcripts are processed through digestion by RNase P and self-cleavage of HDVR, followed by dephosphorylation by T4 PNK to produce 21 tRNAs, which are used for luciferase translation. (B) Luciferase activity after the translation-coupled reaction. The reaction mixture contained the single 21 tRNAs template (50 nM), luciferase template (1 nM), T7 RNA polymerase (3.4 U/μL), T4 PNK (0.094 U/μL), M1 RNA (4 μM), C5 protein (6 μM), and the tfPURE system (composition E). The mixture was incubated at 30 °C for 24 h and luminescence was measured. As a control, a mixture of purified 21 IVS tRNAs was used instead of the tRNA template. Each data point represents three independent experiments, and error bars indicate standard deviations. (C) PAGE analysis of the RNA product after reaction is shown in (B). A mixture of native E. coli tRNAs (Roche) was used as the control for quantification. (D) Frequency of each tRNA after the reaction. The RNA product after the reaction with the single 21 tRNA template conducted in (B) was analyzed using direct RNA sequencing.

We further tested whether additional tRNAs could be included in a single array. First, we constructed four arrays consisting of 8-9 tRNA genes and evaluated the luciferase activity in the translation-coupled reaction (Supplementary Fig. S11A). All the four constructs exhibited similar luminescence at approximately 10^6^. We then chose two constructs (GL and DS) and integrated them into a single array consisting of 17 tRNA genes of two different orders. Luciferase activity of the

GLDS construct was comparable to that of the negative control without tRNA template, whereas the DSGL construct exhibited luciferase activity at a level of approximately 10^5^ (Supplementary Fig. S11B). When an additional PIEN unit was appended to the 3′ end of the DSGL to construct an all 21-tRNA array, luciferase activity was higher than that without the tRNA template, but the level was significantly low (3.6 × 10^2^) (Supplementary Fig. S11C). These results suggest that while the integration of 21 tRNAs into a single operon is feasible, the translation activity is much lower than that of the separated operons used in Fig. 6.

### Quality check of the luciferase produced by the tRNA array method

The tRNA mixture synthesized using the tRNA array method lacks any modifications and has a different tRNA composition from the 21 IVS tRNAs used in previous studies (42, 49). These differences may reduce the fidelity of translation and, consequently, lower the activity of the synthesized protein. To investigate this, we compared the activity per molecule of luciferase translated by three different tRNA mixtures: native tRNAs purified from *E. coli*, 21 *in vitro* synthesized (IVS) tRNAs, and 21 tRNAs expressed by the tRNA array method. Protein concentrations of translated luciferases were quantified by western blotting (Supplementary Fig. S12A), and their activities were assessed by measuring their luminescence. Luciferase activity per molecule (luminescence/fmol) was compared (Supplementary Fig. S12B). The activity per molecule for the 21 tRNAs expressed using the tRNA array method was approximately 46% and 56 % of that of the native *E. coli* tRNAs and 21 IVS tRNAs, respectively, indicating that the current tRNA array method reduced the luciferase activity per molecule by approximately half.

### An application of tRNA array method for genetic code modification

In the experiment described above, we demonstrated the simultaneous expression of all 21 tRNAs using a single polycistronic DNA template in the tfPURE system. If this tRNA array method is used in conventional *in vitro* transcription (IVT) reactions instead of in the PURE system, it can be used as an easy method to prepare the minimum tRNA set, which has been previously prepared by laborious individual IVT (42, 43, 46, 49) or chemical synthesis (45, 49). The minimum tRNA set is useful for genetic code engineering *in vitro*. As a demonstration, we conducted an IVT using the single 21 tRNA template shown in Fig. 6A, T7 RNA polymerase, and T4 PNK at 37°C for 12 h, and then processed with RNase P at 37°C for 12 h to generate the 21 tRNAs (Fig. 7A). PAGE analysis of the reaction mixture after RNase P treatment revealed bands corresponding to the correct tRNAs (Fig. 7B). We obtained 33 µg of total tRNA from 50 µL of the *in vitro* transcription reaction. After purification, tRNA mixture was added to the tfPURE system at 0.6 µg/µl. The translational activity of the 21 tRNA prepared by the tRNA array method (Fig. 7C, Array) was comparable to that for the same total concentration of conventional 21 tRNA prepared individually (Conv), in which each tRNA ratio follows previous studies (42, 49). Note that, in this experiment, we changed the reporter gene from firefly luciferase to Nanoluc (69), a shorter variant of luciferase, for ease of codon rearrangement conducted below.

**Fig. 7.**
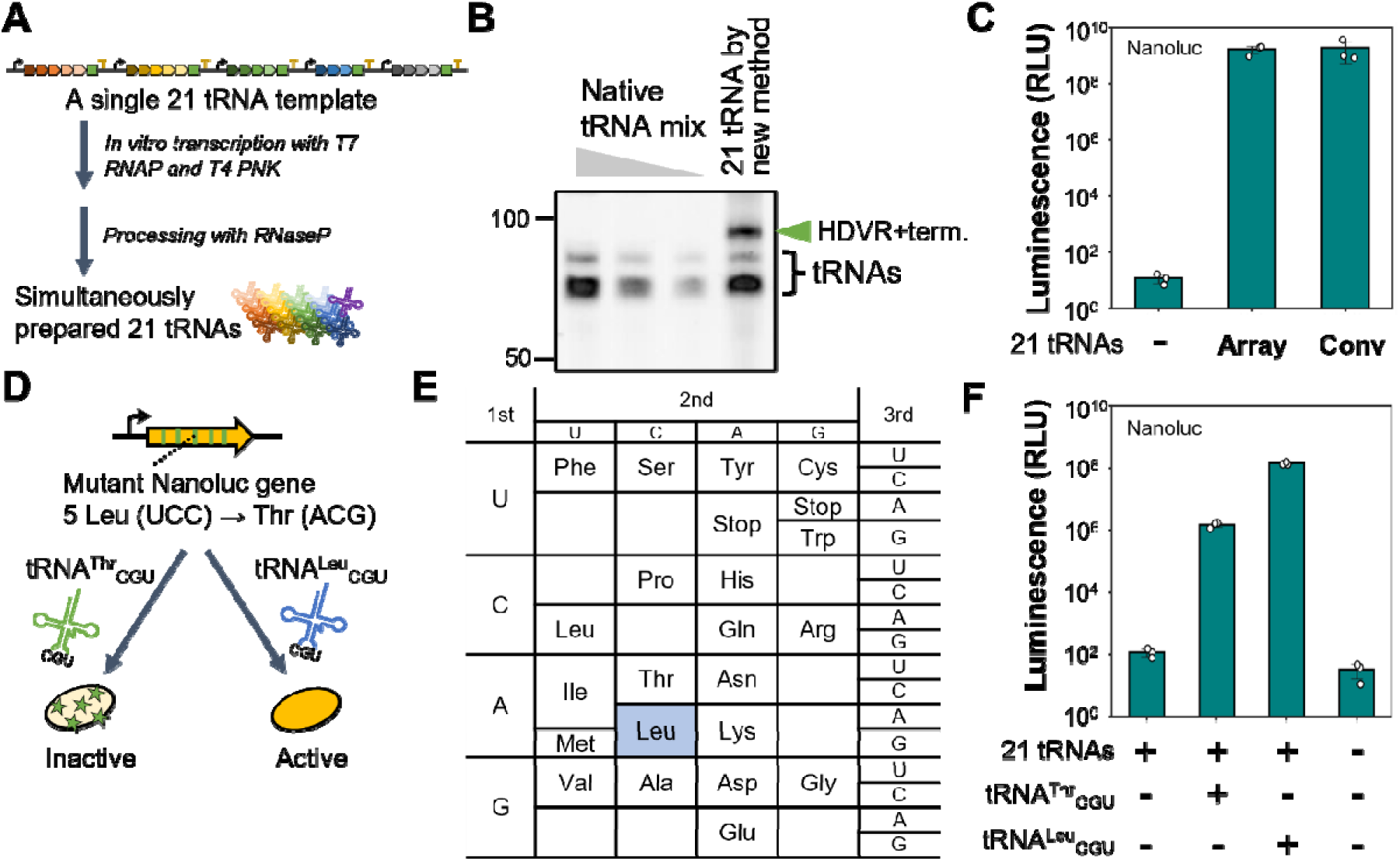
Simultaneous *in vitro* transcription of 21 tRNAs by the tRNA array method and an application for genetic code reassignment. (A) Scheme of the simplified *in vitro* 21 tRNAs synthesis method. For *in vitro* transcription, the single 21 tRNA template (25 nM) was incubated with T7 RNA polymerase (1.0 U/uL) and T4 PNK (0.1 U/uL) at 37°C for 12 h in a buffer containing 2 mM each NTPs (ATP, GTP, CTP, and UTP), 3 mM GMP, Tris-HCl pH 8.0 (40 mM), magnesium chloride (10 mM), spermidine (2 mM), 0.2 U/mL inorganic pyrophosphatase (NEB), and 0.8 U/μL RNasin (Promega). The transcript was purified using a spin column and incubated at 12 μM in buffer R with RNase P (4 μM of M1 RNA and 6 μM of C5 protein) at 37 °C for 16 h. The resulting 21 tRNAs were purified using a spin column before use in the translation assay. (B) PAGE analysis of the tRNA mixture prepared using the method described in (A). E. coli native tRNA mixture (Roche) was used for a control for quantification. The cleaved HDVR with T7 terminator is indicated by a green arrow (HDVR+term). (C) Translation assay using the 21 tRNAs mixture prepared by the tRNA array method (Array) and the conventional individually prepared method (Conv). The reaction mixture containing the 21 tRNAs mixture (600 ng/μL), Nanoluc template (1 nM), T7 RNA polymerase (0.42 U/μL), and the tfPURE system (composition A) was incubated at 30 °C for 16 h, and luminescence was measured. (D) Design of genetic-code modifications. The Nanoluc gene was modified to replace five out of 16 UCC (Leu) codons with ACG codons (Thr in the native codon table). Translation with native tRNA^Thr^_CGU produced a Nanoluc protein with reduced activity due to Thr substitutions at the five Leu residues. In contrast, translation with an anticodon-modified tRNA^Leu^_CGU produced a normal Nanoluc protein, maintaining Nanoluc activity. (E) Reassigned genetic code. In addition to the minimum codon table constructed in a previous study (42), we assigned Leu to the vacant ACG codon (Thr in the native codon table). (F) Translation experiment using the reassigned genetic code. The reaction mixture containing the 21 tRNAs mixture prepared by our method (600 ng/μL), tRNA^Thr^_CGU or tRNA^Leu^_CGU prepared by IVT (12 ng/μL), Nanoluc_ACG template (1 nM), T7 RNA polymerase (0.42 U/μL), and the tfPURE system (composition A). The mixture was incubated at 30 °C for 16 h, and luminescence was measured. Each data point in (C) and (F) represents three independent experiments, and error bars indicate standard deviations.

This method allows for easy customization of the genetic code by introducing additional tRNAs. To demonstrate genetic code reassignment, we replaced five Leu residues (CUU) in Nanoluc with ACG, which is assigned to Thr in the native codon table but is not used in the minimum codon table used here (Fig. 7D). This reassignment has not been used in previous genetic code-engineering studies (42, 43, 46, 47). If this mutant Nanoluc is used, the active Nanoluc should be translated only in the presence of an engineered tRNA (tRNA^Leu^_CGU) that assigns Leu to the ACG codon. To validate this hypothesis, we performed translation reactions of the mutant Nanoluc in the tfPURE system containing the 21 tRNA set prepared by the tRNA array method, and either tRNA^Thr^_CGU (native anticodon) or tRNA^Leu_^CGU (engineered anticodon). As expected, high Nanoluc activity (>10^8^) was detected when engineered tRNA^Leu^_CGU was present, whereas activity decreased by approximately 1/100 when normal tRNA^Thr^_CGU was present. These results demonstrate the usefulness of the minimal 21 tRNAs set for genetic code reassignment.

## Discussion

A major hurdle in constructing a tRNA synthesis system for self-reproducing artificial cells is the processing of the 5’ and 3’ ends of tRNA, which is achieved through complicated multistep processes in the cell (48). In this study, we developed simpler processing methods that work in the PURE system. For the 5’-end, we tested two methods for six tRNA species that could not be directly synthesized from the T7 promoter: the leader method (Fig. 1) and 5’-G variant method (Fig. 2). Using these methods, we demonstrated that all six non-G-start tRNAs could be simultaneously expressed in the PURE system from six monocistronic templates (Fig. 3). For the 3’-end, we developed two methods: the HDVR and linked tRNA methods (Fig. 4). By combining these two methods (termed the tRNA array method), all 21 tRNAs were simultaneously expressed from a single polycistronic DNA template (Fig. 6). The translation ability of the 21 tRNA produced was comparable to the conventional 21 IVS tRNAs that were prepared and purified individually (Fig. 6). These results demonstrate that all minimum tRNA sets can be simultaneously synthesized from multicistronic or single polycistronic DNA template in the PURE system and used for translation, providing a step toward self-reproducing artificial cells.

The tRNA array method established in this study also provides an easy means for the simultaneous *in vitro* synthesis of multiple tRNAs in a conventional IVT buffer, instead of the PURE system (Fig.7). *In vitro*-synthesized tRNA have been utilized for several applications, such as peptide drug screening through the incorporation of non-canonical amino acids (43, 70, 71) and biological containment by constructing a swapped genetic code (42, 46, 47). To date, each tRNA has been individually synthesized through *in vitro* transcription or chemical synthesis, which requires significant effort, time, and cost. Using the method developed in this study, all 21 tRNAs were synthesized from a single DNA template through IVT in the presence of T4 PNK, followed by RNase P processing (Fig. 7A). As shown in Figs. 7D, E, and F, this system also allows the incorporation of additional tRNA to customize the genetic code. The tRNA array method allows easier preparation of the minimum tRNA set and facilitates further development in the field of genetic code reprogramming (72).

One of the remaining challenges in the tRNA array method is the regulation of expression levels of individual tRNAs. The required amounts of each tRNA are expected to vary depending on the protein to be translated, but the current expression level of each tRNA is unregulated. Composition analysis using nanopore sequencing revealed that the expression levels of tRNAs depended on their position in the arrays (Fig. 6D). In particular, central tRNA (e.g., A, fM, H, W, and I) in each array tended to be expressed more. This may have been caused by unexpected transcriptional termination in the middle of the tRNA array. Exploring and utilizing RNA polymerases with higher processivity may address this issue. In addition, given that all tRNAs in the LWY group exhibited lower expression, the composition of the array could affect the expression level. Rearrangement of the order of tRNAs within an array may improve total expression. The importance of the order of the array is also supported by Fig. 5D, in which the expression levels varied significantly depending on the arrangement of IPEN. These differences could be due to the different secondary structures caused by the different orders. The structure can also affect RNase P recognition (73). Understanding the relationship between the linked tRNA structure, transcriptional efficiency, and cleavage efficiency is important for the regulation and improvement of tRNA expression.

In this study, we demonstrated that when tRNAs were directly linked, both 5’- and 3’-end processing were achieved with RNase P. In contrast, extant living systems use various multistep processing for the 3’-end. For example, in *E. coli*, the 3’-end of tRNAs is processed by RNase E, leaving several nucleotides that are further trimmed by multiple exonucleases (48). In eukaryotes, tRNA genes do not generally encode CCA at the 3’-end; instead, extra sequences at the 3’-end are removed by tRNase, and CCA residues are subsequently added by nucleotidyltransferase (74, 75). This complexity in nature raises a fundamental question: why do natural organisms use such intricate processing systems, even though correct processing can be done by only RNase P, at least *in vitro*? One possible explanation is its role in ensuring regulation and quality control (76). Sequential processing steps may act as checkpoints to exclude improperly folded or defective pre-tRNAs, thereby maintaining translational fidelity. On the other hand, a processing scheme using self-cleaving ribozyme to 3’-end of a tRNA, as utilized in this study, has recently been found in many viruses (77), which suggest that such a simpler processing mechanism may also work in nature. Further investigation into the differences between our simplified processing and natural processing would provide useful information for understanding the biological significance of natural processing systems.

The 21 tRNA synthesized by the tRNA array method enabled approximately 3.7 µg/ml translation, comparable to that using 21 IVS tRNAs; however, it remained significantly lower than that with E. coli native tRNAs (56 µg/ml). The translation ability should be improved to realize self-reproducible artificial cells (32, 36). The limitation in translation ability is likely caused by a lack of modification, which reduces aminoacylation efficiency and translational fidelity (42, 78), particularly for tRNAs, such as tRNA^IPEN^. One possible strategy for overcoming this limitation is *in vitro* evolution. The tRNA array method established in this study can be used for a circular DNA template. If tRNA is encoded in a circular DNA template and the translation of phi29 DNA polymerase is coupled, the tRNA sequence can be subjected to *in vitro* evolution using our previous rolling-circle replication scheme (21). Through *in vitro* evolution, tRNA sequences that exhibit a higher translation ability, even in the absence of modifications, would be selected.

## Methods

### Reporter DNA preparation

The luciferase template DNA was prepared by PCR amplification using primers 1 and 2 (Supplementary Tables 1 and 2) and a plasmid (pUC-T7p-Fluc_21tRNA) constructed in our previous study (49). DNA fragments encoding Nanoluc sequences used in Fig. 7 (WT and genetic code reprogrammed version) were obtained using Twist Bioscience’s artificial gene synthesis service. The sequences are listed in Supplementary Table 3. DNA templates for the translation experiments in the PURE system were prepared by PCR amplification using primers 1 and 2. DNA concentrations were quantified based on the A_260_.

### tRNA and M1 RNA preparation

The sequences of the 21 tRNA were based on a previous study (42). The 21 IVS tRNAs were prepared as previous described (49). The 15 IVS tRNAs that has G in the 5’ end (Ala, Arg, Asp, Cys, Gly, Glu, His, Leu, Lys, mMet, Phe, Ser, Thr, Tyr Val) and 5’-G or 5’-A variants shown in Fig. 2A were prepared by *in vitro* transcription utilizing tRNA template prepared by PCR using primer sets shown Supplementary Tables S1 and S2. Six non-G-start tRNAs were chemically synthesized (Invitrogen). The reaction mixture for *in vitro* transcription included 5 mM DTT, 1 U/μl T7 RNA polymerase (Takara, Japan), 2 mM each NTPs(ATP, GTP, CTP, and UTP), 3 mM GMP, 40 mM Tris-HCl pH 8.0, 10 mM magnesium chloride, 2 mM spermidine, 5 μg/μl template DNA, 2 U/ml inorganic pyrophosphatase (New England Biolab), 0.8 U/μl RNasin (Promega). The mixture was incubated at 37°C for 12 h and the RNA product was purified using the PureLink RNA Mini Kit (Invitrogen). Both tRNAs and M1 RNA were dissolved in water and stored at -80 °C until use. RNA concentrations were determined based on the A260. The template DNAs for *in vitro* transcription were prepared by PCR using a plasmid encoding each tRNAs as a template and the primers shown in Supplementary Table S1. Each reverse primer contained a 2’-O-methylation at the second nucleotide from the 5’ terminus to prevent nucleotide addition by T7 RNA polymerase. The template DNA for M1 RNA was prepared by PCR using a plasmid encoding M1 RNA (79)(Supplementary Table S3) as a template, and primers 53 and 54.

To prepare tRNA^Thr^ with a CGU anticodon, plasmid DNA containing tRNA^Thr^ GGU was mutated at the anticodon region by PCR using primers 61 and 62, followed by cloning in *E. coli*. Similarly, tRNA^Leu^ variants were prepared by substituting only the anticodon of tRNA^Leu^CAG with primers 63 and 64. The resultant plasmids were used as PCR templates to prepare the template DNA for *in vitro* transcription.

### tRNA template preparation for expression in PURE system

The tRNA templates used in Fig. 1 (those attached to a leader sequence) were prepared by PCR using the plasmids encoding each tRNA (42) and the primer sets listed in Supplementary Table S1, 2. The tRNA templates used in Fig. 2 (5’-G or-A variants) were the same as those used for *in vitro* transcription, as described above. The tRNAs templates used in Fig. 4 (those attached to the HDVR and linked tRNAs) were prepared as follows. Plasmids encoding each sequence (tRNA^Ala-Ser^, tRNA^Ala-Ser^-HDVR, and tRNA^Ala-Ser^-HDVR-T7hyb10) were constructed using PCR to prepare the vector and insert sequences, followed by ligation by using the In-Fusion cloning kit (Takara, Japan). The tRNAs templates were prepared by PCR using primers 3 and 56 and each of the plasmids. The tRNA templates used in the tRNA array method were prepared as follows. Each plasmid encoding GARCQ, DfMHKV, SmMFT, LWY, or IPEN was obtained from Twist Bioscience’s Artificial Gene Synthesis Service (ex. pTwist_tRNAs^GARCQ^). The plasmids used for the larger tRNA arrays are shown in Supplementary Fig. S11 were constructed sequentially. The plasmid encoding all 21 tRNAs used in Fig. 6 was constructed by PCR amplification of each fragment and assembled using the In-Fusion cloning kit (Takara, Japan). The single 21 tRNA template was prepared by PCR using primer 3 and 56 and each of the plasmid. All tRNA sequences used in this study are listed in Supplementary Table 4.

### Preparation of tfPURE system and C5 protein

All components of the lab-made PURE system were prepared using affinity chromatography with a histidine tag, followed by gel-filtration chromatography based on a previous study (80). EF-Tu was further purified by two times of affinity chromatography to remove residual tRNA as previously reported (49). The detailed purification process is described in the Supplementary Methods.

Ribosome was further purified by using size-separation spin column as follows. Ribosomes purified by the previous method were diluted to a concentration of 200 nM using Buffer D (10 mM Tris-OAc, pH 7.5, 1 mM Mg(OAc)_2_, 60 mM NH_4_Cl, 0.5 mM EDTA, 6 mM 2-mercaptoethanol), which had the same composition as the buffer used for subunit separation (81). Centrifugation was performed using a 100 kDa cut-off Amicon Ultra centrifugal filter until the ribosome concentration reached approximately 1 μM. The sample was then diluted again to approximately 200 nM with Buffer D, followed by repeated centrifugation. This cycle was repeated 20 times. Subsequently, buffer exchange was performed by diluting the sample five-fold in ribosome stock buffer (20 mM HEPES-KOH, pH 7.6, 6 mM Mg(OAc)_2_, 30 mM KCl, and 7 mM 2-mercaptoethanol) and centrifuging once, followed by two consecutive 50-fold dilutions and centrifugation. The tfPURE system was prepared using the further purified EF-Tu and ribosome with other proteins purified by the conventional method. The composition is shown in Supplementary Table S5 and S6.

C5 protein was purified from E. coli harvoring C5 expressing plasmid, pET-C5 (79), using affinity chromatography for the histidine tag by the same method as the component of the PURE system as described above.

### PAGE analysis

Electrophoresis was carried out using 8% (acrylamide:bis = 19:1) polyacrylamide gel containing 8 M urea, 0.1% ammonium persulfate, and 0.1% N, N, N’, N’-tetramethylethylene-diamine in Tris– borate EDTA buffer. The samples were prepared by mixing with stripping buffer composed of 50 mM EDTA, 90% formamide, and 0.025% bromophenol blue. RNA was stained with SYBR Green II (Takara, Japan).

### Translation assays with the 21 IVS tRNAs conducted in Fig. 2B

The reaction mixture containing the 21 IVS tRNAs, luciferase DNA (1 nM), T7 RNA polymerase (0.42 U/µL, Takara, Japan), NTP (0.88 mM), magnesium acetate (7.9 mM), and the laboratory-made tfPURE system was incubated at 30°C for 16 h. The total concentration of the IVS tRNAs was 600 ng/μL (100 ng/μL each for IPEN and 11.7 ng/μL each for the others), as previously reported (49). For assays involving tRNA variants, PURE systems lacking the respective tRNA were prepared, and each tRNA variant was added at the same concentration as the WT tRNA in the IVS tRNA mixture. After the incubation, an aliquot (1 μL) of the reaction mixture was added to 30 μL of Luciferase assay reagent (Promega), and luminescence was measured with GloMax Luminometer (Promega).

### tRNA expression-coupled translation in the PURE system

The reaction mixture containing luciferase or Nanoluc DNA (1 nM), tRNA template DNA, T7 RNA polymerase (Takara, Japan), T4 PNK (New England Biolabs), RNase P (M1 RNA and C5 protein), and the tfPURE system was incubated at 30°C. The concentrations of the tRNA template DNA, T7 RNA polymerase, T4 PNK, and RNase P varied depending on the experiment (see each legend). The reaction time was set to 16 h or 24 h, depending on the experiment (see each legend). For each tRNA expression scheme, the concentrations of NTPs, Mg²L, spermidine, and T7 RNA polymerase were optimized (Supplementary Table S6). After incubation, an aliquot (1 μL) of the reaction mixture was added to 30 μL of Luciferase assay reagent (Promega) for luciferase or Nanoluc assay reagent (Promega) for Nanoluc, and luminescence was measured using GloMax Luminometer (Promega).

### Western blotting

After incubation, the reaction mixture (1 μL) was mixed with SDS buffer (17 mM Tris-HCl, pH 7.4, 0.7% SDS, 0.3 M 2-mercaptoethanol, and 3% glycerol), boiled at 95°C for 5 min, and subjected to 10% SDS-PAGE. Protein transfer was performed using the WSE-4115 system (Atto, Japan), followed by blocking with a primary antibody (Host/Isotyle: Rabbit/IgG, anti-luciferase antibody; PGI Proteintech Group. Inc., 27986-1-AP), and secondary antibodies (Host: Goat anti-rabbit IgG, PGI Proteintech Group. Inc., SA00001-2) using an iBind Western System (Thermo Fisher). Pierce ECL Western Blotting Substrate (Thermo Fisher) was used as a light-emitting chemiluminescent substrate, incubated at room temperature, and detected using a FUSION-SL4 imaging system (Vilber-Lourmat). A dilution series of recombinant luciferase (L9420; Sigma-Aldrich) was prepared and used as a standard for quantification.

### Nanopore tRNA direct sequencing

After the tRNA expression-coupled translation reaction in the PURE system, the total RNA was purified using the PureLink RNA Mini Kit (Invitrogen). To remove the 5’-end triphosphate from specific tRNAs (Gly, Asp, Ser, Leu, and Pro), RppH was added to NEBuffer 2, followed by incubation at 37°C for 30 min and 65°C for 5 min. The RNA was purified using AMPure RNAClean XP. Subsequently, direct tRNA sequencing was performed as previously reported (67, 68) using RNA Direct Sequencing Kit (Oxford Nanopore). Purified RNAs were incubated with 5’ (5’ P-GGCUUCUUCUUGCUCUUAGGAAAAAAAAAAaaa-3’, where lowercase “a” denotes DNA) and 3’ (5’-CCUAAGAGCAAGAAGAAGCCUGGN-3’, where “N” represents A, U, G, or C) splint adapters for the first ligation. The RNA was purified again using AMPure RNAClean XP. RNA was then ligated to the RTA adapter. The RNA was linearized by reverse transcription using Maxima H Minus Reverse Transcriptase (Life Technologies, EP0751). Linearized tRNAs were cleaned using 2× AMPure RNAClean XP beads. Finally, ONT RMX sequencing adapters were ligated and applied to a MinION flow cell (FLO-MIN-106) using the standard ONT SQK-RNA002 protocol. The reads were mapped using minimap2 version 2.24-r1122 with sensitive parameters (-ax map-on -k10).

### Simultaneous 21 tRNAs synthesis for codon reprograming

*In vitro* transcription was carried out with the single 21 tRNA template (25 nM) in the presence of T7 RNA polymerase (1.0 U/ul) and T4 PNK (0.1 U/μL, New England Biolab), NTPs (2 mM each: ATP, GTP, CTP, and UTP), GMP (3 mM), Tris-HCl pH 8.0 (40 mM), magnesium chloride (10 mM), spermidine (2 mM), 0.2 U/ml inorganic pyrophosphatase (New England Biolab), and 0.8 U/μl RNasin (Promega) at 37°C for 12 h. The premature tRNA product was purified using PureLink RNA Mini Kit (Invitrogen). The purified premature tRNA was incubated in buffer R (50 mM Tris HCl(pH7.6), 60 mM NH_4_Cl, 10 mM Mg(OAc)_2_, and 5 mM spermidine) with RNase P (4 μM M1 RNA and 6 μM C5 protein) at 37 °C for 12 h for digestion. The mature tRNA product was purified using PureLink RNA Mini Kit.

## Supplementary Materials

Supplementary material for this article is available at XXX

## Acknowledgements

We thank Mrs. Kayo Aoyama and Ayu Saito for technical support.

## Author contributions

R.M., Y. S., N.I., and Y.S. designed the study and wrote the manuscript. R. M., and N.I. performed experiments and analyses.

## Data and materials availability

Data is available on request from the authors.

## Reference

1. Buddingh’, B.C. and Van Hest, J.C.M. (2017) Artificial Cells: Synthetic Compartments with Life-like Functionality and Adaptivity. Acc Chem Res, 50.

2. Ivanov, I., Castellanos, S.L., Balasbas, S., Otrin, L., Maruscaroniccaron, N., Vidakovicacute-Koch, T. and Sundmacher, K. (2021) Bottom-Up Synthesis of Artificial Cells: Recent Highlights and Future Challenges. 12.

3. Olivi, L., Berger, M., Creyghton, R.N.P., De Franceschi, N., Dekker, C., Mulder, B.M., Claassens, N.J., ten Wolde, P.R. and van der Oost, J. (2021) Towards a synthetic cell cycle. Nat Commun, 12.

4. Biner, O., Fedor, J.G., Yin, Z. and Hirst, J. (2020) Bottom-Up Construction of a Minimal System for Cellular Respiration and Energy Regeneration. ACS Synth Biol, 9.

5. Libicher, K., Hornberger, R., Heymann, M. and Mutschler, H. (2020) In vitro self-replication and multicistronic expression of large synthetic genomes. Nat Commun, 11.

6. Libicher, K. and Mutschler, H. (2020) Probing self-regeneration of essential protein factors required for: In vitro translation activity by serial transfer. Chemical Communications, 56.

7. Doerr, A., Foschepoth, D., Forster, A.C. and Danelon, C. (2021) In vitro synthesis of 32 translation-factor proteins from a single template reveals impaired ribosomal processivity. Sci Rep, 11.

8. Blanken, D., Foschepoth, D., Serrão, A.C. and Danelon, C. (2020) Genetically controlled membrane synthesis in liposomes. Nat Commun, 11.

9. Wang, C., Yang, J. and Lu, Y. (2021) Modularize and Unite: Toward Creating a Functional Artificial Cell. Front Mol Biosci, 8.

10. Liu, Y., Davis, R.G., Thomas, P.M., Kelleher, N.L. and Jewett, M.C. (2021) In vitro-Constructed Ribosomes Enable Multi-site Incorporation of Noncanonical Amino Acids into Proteins. Biochemistry, 10.1021/acs.biochem.0c00829.

11. Vibhute, M.A., Schaap, M.H., Maas, R.J.M., Nelissen, F.H.T., Spruijt, E., Heus, H.A., Hansen, M.M.K. and Huck, W.T.S. (2020) Transcription and Translation in Cytomimetic Protocells Perform Most Efficiently at Distinct Macromolecular Crowding Conditions. ACS Synth Biol, 9.

12. Amy Yewdall, N., Mason, A.F. and Van Hest, J.C.M. (2018) The hallmarks of living systems: Towards creating artificial cells. Interface Focus, 8.

13. Tror, S., Jeon, S.M., Nguyen, H.T., Huh, E. and Shin, K. (2023) A Self-Regenerating Artificial Cell, that is One Step Closer to Living Cells: Challenges and Perspectives. Small Methods, 7, 2300182.

14. Kosaka, Y., Miyawaki, Y., Mori, M., Aburaya, S., Nishizawa, C., Chujo, T., Niwa, T., Miyazaki, T., Sugita, T., Fukuyama, M., et al. (2025) Autonomous ribosome biogenesis in vitro. Nature Communications 2025 16:1, 16, 1–14.

15. Ichihashi, N. (2019) What can we learn from the construction of in vitro replication systems? Ann N Y Acad Sci, 1447, 144–156.

16. Nishikawa, S., Yu, W.C., Jia, T.Z., He, M.J., Khusnutdinova, A., Yakunin, A.F., Chiang, Y.R., Fujishima, K. and Wang, P.H. (2024) Amino Acid Self-Regenerating Cell-Free Protein Synthesis System that Feeds on PLA Plastics, CO2, Ammonium, and α-Ketoglutarate. ACS Catalysis, 14, 7696–7706.

17. Fujiwara, K., Katayama, T. and Nomura, S.I.M. (2013) Cooperative working of bacterial chromosome replication proteins generated by a reconstituted protein expression system. Nucleic Acids Res, 41, 7176–7183.

18. Su’etsugu, M., Takada, H., Katayama, T. and Tsujimoto, H. (2017) Exponential propagation of large circular DNA by reconstitution of a chromosome-replication cycle. Nucleic Acids Res, 45, 11525–11534.

19. van Nies, P., Westerlaken, I., Blanken, D., Salas, M., Mencía, M. and Danelon, C. (2018) Self-replication of DNA by its encoded proteins in liposome-based synthetic cells. Nat Commun, 9, 1583.

20. Abil, Z., Restrepo Sierra, A.M., Stan, A.R., Châne, A., del Prado, A., de Vega, M., Rondelez, Y. and Danelon, C. (2024) Darwinian Evolution of Self-Replicating DNA in a Synthetic Protocell. Nature Communications 2024 15:1, 15, 1–15.

21. Sakatani, Y., Yomo, T. and Ichihashi, N. (2018) Self-replication of circular DNA by a self-encoded DNA polymerase through rolling-circle replication and recombination. Sci Rep, 8.

22. Sakatani, Y., Ichihashi, N., Kazuta, Y. and Yomo, T. (2015) A transcription and translation-coupled DNA replication system using rolling-circle replication. Sci Rep, 5.

23. Okauchi, H. and Ichihashi, N. (2021) Continuous Cell-Free Replication and Evolution of Artificial Genomic DNA in a Compartmentalized Gene Expression System. ACS Synth Biol, 10.

24. Grasemann, L., Thiel Pizarro, P. and Maerkl, S.J. (2023) C2CAplus: A One-Pot Isothermal Circle-to-Circle DNA Amplification System. ACS Synth Biol, 12, 3137–3142.

25. Okauchi, H., Sakatani, Y., Otsuka, K. and Ichihashi, N. (2020) Minimization of Elements for Isothermal DNA Replication by an Evolutionary Approach. ACS Synth Biol, 9, 1771–1780.

26. Eto, S., Matsumura, R., Shimane, Y., Fujimi, M., Berhanu, S., Kasama, T. and Kuruma, Y. (2022) Phospholipid synthesis inside phospholipid membrane vesicles. Commun Biol, 5, 1016.

27. Bailoni, E., Patiño-Ruiz, M.F., Stan, A.R., Schuurman-Wolters, G.K., Exterkate, M., Driessen, A.J.M. and Poolman, B. (2024) Synthetic Vesicles for Sustainable Energy Recycling and Delivery of Building Blocks for Lipid Biosynthesis †. ACS Synth Biol, 13, 1549–1561.

28. Sato, G., Kinoshita, S., Yamada, T.G., Arai, S., Kitaguchi, T., Funahashi, A., Doi, N. and Fujiwara, K. (2024) Metabolic Tug-of-War between Glycolysis and Translation Revealed by Biochemical Reconstitution. ACS Synth Biol, 13, 1572–1581.

29. Giaveri, S., Bohra, N., Diehl, C., Yang, H.Y., Ballinger, M., Paczia, N., Glatter, T. and Erb, T.J. (2024) Integrated translation and metabolism in a partially self-synthesizing biochemical network. Science, 385, 174–178.

30. Kohyama, S., Merino-Salomón, A. and Schwille, P. (2022) In vitro assembly, positioning and contraction of a division ring in minimal cells. Nature Communications 2022 13:1, 13, 1–14.

31. Chang, M.Y., Ariyama, H., Huck, W.T.S. and Deng, N.N. (2023) Division in synthetic cells. Chem Soc Rev, 52, 3307–3325.

32. De Capitani, J. and Mutschler, H. (2023) The Long Road to a Synthetic Self-Replicating Central Dogma. Biochemistry, 62, 1221–1232.

33. Shimizu, Y., Inoue, A., Tomari, Y., Suzuki, T., Yokogawa, T., Nishikawa, K. and Ueda, T. (2001) Cell-free translation reconstituted with purified components. Nat Biotechnol, 19.

34. Lavickova, B., Laohakunakorn, N. and Maerkl, S.J. (2020) A partially self-regenerating synthetic cell. Nat Commun, 11.

35. Schwarz-Schilling, M., Dupin, A., Avidan, N., Barak, Y., Shimizu, Y., Daube, S.S. and Bar-Ziv, R.H. (2024) Autonomous biogenesis of the entire protein translation machinery excluding ribosomes. bioRxiv, 10.1101/2024.10.20.619270.

36. Hagino, K., Masuda, K., Shimizu, Y. and Ichihashi, N. (2024) Sustainable Regeneration of 20 Aminoacyl-tRNA Synthetases in a Reconstituted System Toward Self-Synthesizing Artificial Systems. bioRxiv, 10.1101/2024.10.03.616507.

37. Levy, M., Falkovich, R., Daube, S.S. and Bar-Ziv, R.H. (2020) Autonomous synthesis and assembly of a ribosomal subunit on a chip. Sci Adv, 6.

38. Li, J., Haas, W., Jackson, K., Kuru, E., Jewett, M.C., Fan, Z.H., Gygi, S. and Church, G.M. (2017) Cogenerating Synthetic Parts toward a Self-Replicating System. ACS Synth Biol, 6, 1327–1336.

39. Shimojo, M., Amikura, K., Masuda, K., Kanamori, T., Ueda, T. and Shimizu, Y. (2020) In vitro reconstitution of functional small ribosomal subunit assembly for comprehensive analysis of ribosomal elements in E. coli. Commun Biol, 3, 142.

40. Aoyama, R., Masuda, K., Shimojo, M., Kanamori, T., Ueda, T. and Shimizu, Y. (2022) In vitro reconstitution of the Escherichia coli 70S ribosome with a full set of recombinant ribosomal proteins. J Biochem, 171, 227–237.

41. Shepherd, J. and Ibba, M. (2015) Bacterial transfer RNAs. FEMS Microbiol Rev, 39, 280.

42. Hibi, K., Amikura, K., Sugiura, N., Masuda, K., Ohno, S., Yokogawa, T., Ueda, T. and Shimizu, Y. (2020) Reconstituted cell-free protein synthesis using in vitro transcribed tRNAs. Commun Biol, 3.

43. Iwane, Y., Hitomi, A., Murakami, H., Katoh, T., Goto, Y. and Suga, H. (2016) Expanding the amino acid repertoire of ribosomal polypeptide synthesis via the artificial division of codon boxes. Nat Chem, 8, 317–325.

44. Li, J., Tang, M. and Qi, H. (2022) Codon-Reduced Protein Synthesis With Manipulating tRNA Components in Cell-Free System. Front Bioeng Biotechnol, 10, 891808.

45. Calles, J., Justice, I., Brinkley, D., Garcia, A. and Endy, D. (2019) Fail-safe genetic codes designed to intrinsically contain engineered organisms. Nucleic Acids Res, 47.

46. Fujino, T., Tozaki, M. and Murakami, H. (2020) An Amino Acid-Swapped Genetic Code. ACS Synth Biol, 9, 2703–2713.

47. Fujino, T., Sonoda, R., Higashinagata, T., Mishiro-Sato, E., Kano, K. and Murakami, H. (2024) Ser/Leu-swapped cell-free translation system constructed with natural/in vitro transcribed-hybrid tRNA set. Nature Communications 2024 15:1, 15, 1–10.

48. Mohanty, B.K. and Kushner, S.R. (2019) New Insights into the Relationship between tRNA Processing and Polyadenylation in Escherichia coli. Trends in Genetics, 35, 434–445.

49. Miyachi, R., Shimizu, Y. and Ichihashi, N. (2022) Transfer RNA Synthesis-Coupled Translation and DNA Replication in a Reconstituted Transcription/Translation System. ACS Synth Biol, 11, 2791–2799.

50. Su’etsugu, M., Takada, H., Katayama, T. and Tsujimoto, H. (2017) Exponential propagation of large circular DNA by reconstitution of a chromosome-replication cycle. Nucleic Acids Res, 45.

51. Cobaleda, C. and Sánchez-García, I. (2001) RNase P: from biological function to biotechnological applications. Trends Biotechnol, 19, 406–411.

52. Guerrier-Takada, C., Gardiner, K., Marsh, T., Pace, N. and Altman, S. (1983) The RNA moiety of ribonuclease P is the catalytic subunit of the enzyme. Cell, 35, 849–857.

53. Baer, M.F., Reilly, R.M., McCorkle, G.M., Hai, T.Y., Altman, S. and RajBhandary, U.L. (1988) The recognition by RNase P of precursor tRNAs. Journal of Biological Chemistry, 263, 2344–2351.

54. Samhita, L., Virumäe, K., Remme, J. and Varshney, U. (2013) Initiation with Elongator tRNAs. J Bacteriol, 195, 4202.

55. Coleman, T.M., Wang, G. and Huang, F. (2004) Superior 5’ homogeneity of RNA from ATP-initiated transcription under the T7 phi 2.5 promoter. Nucleic Acids Res, 32, e14.

56. Himeno, H., Hasegawa, T., Asahara, H., Tamura, K. and Shimizu, M. (1991) Identity determinants of E. coli tryptophan tRNA. Nucleic Acids Res, 19, 6379.

57. Jahn, M., Rogers, M.J. and Söll, D. (1991) Anticodon and acceptor stem nucleotides in tRNA(Gln) are major recognition elements for E. coli glutaminyl-tRNA synthetase. Nature, 352, 258–260.

58. Nureki, O., Niimi, T., Muramatsu, T., Kanno, H., Kohno, T., Florentz, C., Giegé, R. and Yokoyama, S. (1994) Molecular recognition of the identity-determinant set of isoleucine transfer RNA from Escherichia coli. J Mol Biol, 236, 710–724.

59. Liu, H., Peterson, R., Kessler, J. and Musier-forsyth, K. (1995) Molecular recognition of tRNA(Pro) by Escherichia coli proline tRNA synthetase in vitro. Nucleic Acids Res, 23, 165.

60. Guillon, J.M., Meinnel, T., Mechulam, Y., Lazennec, C., Blanquet, S. and Fayat, G. (1992) Nucleotides of tRNA governing the specificity of Escherichia coli methionyl-tRNAfMet formyltransferase. J Mol Biol, 224, 359–367.

61. Schürer, H., Lang, K., Schuster, J. and Mörl, M. (2002) A universal method to produce in vitro transcripts with homogeneous 3′ ends. Nucleic Acids Res, 30, e56.

62. Calvopina-Chavez, D.G., Gardner, M.A. and Griffitts, J.S. (2022) Engineering efficient termination of bacteriophage T7 RNA polymerase transcription. G3: Genes|Genomes|Genetics, 12, jkac070.

63. Das, U. and Shuman, S. (2012) Mechanism of RNA 2′, 3′-cyclic phosphate end healing by T4 polynucleotide kinase–phosphatase. Nucleic Acids Res, 41, 355.

64. Zhang, J. and Ferré-D’Amaré, A.R. (2016) Trying on tRNA for Size: RNase P and the T-box Riboswitch as Molecular Rulers. Biomolecules 2016, *Vol. 6, Page 18*, 6, 18.

65. Niland, C.N., Anderson, D.R., Jankowsky, E. and Harris, M.E. (2017) The contribution of the C5 protein subunit of Escherichia coli ribonuclease P to specificity for precursor tRNA is modulated by proximal 5′ leader sequences. RNA, 23, 1502–1511.

66. Sun, L., Campbell, F.E., Zahler, N.H. and Harris, M.E. (2006) Evidence that substrate-specific effects of C5 protein lead to uniformity in binding and catalysis by RNase P. EMBO Journal, 25, 3998–4007.

67. Lucas, M.C., Pryszcz, L.P., Medina, R., Milenkovic, I., Camacho, N., Marchand, V., Motorin, Y., Ribas de Pouplana, L. and Novoa, E.M. (2023) Quantitative analysis of tRNA abundance and modifications by nanopore RNA sequencing. Nature Biotechnology 2023 42:1, 42, 72–86.

68. Thomas, N.K., Poodari, V.C., Jain, M., Olsen, H.E., Akeson, M. and Abu-Shumays, R.L. (2021) Direct Nanopore Sequencing of Individual Full Length tRNA Strands. ACS Nano, 15, 16642–16653.

69. Hall, M.P., Unch, J., Binkowski, B.F., Valley, M.P., Butler, B.L., Wood, M.G., Otto, P., Zimmerman, K., Vidugiris, G., MacHleidt, T., et al. (2012) Engineered luciferase reporter from a deep sea shrimp utilizing a novel imidazopyrazinone substrate. ACS Chem Biol, 7, 1848–1857.

70. Ohta, A., Tanada, M., Shinohara, S., Morita, Y., Nakano, K., Yamagishi, Y., Takano, R., Kariyuki, S., Iida, T., Matsuo, A., et al. (2023) Validation of a New Methodology to Create Oral Drugs beyond the Rule of 5 for Intracellular Tough Targets. J Am Chem Soc, 145, 24035–24051.

71. Goto, Y. and Suga, H. (2021) The RaPID Platform for the Discovery of Pseudo-Natural Macrocyclic Peptides. Acc Chem Res, 54, 3604–3617.

72. Costello, A., Peterson, A.A., Chen, P.H., Bagirzadeh, R., Lanster, D.L. and Badran, A.H. (2024) Genetic Code Expansion History and Modern Innovations. Chem Rev, 10.1021/ACS.CHEMREV.4C00275/ASSET/IMAGES/LARGE/CR4C00275_0015.JPEG.

73. Krupp, G., Kahle, D., Vogt, T. and Char, S. (1991) Sequence changes in both flanking sequences of a Pre-tRNA influence the cleavage specificity of RNase P. J Mol Biol, 217, 637–648.

74. Xiong, Y. and Steitz, T.A. (2006) A story with a good ending: tRNA 3′-end maturation by CCA-adding enzymes. Curr Opin Struct Biol, 16, 12–17.

75. Mohan, A., Whyte, S., Wang, X., Nashimoto, M. and Levinger, L. (1999) The 3’ end CCA of mature tRNA is an antideterminant for eukaryotic 3’-tRNase. RNA, 5, 245–256.

76. Mohanty, B.K., Maples, V.F. and Kushner, S.R. (2012) Polyadenylation helps regulate functional tRNA levels in Escherichia coli. Nucleic Acids Res, 40, 4589–4603.

77. Kienbeck, K., Malfertheiner, L., Zelger-Paulus, S., Johannsen, S., von Mering, C. and Sigel, R.K.O. (2024) Identification of HDV-like theta ribozymes involved in tRNA-based recoding of gut bacteriophages. Nature Communications 2024 15:1, 15, 1–10.

78. Cui, Z., Stein, V., Tnimov, Z., Mureev, S. and Alexandrov, K. (2015) Semisynthetic tRNA complement mediates in vitro protein synthesis. J Am Chem Soc, 137, 4404–4413.

79. Fukunaga, J.I., Gouda, M., Umeda, K., Ohno, S., Yokogawa, T. and Nishikawa, K. (2006) Use of RNase P for efficient preparation of yeast tRNATyr transcript and its mutants. J Biochem, 139, 123–127.

80. Fujii, S., Matsuura, T., Sunami, T., Nishikawa, T., Kazuta, Y. and Yomo, T. (2014) Liposome display for in vitro selection and evolution of membrane proteins. Nature Protocols 2014 9:7, 9, 1578–1591.

81. Jewett, M.C., Fritz, B.R., Timmerman, L.E. and Church, G.M. (2013) In vitro integration of ribosomal RNA synthesis, ribosome assembly, and translation. Mol Syst Biol, 9.

