## Supplemental information for "Simultaneous in vitro expression of minimal 21 transfer RNAs by tRNA array method"

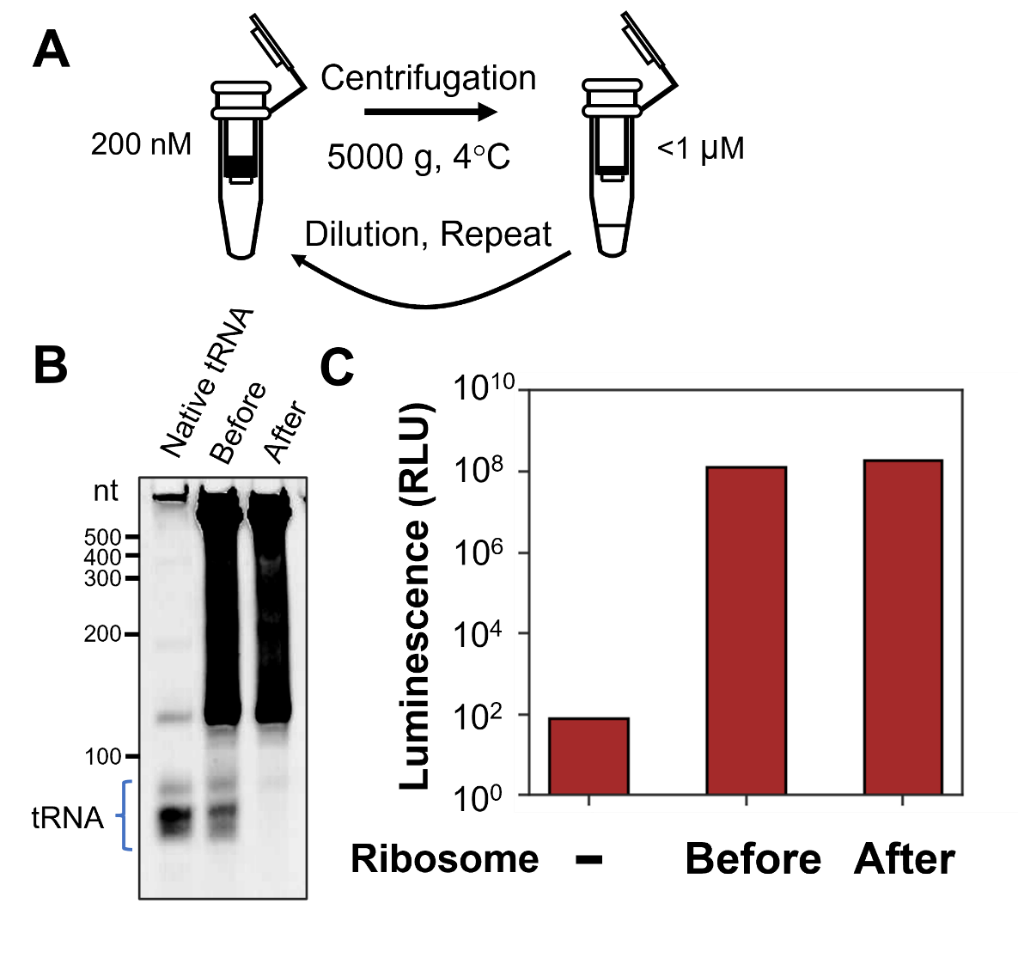

**Supplementary Figure S1. Removal of tRNA from ribosomes with size-separation**

(A) Procedure for tRNA removal using a size separation spin column. Ribosomes were diluted to 200 nM in Buffer D and concentrated to 1 μM using a size-separation spin column (Millipore, Amicon Ultra Centrifugal Filter, 100 kDa MWCO). This cycle of dilution and concentration was repeated approximately 20 times until tRNA was removed. (B) PAGE analysis of ribosomes before and after tRNA removal. A total of 1 pmol of ribosomes before and after tRNA removal were subjected to PAGE analysis. *E. coli* native tRNA (Roche) was applied for comparison. (C) The translation activity of ribosomes before and after tRNA removal. PURE systems lacking ribosomes were prepared, and the translation activity was evaluated by adding ribosomes before or after tRNA removal. Luciferase was used as a reporter, and the reaction mixture contained a luciferase template (1 nM), T7 RNA polymerase (0.42 U/μL), and *an E. coli* native tRNA mixture (Roche). The samples were incubated at 30 °C for 16 h, and luciferase activity was measured.

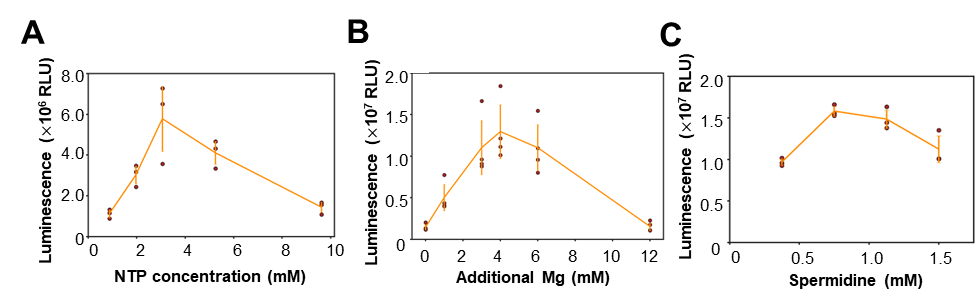

**Supplementary Figure S2. Optimization of PURE system composition for tRNA^Gln^ expression using the leader method**

(A) Optimization of the NTP concentration. The concentration of NTPs was optimized by incrementally adding NTPs and equimolar concentrations of Mg(OAc)_2_. The optimal composition was determined to be 3.1 mM NTPs. (B) Optimization of the Mg(OAc)_2_ concentration. After fixing the NTP concentration at the optimal 3.1 mM, the additional Mg(OAc)_2_ concentration was varied and determined to be 4 mM. (C) Optimization of spermidine. The additional Mg(OAc)_2_ concentration was fixed at 3 mM, and the additional spermidine was varied and determined to be 0.75 μM. The final optimized compositions are summarized in Supplementary Table 6 (Composition B).

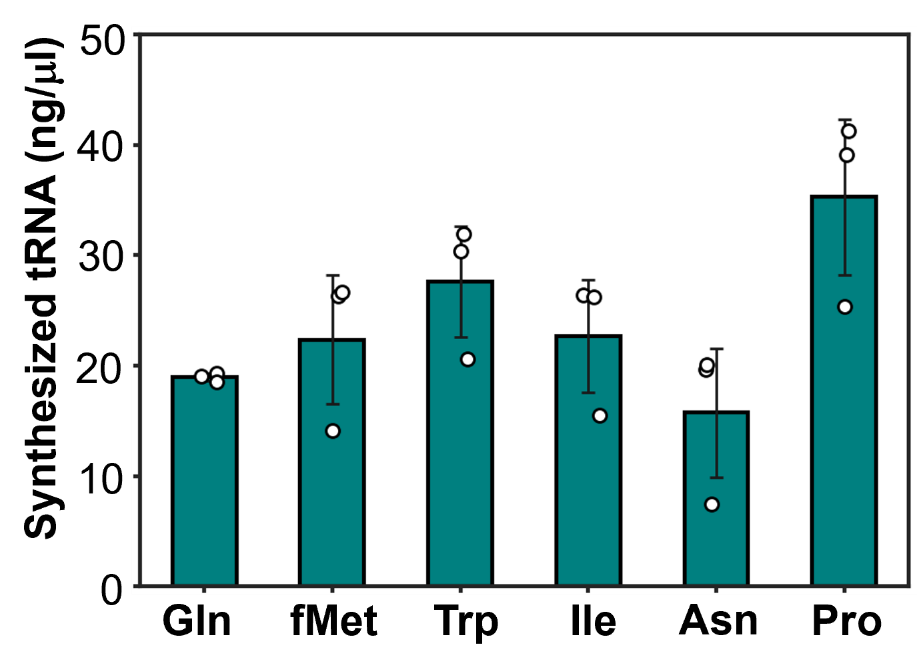

**Supplementary Figure S3. Quantification of tRNA concentrations synthesized by the leader method**

The tRNA products shown in Fig. 1E and 1F were quantified using ImageJ software. Quantification was based on comparison with control tRNAs of known concentrations. Each data point represents the result of three independent experiments, with error bars indicating standard deviations.

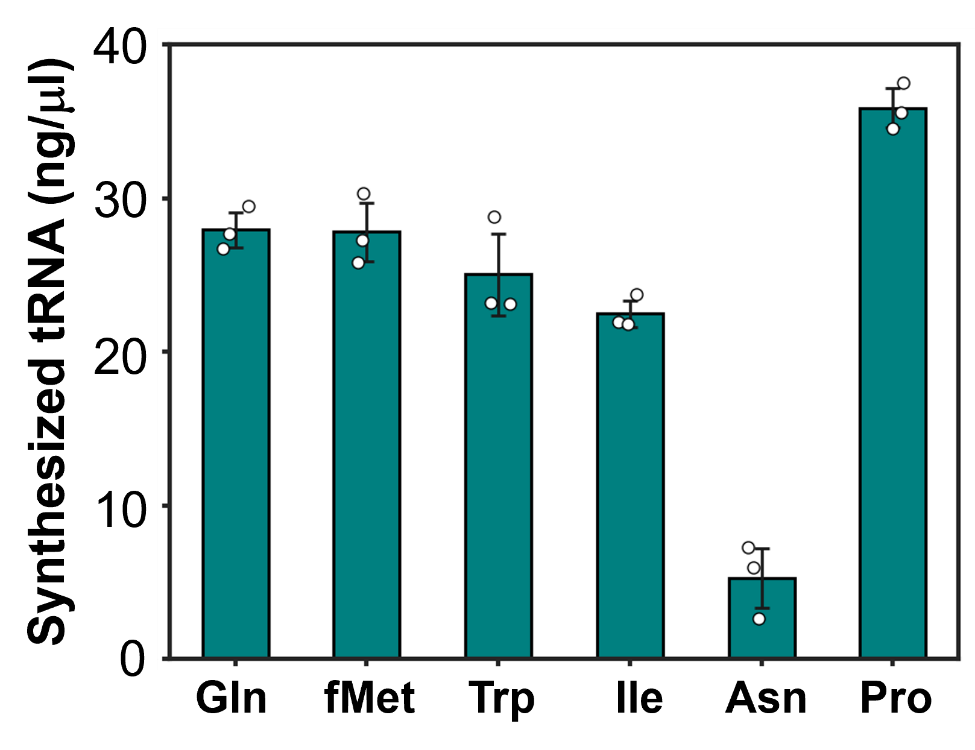

**Supplementary Figure S4. Quantification of tRNA concentrations synthesized by the 5’-G variant method**

The tRNA products shown in Fig. 2E were quantified using ImageJ software. Quantification was based on comparison with control tRNAs of known concentrations. Each data point represents the result of three independent experiments, with error bars indicating standard deviations.

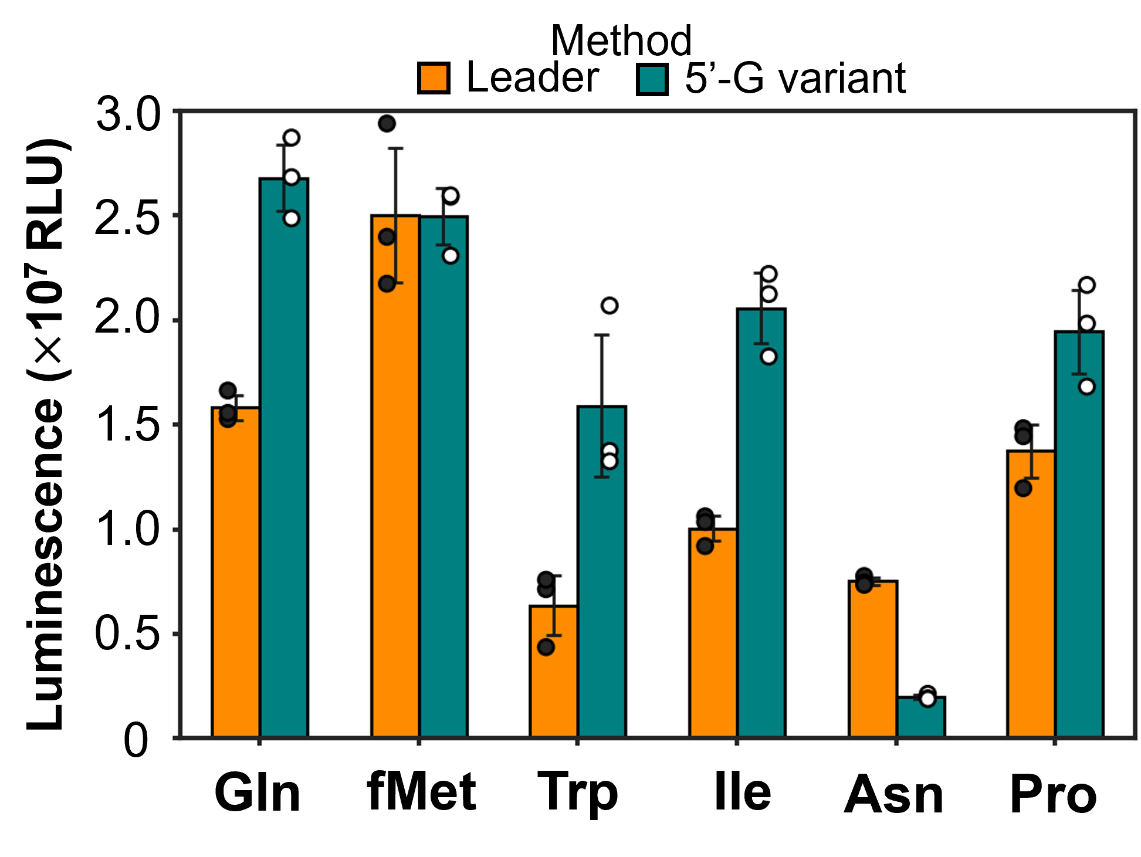

**Supplementary Figure S5. Comparison of the translation activity of tRNA prepared by each 5’-end preparation method.**

The 5'-G variant method showed a higher translation activity than the leader method for tRNA^Gln, Ile Pro, and Trp^. For tRNA^Asn^, the leader method outperformed the 5'-G method. Each data point represents the result of three independent experiments, with error bars indicating standard deviations.

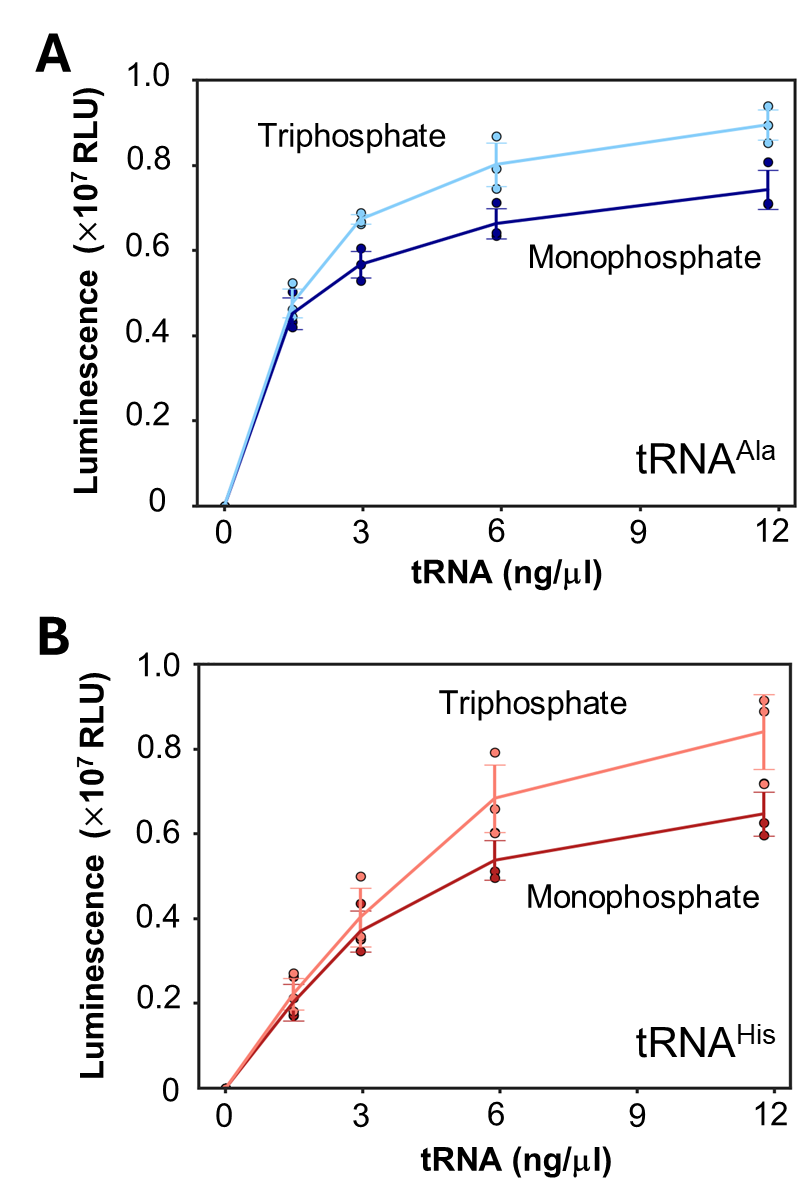

**Supplementary Figure S6. Comparison of translation activity using tRNAs with 5'-triphosphate and 5'-monophosphate ends**

(A) Comparison of the 5'-phosphate groups of tRNA^Ala^. (B) Comparison of the 5'-phosphate groups of tRNA^His^. 5’-Triphosphate tRNAs were prepared using simple in vitro transcription. 5’-Monophosphate tRNAs were prepared by in vitro transcription of a tRNA template attached to the leader sequence, followed by RNase P digestion. Each data point represents the result of three independent experiments, with error bars indicating standard deviations.

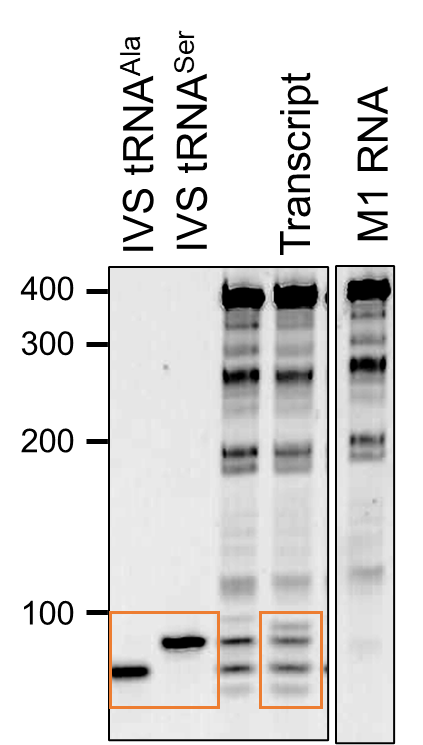

**Supplementary Figure S7. The whole image of Fig. 4E**

The parts shown in Fig. 4E were enclosed. For comparison, results for only M1 RNA are shown.

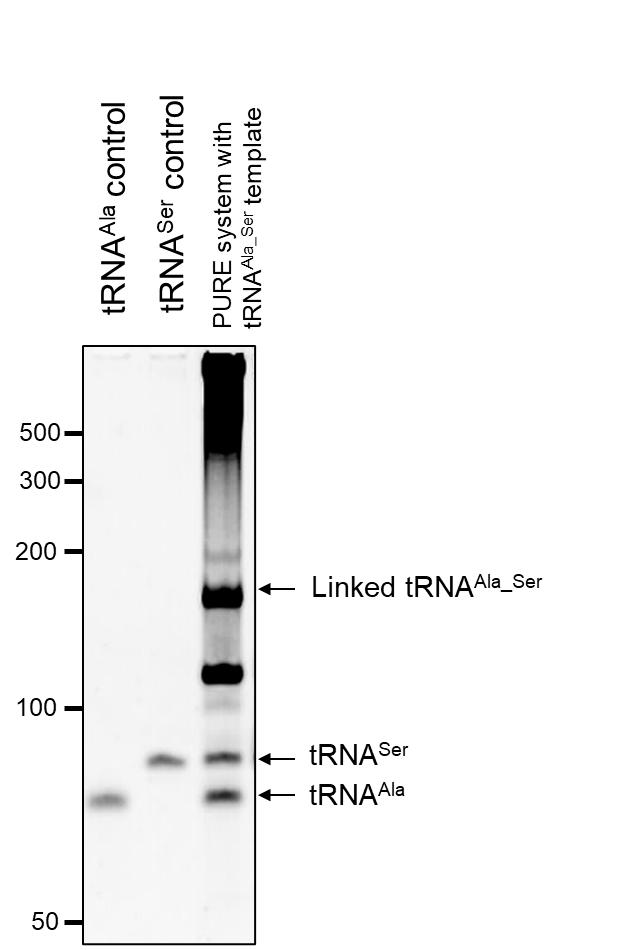

**Supplementary Figure S8. tRNA synthesis using linked tRNA method in the tfPURE system**

A reaction mixture containing 50 nM linked tRNA^Ala-Ser^ template (Linked tRNA^Ala-Ser^), RNase P (M1 RNA, 500 nM), and tfPURE (composition F) was incubated at 30°C for 16 h, followed by PAGE analysis of a 1 µL aliquot of the reaction mixture. Length controls included IVS tRNA^Ala^, IVS tRNA^Ser^

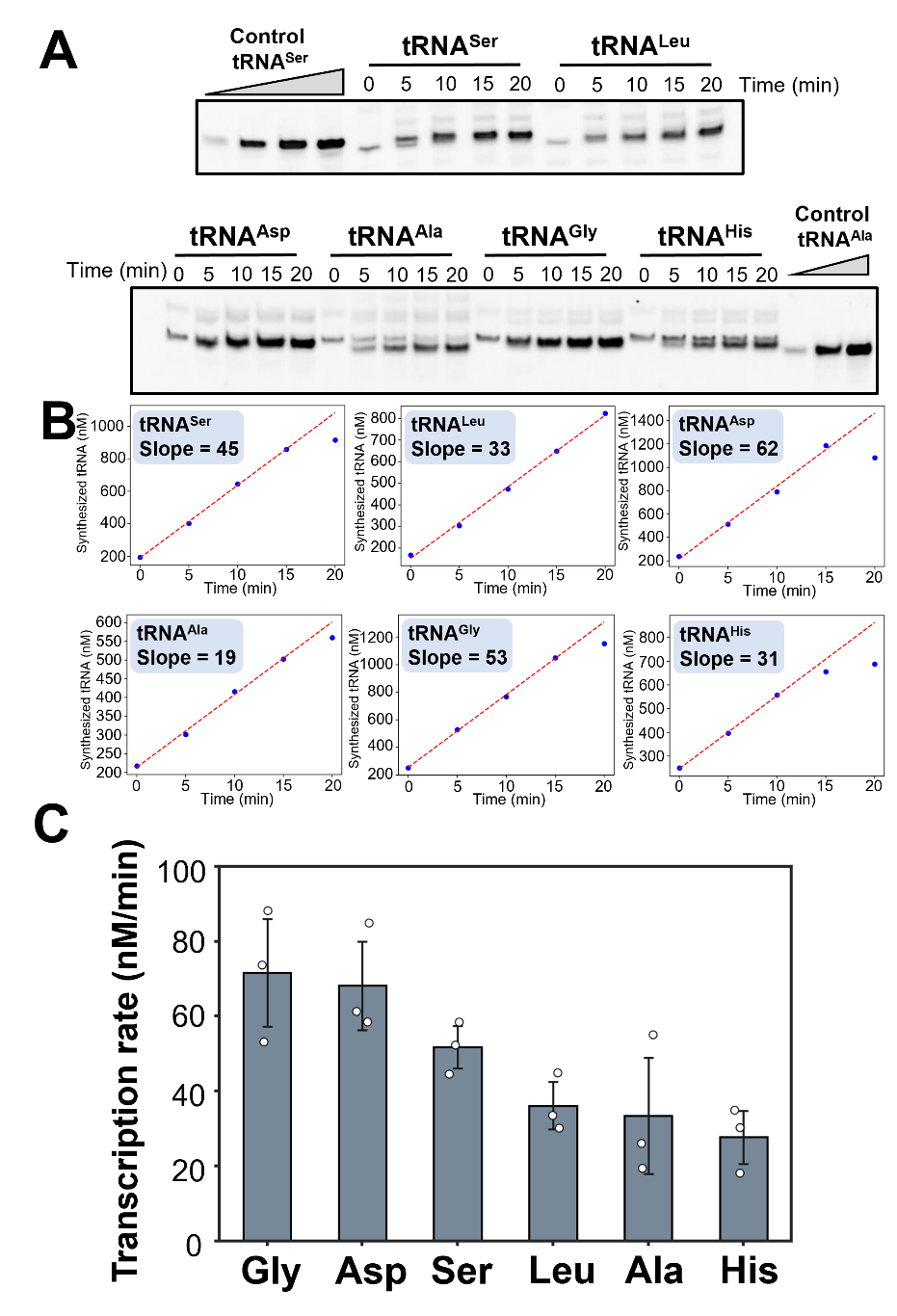

**Supplementary Figure S9. Comparison of tRNA synthesis rates**

(A) In vitro transcription was performed using tRNA templates with various tRNAs placed directly downstream of the T7 promoter, and subjected to PAGE analysis. Representative samples are shown. (B) The intensities of bands corresponding to each tRNA were quantified and plotted over time. (C) The synthesis rate was calculated from linear regression of the data shown in (B). Each data point represents three independent experiments, and error bars indicate standard deviations.

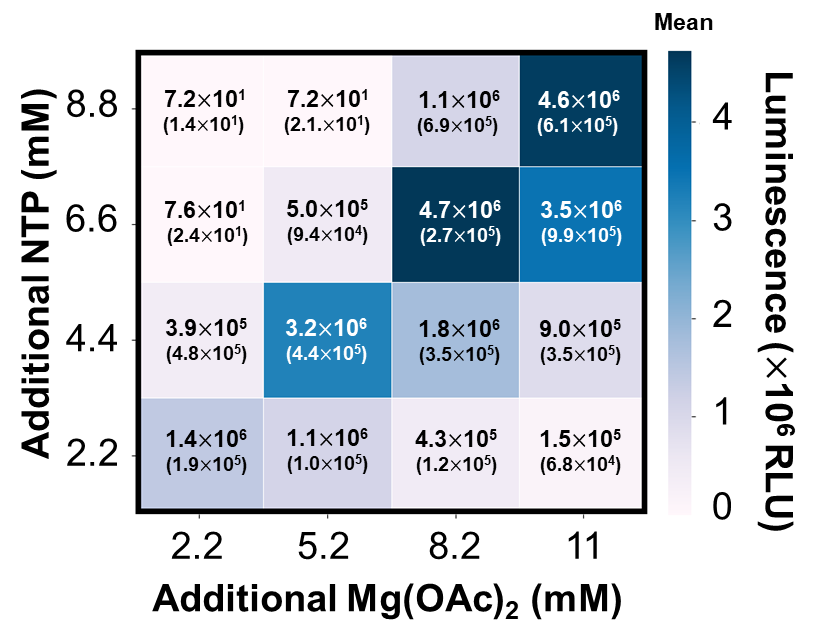

**Supplementary Figure S10. Optimization of NTP and Mg(OAc)₂ for simultaneous 21 tRNA expression**

The basic reaction mixture contains NTP (0.88 mM), Mg(OAc)₂ (7.9 mM), #1 GARCQ tRNA template (50 nM) spermidine (0.75 μM), T7 RNA polymerase (3.4 U/μL), T4 PNK (0.094 U/μL), RNase P (4 μM M1 RNA and 6 μM C5 protein) in the tfPURE system. To identify the optimal conditions for the simultaneous expression of the 21 tRNAs, additional NTP and Mg(OAc)₂ concentrations were varied in combination. The mean values of three independent experiments are shown in the box and visualized as a heatmap, with each standard deviation in parentheses. The optimum concentration (6.6 mM additional NTP and 8.2 mM additional Mg(OAc)₂) was used in Fig. 6.

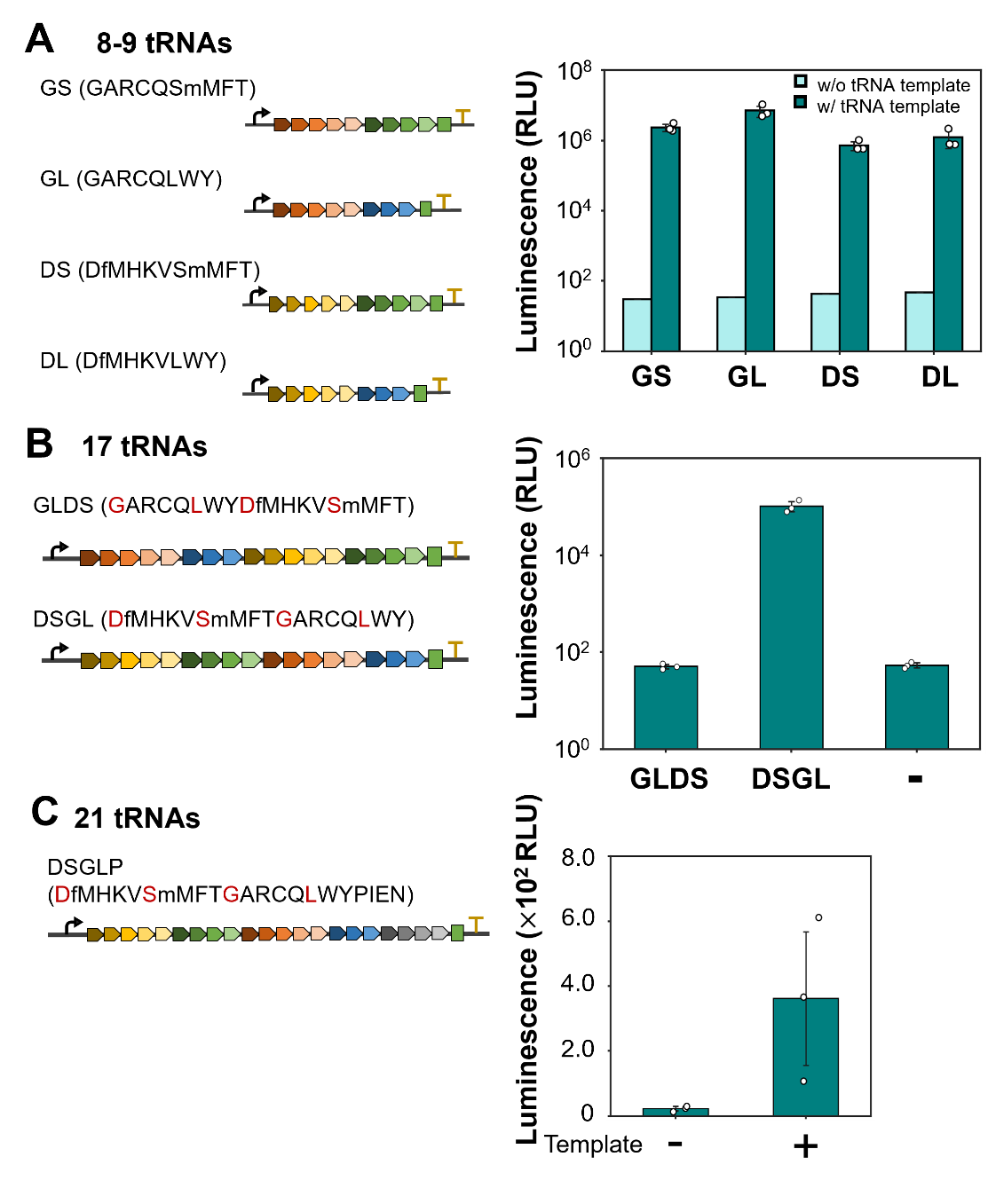

**Supplementary Figure 11. Translation-coupled reaction with tRNA arrays encoding 8-21 tRNAs**

(A) Examination of tRNA arrays with 8 or 9 linked tRNAs. Four types of tRNA templates were designed by connecting #3 (SmMFT) or #4 (LWY) to the 3’ end of either #1 (GARCQ) or #2 (DfMHKV), respectively. The tRNA templates were named based on the initials of the tRNAs (e.g., GS for GARCQ + SmMFT). The reaction mixture contained one of the tRNA templates (50 nM), luciferase template (1 nM), T7 RNA polymerase (1.7 U/μL), T4 PNK (0.094 U/μL), RNase P (1 μM of M1 RNA and 1.5 μM C5 protein), other 12 or 13 IVS tRNAs, and the tfPURE system (composition F) was incubated at 30 °C for 24 h, and luminescence was measured. (B) Examination of tRNA arrays containing the 17 linked tRNAs genes. The GL (GARCQ + LWY), which had the highest luminescence value in (A), was extended by linking the DS to either the 3’ end (GLDS) or 5’ end (DSGL). Reaction conditions were the same as in (A), except that four IVS tRNAs (IPEN) were included instead of 12 or 13 IVS tRNAs. (C) Examination of tRNAs containing 21 linked tRNAs. The DSGL in(B) was further extended by linking PIEN to the 3’ end, resulting in DSGLP. The reaction mixture containing the linear DNA encoding the linked 21 tRNAs (50 nM), luciferase template (1 nM), T7 RNA polymerase (3.4 U/μL), T4 PNK (0.094 U/μL), RNase P (4 μM M1 RNA and 6 μM C5 protein), and the tfPURE system (composition G) was incubated at 30 °C for 24 h, and luminescence was measured. Each data point represents the result of three independent experiments, with error bars indicating standard deviations.

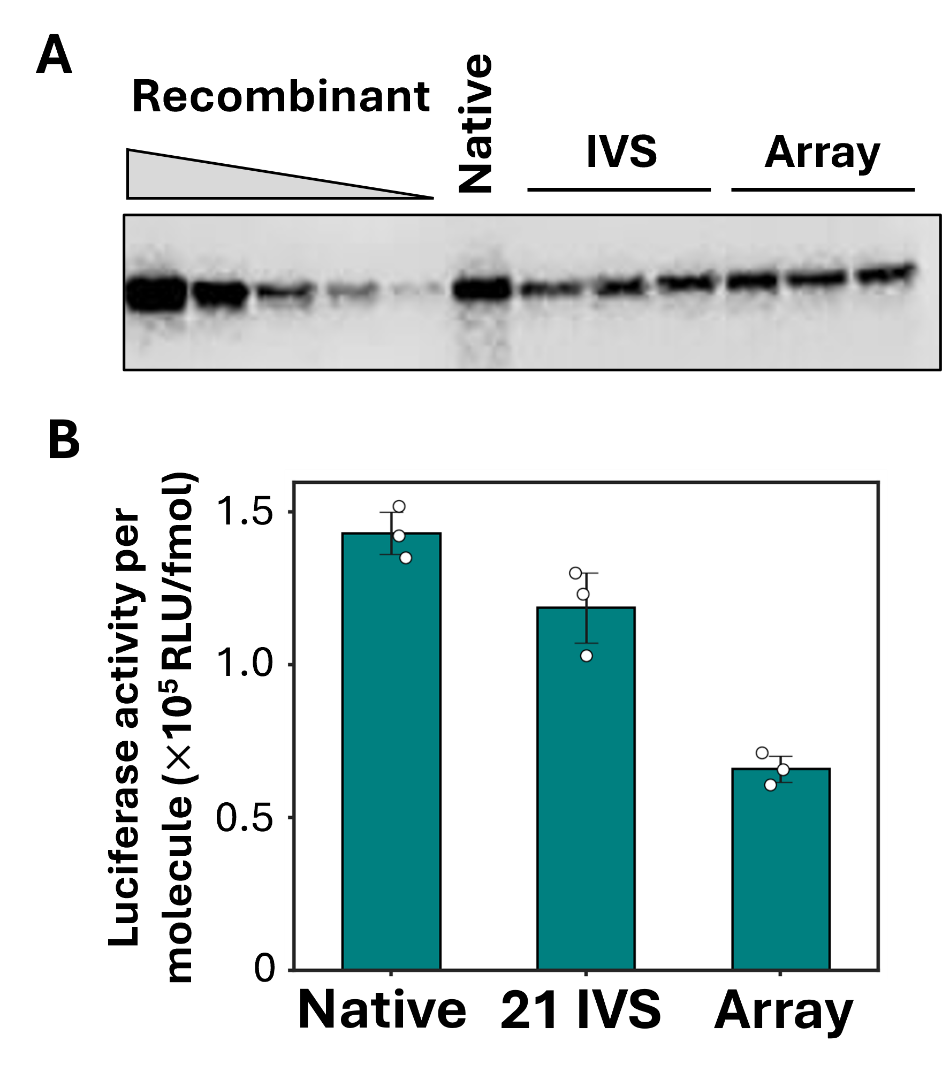

**Supplementary Figure S12. Activity per molecule of synthesized luciferase quality**

(A) Western blot analysis of the luciferase synthesized in the PURE system. The luciferase protein synthesized with E. coli native tRNA (Native), individually prepared 21 in vitro synthesized tRNA (IVS), and simultaneously prepared 21 tRNA by the tRNA array method (Array) were analyzed by western blotting for protein quantification. For the native and IVS tRNAs, the reaction mixture containing each tRNA mixture (600 ng/μL, Roche), luciferase template (1 nM), T7 RNA polymerase (0.42 U/μL), and the tfPURE system (Composition A) was incubated at 30 °C for 16 h. For simultaneous 21 expression by the tRNA array method, the reaction mixture containing the single 21 tRNAs template (50 nM), luciferase template (1 nM), T7 RNA polymerase (3.4 U/μL), T4 PNK (0.094 U/μL), RNase P (4 μM M1 RNA and 6 μM C5 protein), and the tfPURE system (Composition G) was incubated at 30 °C for 24 h. After incubation, aliquots were subjected to western blotting with recombinant luciferase of known concentrations as a control. Luciferase activity was also measured. (B) Comparison of the activity per molecule of the synthesized luciferase. The quality of the synthesized luciferase was evaluated by calculating the luminescence per molecule based on the western blot quantification results and luciferase activity. Each data point represents the result of three independent experiments, with error bars indicating standard deviations.

**Supplementary Table S1. Primer Sequences used in this study**

| Primer No. | Sequence |
| --- | --- |
| 1 | GGCGATTAAGTTGGGTAACGCCAG |
| 2 | CCGGCTCGTATGTTGTGTGG |
| 3 | AAGCGGAAGAGCGCCCAATACGC |
| 4 | TGGCTCCTCTGACTGGACTCG |
| 5 | TGGCTGGGGTACGAGGATTCG |
| 6 | TGGTAGGCCTGAGTGGACTTG |
| 7 | TGGTTGCGGGGGCCGGA |
| 8 | TGGTCGGCACGAGAGGATTT |
| 9 | TGGCAGGGGCGGAGAGACTCG |
| 10 | GAGATTAATACGACTCACTATAGCCTCTGTAGTTCAGTCGGTAGAAC |
| 11 | TGGCGCCTCTGACTGGACTCG |
| 12 | GAGATTAATACGACTCACTATAGGGGGTATCGCCAAGCGGTAAG |
| 13 | TGGCGGGGGTACGAGGATTCG |
| 14 | GAGATTAATACGACTCACTATAGGGCTTGTAGCTCAGGTGGTTAG |
| 15 | TGGTGGGCCTGAGTGGACTTGAACC |
| 16 | GAGATTAATACGACTCACTATAGGCGGGGTGGAGCAGCCTG |
| 17 | GAGATTAATACGACTCACTATTAGCGGGGTGGAGCAGCCTG |
| 18 | TGGTAGCGGGGGCCGGATTTG |
| 19 | GAGATTAATACGACTCACTATAGGGCACGTAGCGCAGCCTGG |
| 20 | TGGTGGGCACGAGAGGATTTG |
| 21 | GAGATTAATACGACTCACTATAGGGGGCGTAGTTCAATTGGTAG |
| 22 | TGGCGGGGGCGGAGAGAC |
| 23 | CCGCGTAATACGACTCACTATAGGGGCTATAGCTCAGCTG |
| 24 | TGGTGGAGCTAAGCGGGATCG |
| 25 | CCGCGTAATACGACTCACTATAGCGCCCGTAGCTCAG |
| 26 | TGGCGCGCCCGACAGGATTCG |
| 27 | CCGCGTAATACGACTCACTATAGGAGCGGTAGTTCAGTCG |
| 28 | TGGCGGAACGGACGGGACTCG |
| 29 | CCGCGTAATACGACTCACTATAGGCGCGTTAACAAAGCG |
| 30 | TGGAGGCGCGTTCCGGAGTCG |
| 31 | CCGCGTAATACGACTCACTATAGCGGGAATAGCTCAGTTG |
| 32 | TGGAGCGGGAAACGAGAC |
| 33 | CCGCGTAATACGACTCACTATAGTCCCCTTCGTCTAGAGG |
| 34 | TGGCGTCCCCTAGGGGATTCG |
| 35 | CCGCGTAATACGACTCACTATAGGTGGCTATAGCTCAGTTG |
| 36 | TGGGGTGGCTAATGGGATTCG |
| 37 | CCGCGTAATACGACTCACTATAGCGAAGGTGGCGGAATTG |
| 38 | TGGTGCGAGGGGGGGGA |
| 39 | CCGCGTAATACGACTCACTATAGGGTCGTTAGCTCAGTTGG |
| 40 | TGGTGGGTCGTGCAGGATT |
| 41 | CCGCGTAATACGACTCACTATAGGCTACGTAGCTCAGTTG |
| 42 | TGGTGGCTACGACGGGATTCG |
| 43 | CCGCGTAATACGACTCACTATAGCCCGGATAGCTCAGTC |
| 44 | TGGTGCCCGGACTCGGAA |
| 45 | CCGCGTAATACGACTCACTATAGGTGAGGTGTCCGAGTG |
| 46 | TGGCGGTGAGGGGGGGATTCG |
| 47 | CCGCGTAATACGACTCACTATAGCTGATATGGCTCAGTTGG |
| 48 | TGGTGCTGATACCCAGAGTCG |
| 49 | CCGCGTAATACGACTCACTATAGGTGGGGTTCCCGAG |
| 50 | TGGTGGTGGGGGAAGGATTCG |
| 51 | CCGCGTAATACGACTCACTATAGCGTCCGTAGCTCAGTTG |
| 52 | TGGTGCGTCCGAGTGGACTCG |
| 53 | GGGGCTGCAGTAATACGACTCACTATA |
| 54 | AGGTGAAACTGACCGATAAG |
| 55 | CCGCGTAATACGACTCACTATAGGTGAGGTGTCCGAGTG |
| 56 | GCCAGGTTATCCGGATATAGTTCCTC |
| 57 | CCGCGTAATACGACTCACTATAGGGGCTATAGCTCAGCTG |
| 58 | TGGCGGTGAGGGGGGGATTCG |
| 59 | GTAGAGCGCACCCTTCGTAAGGGTGAGGTCCCCAG |
| 60 | AAGGGTGCGCTCTACCAACTGAGC |
| 61 | TAGACGCGCTAGCTTCGTGTGTTAGTGTCCTTAC |
| 62 | AAGCTAGCGCGTCTACCAATTCCGC |

**Supplementary Table S2. Primer sets used for preparing PCR products in the experiments**

| Primer sets | Template | PCR product | Purpose  (Experiment) |
| --- | --- | --- | --- |
| 1,2 | pUC-T7p-Fluc_21tRNA | Linear_Fluc_21tRNA | Fig.1C,D  Fig.2B,C,D  Fig.3  Fig.4C,F,H  Fig.5B,C,E  Fig.6A,B  Fig.S1C  Fig.S2-7  Fig.S12-14 |
| 1,2 | Nanoluc_21tRNA_flagment | Linear_Nanoluc_21tRNA | Fig.7C |
| 1,2 | Nanoluc_Thr ACG_to_Leu_flagment | Linear_Nanoluc_Thr_Leu_ACG | Fig.7D,E,F |
| 3,4 | pGEMEX_tRNA^Asn^GUU | Linear_leader_tRNA^Asn^ | Fig.1C,D,F  Fig.3  Fig.S3,6 |
| 3,5 | pGEMEX_tRNA^Gln^CUG | Linear_leader_tRNA^Gln^ | Fig.1C,E,D  Fig.S2,3,6 |
| 3,6 | pGEMEX_tRNA^Ile^GAU | Linear_leader_tRNA^Ile^ | Fig.1C,D,F  Fig.S3,6 |
| 3,7 | pGEMEX_tRNA^fMet^CAU | Linear_leader_tRNA^fMet^ | Fig.1C,D,F  Fig.S3,6 |
| 3,8 | pGEMEX_tRNA^Pro^GGG | Linear_leader_tRNA^Pro^ | Fig.1C,D,F  Fig.S3,6 |
| 3,9 | pGEMEX_tRNA^Trp^CCA | Linear_leader_tRNA^Trp^ | Fig.1C,D,F  Fig.S3,6 |
| 10,4 | pGEMEX_tRNA^Asn^GUU | Linear_5G\|A_tRNA^Asn^ | Fig.2B(IVT) |
| 10,11 | pGEMEX_tRNA^Asn^GUU | Linear_5G\|C_tRNA^Asn^ | Fig.2B(IVT)  Fig.2C,D,E  Fig.S5 |
| 12,5 | pGEMEX_tRNA^Gln^CUG | Linear_5G\|A_tRNA^Gln^ | Fig.2B(IVT) |
| 12,13 | pGEMEX_tRNA^Gln^CUG | Linear_5G\|C_tRNA^Gln^ | Fig.2B(IVT)  Fig.2C,D,E  Fig.3  Fig.S5 |
| 14,6 | pGEMEX_tRNA^Ile^GAU | Linear_5G\|U_tRNA^Ile^ | Fig.2B(IVT)  Fig.2C,D,E  Fig.3  Fig.S5 |
| 14,15 | pGEMEX_tRNA^Ile^GAU | Linear_5G\|C_tRNA^Ile^ | Fig.2B(IVT) |
| 16,7 | pGEMEX_tRNA^fMet^CAU | Linear_5G\|A_tRNA^fMet^ | Fig.2B(IVT)  Fig.2C,D,E  Fig.3  Fig.S5 |
| 16,18 | pGEMEX_tRNA^fMet^CAU | Linear_5G\|U_tRNA^fMet^ | Fig.2B(IVT) |
| 17,7 | pGEMEX_tRNA^fMet^CAU | Linear_5A\|A_tRNA^fMet^ | Fig.2B(IVT) |
| 17,18 | pGEMEX_tRNA^fMet^CAU | Linear_5A\|U_tRNA^fMet^ | Fig.2B(IVT) |
| 19,8 | pGEMEX_tRNA^Pro^GGG | Linear_5G\|G_tRNA^Pro^ | Fig.2B(IVT)  Fig.2C,D,E  Fig.3  Fig.S5 |
| 19,20 | pGEMEX_tRNA^Pro^GGG | Linear_5G\|C_tRNA^Pro^ | Fig.2B(IVT) |
| 21,9 | pGEMEX_tRNA^Trp^CCA | Linear_5G\|U_tRNA^Trp^ | Fig.2B(IVT) |
| 21,22 | pGEMEX_tRNA^Trp^CCA | Linear_5G\|C_tRNA^Trp^ | Fig.2B(IVT)  Fig.2C,D,E  Fig.3  Fig.S5 |
| 23,24 | pGEMEX_tRNA^Ala^GGC | Linear_tRNA^Ala^ | IVT tRNA preparation |
| 25,26 | pGEMEX_tRNA^Arg^CCG | Linear_tRNA^Arg^ |  |
| 27,28 | pGEMEX_tRNA^Asp^GUC | Linear_tRNA^Asp^ |  |
| 29,30 | pGEMEX_tRNA^Cys^GCA | Linear_tRNA^Cys^ |  |
| 31,32 | pGEMEX_tRNA^Gly^GCG | Linear_tRNA^Gly^ |  |
| 33,34 | pGEMEX_tRNA^Glu^GUC | Linear_tRNA^Glu^ |  |
| 35,36 | pGEMEX_tRNA^His^GUG | Linear_tRNA^His^ |  |
| 37,38 | pGEMEX_tRNA^Leu^CAG | Linear_tRNA^Leu^ |  |
| 39,40 | pGEMEX_tRNA^Lys^CAG | Linear_tRNA^Lye^ |  |
| 41,42 | pGEMEX_tRNA^mMet^CAU | Linear_tRNA^mMet^ |  |
| 43,44 | pGEMEX_tRNA^Phe^GAA | Linear_tRNA^Phe^ |  |
| 45,46 | pGEMEX_tRNA^Ser^GGA | Linear_tRNA^Ser^ |  |
| 47,48 | pGEMEX_tRNA^Thr^GGU | Linear_tRNA^Thr^ |  |
| 49,50 | pGEMEX_tRNA^Tyr^GUA | Linear_tRNA^Tyr^ |  |
| 51,52 | pGEMEX_tRNA^Val^GAC | Linear_tRNA^Val^ |  |
| 53.54 | pUC_M1 | M1_IVT_Template | IVT |
| 55,56 | pGEMEX_tRNA^Ala_Ser^_HDVR_T7hyb10 | Linear_tRNA^Ser^_HDVR | Fig.4A,B,C |
| 57,58 | pGEMEX_tRNA^Ala_Ser^_HDVR_T7hyb10 | Linear_tRNA^Ala_Ser^ | Fig.4D,E,F  Fig.S8,9 |
| 3,56 | pGEMEX_tRNA^Ala_Ser^_HDVR_T7hyb10 | Linear_tRNA^Ala_Ser^_HDVR | Fig.4G,H |
| 3,56 | pTwist_tRNA^GARCQ^-HDVR | Linear_tRNA^GARCQ^ | Fig.5A,B,C,D |
| 3,56 | pTwist_tRNA^DfMHKV^-HDVR | Linear_tRNA^DfMHKV^ | Fig.5A,B,C,D |
| 3,56 | pTwist_tRNA^SmMFT^-HDVR | Linear_tRNA^SmMFT^ | Fig.5A,B,C,D |
| 3,56 | pTwist_tRNA^LWY^-HDVR | Linear_tRNA^LWY^ | Fig.5A,B,C,D |
| 3,56 | pTwist_tRNA^IPEN^-HDVR | Linear_tRNA^IPEN^ | Fig.5A,B,E,F |
| 3,56 | pTwist_tRNA^PIEN^-HDVR | Linear_tRNA^PIEN^ | Fig.5A,B,E,F |
| 3,56 | pTwist_tRNA^EIPN^-HDVR | Linear_tRNA^EIPN^ | Fig.5A,B,E,F |
| 3,56 | pTwist_tRNA^NIPE^-HDVR | Linear_tRNA^NIPE^ | Fig.5A,B,E,F |
| 3,56 | pTwist_21 tRNAs | Linear_21 tRNAs | Fig.6A,B,C  Fig.7A,B  Fig.S12,14 |
| 3,56 | pTwist_21 tRNAs^DSGLP^_all_linked | Linear_21 tRNAs^DSGLP^_all_linked | Fig.S13 |

**Supplementary Table S3. Sequences of plasmid DNA and fragments used in this study**

| Template | Sequence |
| --- | --- |
| pUC-T7p-Fluc_21tRNA  *Fluc_gene | TCGCGCGTTTCGGTGATGACGGTGAAAACCTCTGACACATGCAGCTCCCGGAGACGGTCACAGCTTGTCTGTAAGCGGATGCCGGGAGCAGACAAGCCCGTCAGGGCGCGTCAGCGGGTGTTGGCGGGTGTCGGGGCTGGCTTAACTATGCGGCATCAGAGCAGATTGTACTGAGAGTGCACCATATGCGGTGTGAAATACCGCACAGATGCGTAAGGAGAAAATACCGCATCAGGCGCCATTCGCCATTCAGGCTGCGCAACTGTTGGGAAGGGCGATCGGTGCGGGCCTCTTCGCTATTACGCCAGCTGGCGAAAGGGGGATGTGCTGCAAGGCGATTAAGTTGGGTAACGCCAGGGTTTTCCCAGTCACGACGTTGTAAAACGACGGCCAGTGAATTCTAATACGACTCACTATAGGGGAATTGTGAGCGGATAACAATTCCCCTCTAGAAATAATTTTGTTTAACTTTAAGAAGGAGATATACATATGGAGGACGCTAAGAACATTAAGAAGGGTCCTGCTCCTTTTTACCCTCTGGAGGACGGTACTGCTGGTGAGCAGCTGCACAAGGCTATGAAGCGGTACGCTCTGGTTCCTGGTACTATTGCTTTTACTGACGCTCACATTGAGGTTAACATTACTTACGCTGAGTACTTTGAGATGTCTGTTCGGCTGGCTGAGGCTATGAAGCGGTACGGTCTGAACACTAACCACCGGATTGTTGTTTGTTCTGAGAACTCTCTGCAGTTTTTTATGCCTGTTCTGGGTGCTCTGTTTATTGGTGTTGCTGTTGCTCCTGCTAACGACATTTACAACGAGCGGGAGCTGCTGAACTCTATGAACATTTCTCAGCCTACTGTTGTTTTTGTTTCTAAGAAGGGTCTGCAGAAGATTCTGAACGTTCAGAAGAAGCTGCCTATTATTCAGAAGATTATTATTATGGACTCTAAGACTGACTACCAGGGTTTTCAGTCTATGTACACTTTTGTTACTTCTCACCTGCCTCCTGGTTTTAACGAGTACGACTTTGTTCCTGAGTCTTTTGACCGGGACAAGACTATTGCTCTGATTATGAACTCTTCTGGTTCTACTGGTCTGCCTAAGGGTGTTGCTCTGCCTCACCGGACTGCTTGTGTTCGGTTTTCTCACGCTCGGGACCCTATTTTTGGTAACCAGATTATTCCTGACACTGCTATTCTGTCTGTTGTTCCTTTTCACCACGGTTTTGGTATGTTTACTACTCTGGGTTACCTGATTTGTGGTTTTCGGGTTGTTCTGATGTACCGGTTTGAGGAGGAGCTGTTTCTGCGGTCTCTGCAGGACTACAAGATTCAGTCTGCTCTGCTGGTTCCTACTCTGTTTTCTTTTTTTGCTAAGTCTACTCTGATTGACAAGTACGACCTGTCTAACCTGCACGAGATTGCTTCTGGTGGTGCTCCTCTGTCTAAGGAGGTTGGTGAGGCTGTTGCTAAGCGGTTTCACCTGCCTGGTATTCGGCAGGGTTACGGTCTGACTGAGACTACTTCTGCTATTCTGATTACTCCTGAGGGTGACGACAAGCCTGGTGCTGTTGGTAAGGTTGTTCCTTTTTTTGAGGCTAAGGTTGTTGACCTGGACACTGGTAAGACTCTGGGTGTTAACCAGCGGGGTGAGCTGTGTGTTCGGGGTCCTATGATTATGTCTGGTTACGTTAACAACCCTGAGGCTACTAACGCTCTGATTGACAAGGACGGTTGGCTGCACTCTGGTGACATTGCTTACTGGGACGAGGACGAGCACTTTTTTATTGTTGACCGGCTGAAGTCTCTGATTAAGTACAAGGGTTACCAGGTTGCTCCTGCTGAGCTGGAGTCTATTCTGCTGCAGCACCCTAACATTTTTGACGCTGGTGTTGCTGGTCTGCCTGACGACGACGCTGGTGAGCTGCCTGCTGCTGTTGTTGTTCTGGAGCACGGTAAGACTATGACTGAGAAGGAGATTGTTGACTACGTTGCTTCTCAGGTTACTACTGCTAAGAAGCTGCGGGGTGGTGTTGTTTTTGTTGACGAGGTTCCTAAGGGTCTGACTGGTAAGCTGGACGCTCGGAAGATTCGGGAGATTCTGATTAAGGCTAAGAAGGGTGGTAAGTCTAAGCTGTAAAAGCTTGGCGTAATCATGGTCATAGCTGTTTCCTGTGTGAAATTGTTATCCGCTCACAATTCCACACAACATACGAGCCGGAAGCATAAAGTGTAAAGCCTGGGGTGCCTAATGAGTGAGCTAACTCACATTAATTGCGTTGCGCTCACTGCCCGCTTTCCAGTCGGGAAACCTGTCGTGCCAGCTGCATTAATGAATCGGCCAACGCGCGGGGAGAGGCGGTTTGCGTATTGGGCGCTCTTCCGCTTCCTCGCTCACTGACTCGCTGCGCTCGGTCGTTCGGCTGCGGCGAGCGGTATCAGCTCACTCAAAGGCGGTAATACGGTTATCCACAGAATCAGGGGATAACGCAGGAAAGAACATGTGAGCAAAAGGCCAGCAAAAGGCCAGGAACCGTAAAAAGGCCGCGTTGCTGGCGTTTTTCCATAGGCTCCGCCCCCCTGACGAGCATCACAAAAATCGACGCTCAAGTCAGAGGTGGCGAAACCCGACAGGACTATAAAGATACCAGGCGTTTCCCCCTGGAAGCTCCCTCGTGCGCTCTCCTGTTCCGACCCTGCCGCTTACCGGATACCTGTCCGCCTTTCTCCCTTCGGGAAGCGTGGCGCTTTCTCATAGCTCACGCTGTAGGTATCTCAGTTCGGTGTAGGTCGTTCGCTCCAAGCTGGGCTGTGTGCACGAACCCCCCGTTCAGCCCGACCGCTGCGCCTTATCCGGTAACTATCGTCTTGAGTCCAACCCGGTAAGACACGACTTATCGCCACTGGCAGCAGCCACTGGTAACAGGATTAGCAGAGCGAGGTATGTAGGCGGTGCTACAGAGTTCTTGAAGTGGTGGCCTAACTACGGCTACACTAGAAGAACAGTATTTGGTATCTGCGCTCTGCTGAAGCCAGTTACCTTCGGAAAAAGAGTTGGTAGCTCTTGATCCGGCAAACAAACCACCGCTGGTAGCGGTGGTTTTTTTGTTTGCAAGCAGCAGATTACGCGCAGAAAAAAAGGATCTCAAGAAGATCCTTTGATCTTTTCTACGGGGTCTGACGCTCAGTGGAACGAAAACTCACGTTAAGGGATTTTGGTCATGAGATTATCAAAAAGGATCTTCACCTAGATCCTTTTAAATTAAAAATGAAGTTTTAAATCAATCTAAAGTATATATGAGTAAACTTGGTCTGACAGTTACCAATGCTTAATCAGTGAGGCACCTATCTCAGCGATCTGTCTATTTCGTTCATCCATAGTTGCCTGACTCCCCGTCGTGTAGATAACTACGATACGGGAGGGCTTACCATCTGGCCCCAGTGCTGCAATGATACCGCGAGACCCACGCTCACCGGCTCCAGATTTATCAGCAATAAACCAGCCAGCCGGAAGGGCCGAGCGCAGAAGTGGTCCTGCAACTTTATCCGCCTCCATCCAGTCTATTAATTGTTGCCGGGAAGCTAGAGTAAGTAGTTCGCCAGTTAATAGTTTGCGCAACGTTGTTGCCATTGCTACAGGCATCGTGGTGTCACGCTCGTCGTTTGGTATGGCTTCATTCAGCTCCGGTTCCCAACGATCAAGGCGAGTTACATGATCCCCCATGTTGTGCAAAAAAGCGGTTAGCTCCTTCGGTCCTCCGATCGTTGTCAGAAGTAAGTTGGCCGCAGTGTTATCACTCATGGTTATGGCAGCACTGCATAATTCTCTTACTGTCATGCCATCCGTAAGATGCTTTTCTGTGACTGGTGAGTACTCAACCAAGTCATTCTGAGAATAGTGTATGCGGCGACCGAGTTGCTCTTGCCCGGCGTCAATACGGGATAATACCGCGCCACATAGCAGAACTTTAAAAGTGCTCATCATTGGAAAACGTTCTTCGGGGCGAAAACTCTCAAGGATCTTACCGCTGTTGAGATCCAGTTCGATGTAACCCACTCGTGCACCCAACTGATCTTCAGCATCTTTTACTTTCACCAGCGTTTCTGGGTGAGCAAAAACAGGAAGGCAAAATGCCGCAAAAAAGGGAATAAGGGCGACACGGAAATGTTGAATACTCATACTCTTCCTTTTTCAATATTATTGAAGCATTTATCAGGGTTATTGTCTCATGAGCGGATACATATTTGAATGTATTTAGAAAAATAAACAAATAGGGGTTCCGCGCACATTTCCCCGAAAAGTGCCACCTGACGTCTAAGAAACCATTATTATCATGACATTAACCTATAAAAATAGGCGTATCACGAGGCCCTTTCGTC |
| Nanoluc_21tRNA_flagment  *Nannoluc gene | GGCGATTAAGTTGGGTAACGCCAGGGTTTTCCCAGTCACGACGTTGTAAAACGACGGCCAGTGAATTCTAATACGACTCACTATAGGGGAATTGTGAGCGGATAACAATTCCCCTCTAGAAATAATTTTGTTTAACTTTAAGAAGGAGATATACATATGGTTTTTACTCTGGAGGACTTTGTTGGTGACTGGCGGCAGACTGCTGGTTACAACCTGGACCAGGTTCTGGAGCAGGGTGGTGTTTCTTCTCTGTTTCAGAACCTGGGTGTTTCTGTTACTCCTATTCAGCGGATTGTTCTGTCTGGTGAGAACGGTCTGAAGATTGACATTCACGTTATTATTCCTTACGAGGGTCTGTCTGGTGACCAGATGGGTCAGATTGAGAAGATTTTTAAGGTTGTTTACCCTGTTGACGACCACCACTTTAAGGTTATTCTGCACTACGGTACTCTGGTTATTGACGGTGTTACTCCTAACATGATTGACTACTTTGGTCGGCCTTACGAGGGTATTGCTGTTTTTGACGGTAAGAAGATTACTGTTACTGGTACTCTGTGGAACGGTAACAAGATTATTGACGAGCGGCTGATTAACCCTGACGGTTCTCTGCTGTTTCGGGTTACTATTAACGGTGTTACTGGTTGGCGGCTGTGTGAGCGGATTCTGGCTTAAAAGCTTGGCGTAATCATGGTCATAGCTGTTTCCTGTGTGAAATTGTTATCCGCTCACAATTCCACACAACATACGAGCCGG |
| Nanoluc_Thr ACG_to_Leu_flagment  *Nannoluc gene | GGCGATTAAGTTGGGTAACGCCAGGGTTTTCCCAGTCACGACGTTGTAAAACGACGGCCAGTGAATTCTAATACGACTCACTATAGGGGAATTGTGAGCGGATAACAATTCCCCTCTAGAAATAATTTTGTTTAACTTTAAGAAGGAGATATACATATGGTTTTTACTCTGGAGGACTTTGTTGGTGACTGGCGGCAGACTGCTGGTTACAACCTGGACCAGGTTACGGAGCAGGGTGGTGTTTCTTCTCTGTTTCAGAACCTGGGTGTTTCTGTTACTCCTATTCAGCGGATTGTTACGTCTGGTGAGAACGGTCTGAAGATTGACATTCACGTTATTATTCCTTACGAGGGTCTGTCTGGTGACCAGATGGGTCAGATTGAGAAGATTTTTAAGGTTGTTTACCCTGTTGACGACCACCACTTTAAGGTTATTACGCACTACGGTACTCTGGTTATTGACGGTGTTACTCCTAACATGATTGACTACTTTGGTCGGCCTTACGAGGGTATTGCTGTTTTTGACGGTAAGAAGATTACTGTTACTGGTACTCTGTGGAACGGTAACAAGATTATTGACGAGCGGACGATTAACCCTGACGGTTCTCTGCTGTTTCGGGTTACTATTAACGGTGTTACTGGTTGGCGGACGTGTGAGCGGATTCTGGCTTAAAAGCTTGGCGTAATCATGGTCATAGCTGTTTCCTGTGTGAAATTGTTATCCGCTCACAATTCCACACAACATACGAGCCGG |
| pTwist_21tRNAs  *GARCQ group  *DfMHKV group  *SmMFT group  *LWY group  *PIEN group | AAGCGGAAGAGCGCCCAATACGCAAACCGCCTCTCCCCGCGCGTTGGCCGATTCATTAATGCAGCCCGGGAGATCTCGATCCCGCGAAAAATTCTAATACGACTCACTATAGCGGGAATAGCTCAGTTGGTAGAGCACGACCTTGCCAAGGTCGGGGTCGCGAGTTCGAGTCTCGTTTCCCGCTCCAGGGGCTATAGCTCAGCTGGGAGAGCGCTTGCATGGCATGCAAGAGGTCAGCGGTTCGATCCCGCTTAGCTCCACCAGCGCCCGTAGCTCAGCTGGATAGAGCGCTGCCCTCCGGAGGCAGAGGTCTCAGGTTCGAATCCTGTCGGGCGCGCCAGGCGCGTTAACAAAGCGGTTATGTAGCGGATTGCAAATCCGTCTAGTCCGGTTCGACTCCGGAACGCGCCTCCATGGGGTATCGCCAAGCGGTAAGGCACCGGATTCTGATTCCGGCATTCCGAGGTTCGAATCCTCGTACCCCAGCCAGGGTCGGCATGGCATCTCCACCTCCTCGCGGTCCGACCTGGGCTACTTCGGTAGGCTAAGGGAGAAGAGATAACAGATACTTCGGTATCTGTTATCTGTTTTTTTTCAACAGATAGCCGCGTTCGCGCGGCTATCTGTTTTTTTTAGGCTAGGTGGAGGCTCAGTGATGCCGTACTGGCTAAAACCTACGGCGGTGCTGCTCGCGCATTCGACATTCTAATACGACTCACTATAGCGAAGGTGGCGGAATTGGTAGACGCGCTAGCTTCAGGTGTTAGTGTCCTTACGGACGTGGGGGTTCAAGTCCCCCCCCTCGCACCAAGGGGCGTAGTTCAATTGGTAGAGCACCGGTCTCCAAAACCGGGTGTTGGGAGTTCGAGTCTCTCCGCCCCTGCCAGGTGGGGTTCCCGAGCGGCCAAAGGGAGCAGACTGTAAATCTGCCGTCACAGACTTCGAAGGTTCGAATCCTTCCCCCACCACCAGGGTCGGCATGGCATCTCCACCTCCTCGCGGTCCGACCTGGGCTACTTCGGTAGGCTAAGGGAGAAGAGATAACAGATACTTCGGTATCTGTTATCTGTTTTTTTTCAACAGATAGCCGCGTTCGCGCGGCTATCTGTTTTTTTTAGGCTAGGTGGAGGCTCAGTGATGAGATCGATAACGCGCCGGAAGAAAAAGCTCGTGGTATCACCATCAAATTCTAATACGACTCACTATAGGAGCGGTAGTTCAGTCGGTTAGAATACCTGCCTGTCACGCAGGGGGTCGCGGGTTCGAGTCCCGTCCGTTCCGCCACGCGGGGTGGAGCAGCCTGGTAGCTCGTCGGGCTCATAACCCGAAGATCGTCGGTTCAAATCCGGCCCCCGCAACCAGGTGGCTATAGCTCAGTTGGTAGAGCCCTGGATTGTGATTCCAGTTGTCGTGGGTTCGAATCCCATTAGCCACCCCAGGGTCGTTAGCTCAGTTGGTAGAGCAGTTGACTCTTAATCAATTGGTCGCAGGTTCGAATCCTGCACGACCCACCAGCGTCCGTAGCTCAGTTGGTTAGAGCACCACCTTGACATGGTGGGGGTCGGTGGTTCGAGTCCACTCGGACGCACCAGGGTCGGCATGGCATCTCCACCTCCTCGCGGTCCGACCTGGGCTACTTCGGTAGGCTAAGGGAGAAGAGATAACAGATACTTCGGTATCTGTTATCTGTTTTTTTTCAACAGATAGCCGCGTTCGCGCGGCTATCTGTTTTTTTTAGGCTAGGTGGAGGCTCAGTGATGACACTTCTCACGTTGAATACGACACCCCGACCCGTCACTACGCACAATTCTAATACGACTCACTATAGGTGAGGTGTCCGAGTGGCTGAAGGAGCACGCCTGGAAAGTGTGTATACGGCAACGTATCGGGGGTTCGAATCCCCCCCTCACCGCCAGGCTACGTAGCTCAGTTGGTTAGAGCACATCACTCATAATGATGGGGTCACAGGTTCGAATCCCGTCGTAGCCACCAGCCCGGATAGCTCAGTCGGTAGAGCAGGGGATTGAAAATCCCCGTGTCCTTGGTTCGATTCCGAGTCCGGGCACCAGCTGATATGGCTCAGTTGGTAGAGCGCACCCTTGGTAAGGGTGAGGTCCCCAGTTCGACTCTGGGTATCAGCACCAGGGTCGGCATGGCATCTCCACCTCCTCGCGGTCCGACCTGGGCTACTTCGGTAGGCTAAGGGAGAAGAGATAACAGATACTTCGGTATCTGTTATCTGTTTTTTTTCAACAGATAGCCGCGTTCGCGCGGCTATCTGTTTTTTTTAGGCTAGGTGGAGGCTCAGTGATGACGTAGACTGCCCGGGGCACGCCGACTATGTTAAAAACATGATCAAATTCTAATACGACTCACTATAGGGCACGTAGCGCAGCCTGGTAGCGCACCGTCATGGGGTGTCGGGGGTCGGAGGTTCAAATCCTCTCGTGCCGACCAAGGCTTGTAGCTCAGGTGGTTAGAGCGCACCCCTGATAAGGGTGAGGTCGGTGGTTCAAGTCCACTCAGGCCTACCAGTCCCCTTCGTCTAGAGGCCCAGGACACCGCCCTCTCACGGCGGTAACAGGGGTTCGAATCCCCTAGGGGACGCCATCCTCTGTAGTTCAGTCGGTAGAACGGCGGACTGTTAATCCGTATGTCACTGGTTCGAGTCCAGTCAGAGGAGCCAGGGTCGGCATGGCATCTCCACCTCCTCGCGGTCCGACCTGGGCTACTTCGGTAGGCTAAGGGAGAAGAGATAACAGATACTTCGGTATCTGTTATCTGTTTTTTTTCAACAGATAGCCGCGTTCGCGCGGCTATCTGTTTTTTTTGCTGAAAGGAGGAACTATATCCGGATAACCTGGCAGGCTAGGTGGAGGCTCAGTGATGATAAGTCTGCGATGGTGGATGCATGTGTCATGGTCATAGCTGTTTCCTGTGTGAAATTGTTATCCGCTCAGAGGGCACAATCCTATTCCGCGCTATCCGACAATCTCCAAGACATTAGGTGGAGTTCAGTTCGGCGTATGGCATATGTCGCTGGAAAGAACATGTGAGCAAAAGGCCAGCAAAAGGCCAGGAACCGTAAAAAGGCCGCGTTGCTGGCGTTTTTCCATAGGCTCCGCCCCCCTGACGAGCATCACAAAAATCGACGCTCAAGTCAGAGGTGGCGAAACCCGACAGGACTATAAAGATACCAGGCGTTTCCCCCTGGAAGCTCCCTCGTGCGCTCTCCTGTTCCGACCCTGCCGCTTACCGGATACCTGTCCGCCTTTCTCCCTTCGGGAAGCGTGGCGCTTTCTCATAGCTCACGCTGTAGGTATCTCAGTTCGGTGTAGGTCGTTCGCTCCAAGCTGGGCTGTGTGCACGAACCCCCCGTTCAGCCCGACCGCTGCGCCTTATCCGGTAACTATCGTCTTGAGTCCAACCCGGTAAGACACGACTTATCGCCACTGGCAGCAGCCACTGGTAACAGGATTAGCAGAGCGAGGTATGTAGGCGGTGCTACAGAGTTCTTGAAGTGGTGGCCTAACTACGGCTACACTAGAAGAACAGTATTTGGTATCTGCGCTCTGCTGAAGCCAGTTACCTTCGGAAAAAGAGTTGGTAGCTCTTGATCCGGCAAACAAACCACCGCTGGTAGCGGTGGTTTTTTTGTTTGCAAGCAGCAGATTACGCGCAGAAAAAAAGGATCTCAAGAAGATCCTTTGATCTTTTCTACGGGGTCTGACGCTCTATTCAACAAAGCCGCCGTCCCGTCAAGTCAGCGTAAATGGGTAGGGGGCTTCAAATCGTCCTCGTGATACCAATTCGGAGCCTGCTTTTTTGTACAAACTTGTTGATAATGGCAATTCAAGGATCTTCACCTAGATCCTTTTAAATTAAAAATGAAGTTTTAAATCAATCTAAAGTATATATGAGTAAACTTGGTCTGACAGTTACCAATGCTTAATCAGTGAGGCACCTATCTCAGCGATCTGTCTATTTCGTTCATCCATAGTTGCCTGACTCCCCGTCGTGTAGATAACTACGATACGGGAGGGCTTACCATCTGGCCCCAGTGCTGCAATGATACCGCGAGAGCCACGCTCACCGGCTCCAGATTTATCAGCAATAAACCAGCCAGCCGGAAGGGCCGAGCGCAGAAGTGGTCCTGCAACTTTATCCGCCTCCATCCAGTCTATTAATTGTTGCCGGGAAGCTAGAGTAAGTAGTTCGCCAGTTAATAGTTTGCGCAACGTTGTTGCCATTGCTACAGGCATCGTGGTGTCACGCTCGTCGTTTGGTATGGCTTCATTCAGCTCCGGTTCCCAACGATCAAGGCGAGTTACATGATCCCCCATGTTGTGCAAAAAAGCGGTTAGCTCCTTCGGTCCTCCGATCGTTGTCAGAAGTAAGTTGGCCGCAGTGTTATCACTCATGGTTATGGCAGCACTGCATAATTCTCTTACTGTCATGCCATCCGTAAGATGCTTTTCTGTGACTGGTGAGTACTCAACCAAGTCATTCTGAGAATAGTGTATGCGGCGACCGAGTTGCTCTTGCCCGGCGTCAATACGGGATAATACCGCGCCACATAGCAGAACTTTAAAAGTGCTCATCATTGGAAAACGTTCTTCGGGGCGAAAACTCTCAAGGATCTTACCGCTGTTGAGATCCAGTTCGATGTAACCCACTCGTGCACCCAACTGATCTTCAGCATCTTTTACTTTCACCAGCGTTTCTGGGTGAGCAAAAACAGGAAGGCAAAATGCCGCAAAAAAGGGAATAAGGGCGACACGGAAATGTTGAATACTCATACTCTTCCTTTTTCAATATTATTGAAGCATTTATCAGGGTTATTGTCTCATGAGCGGATACATATTTGAATGTATTTAGAAAAATAAACAAATAGGGGTTCCGCGCACATTTCCCCGAAAAGTGCCAGATACCTGAAACAAAACCCATCGTACGGCCAAGGAAGTCTCCAATAACTGTGATCCACCACAAGCGCCAGGGTTTTCCCAGTCACGACGTTGTAAAACGACGGCCAGTCATGCATAATCCGCACGCATCTGGAATAAGGAAGTGCCATTCCGCCTGACCT |
| pUC_M1  *M1RNA | AGTTCGGTGTAGGTCGTTCGCTCCAAGCTGGGCTGTGTGCACGAACCCCCCGTTCAGCCCGACCGCTGCGCCTTATCCGGTAACTATCGTCTTGAGTCCAACCCGGTAAGACACGACTTATCGCCACTGGCAGCAGCCACTGGTAACAGGATTAGCAGAGCGAGGTATGTAGGCGGTGCTACAGAGTTCTTGAAGTGGTGGCCTAACTACGGCTACACTAGAAGAACAGTATTTGGTATCTGCGCTCTGCTGAAGCCAGTTACCTTCGGAAAAAGAGTTGGTAGCTCTTGATCCGGCAAACAAACCACCGCTGGTAGCGGTGGTTTTTTTGTTTGCAAGCAGCAGATTACGCGCAGAAAAAAAGGATCTCAAGAAGATCCTTTGATCTTTTCTACGGGGTCTGACGCTCAGTGGAACGAAAACTCACGTTAAGGGATTTTGGTCATGAGATTATCAAAAAGGATCTTCACCTAGATCCTTTTAAATTAAAAATGAAGTTTTAAATCAATCTAAAGTATATATGAGTAAACTTGGTCTGACAGTTACCAATGCTTAATCAGTGAGGCACCTATCTCAGCGATCTGTCTATTTCGTTCATCCATAGTTGCCTGACTCCCCGTCGTGTAGATAACTACGATACGGGAGGGCTTACCATCTGGCCCCAGTGCTGCAATGATACCGCGAGACCCACGCTCACCGGCTCCAGATTTATCAGCAATAAACCAGCCAGCCGGAAGGGCCGAGCGCAGAAGTGGTCCTGCAACTTTATCCGCCTCCATCCAGTCTATTAATTGTTGCCGGGAAGCTAGAGTAAGTAGTTCGCCAGTTAATAGTTTGCGCAACGTTGTTGCCATTGCTACAGGCATCGTGGTGTCACGCTCGTCGTTTGGTATGGCTTCATTCAGCTCCGGTTCCCAACGATCAAGGCGAGTTACATGATCCCCCATGTTGTGCAAAAAAGCGGTTAGCTCCTTCGGTCCTCCGATCGTTGTCAGAAGTAAGTTGGCCGCAGTGTTATCACTCATGGTTATGGCAGCACTGCATAATTCTCTTACTGTCATGCCATCCGTAAGATGCTTTTCTGTGACTGGTGAGTACTCAACCAAGTCATTCTGAGAATAGTGTATGCGGCGACCGAGTTGCTCTTGCCCGGCGTCAATACGGGATAATACCGCGCCACATAGCAGAACTTTAAAAGTGCTCATCATTGGAAAACGTTCTTCGGGGCGAAAACTCTCAAGGATCTTACCGCTGTTGAGATCCAGTTCGATGTAACCCACTCGTGCACCCAACTGATCTTCAGCATCTTTTACTTTCACCAGCGTTTCTGGGTGAGCAAAAACAGGAAGGCAAAATGCCGCAAAAAAGGGAATAAGGGCGACACGGAAATGTTGAATACTCATACTCTTCCTTTTTCAATATTATTGAAGCATTTATCAGGGTTATTGTCTCATGAGCGGATACATATTTGAATGTATTTAGAAAAATAAACAAATAGGGGTTCCGCGCACATTTCCCCGAAAAGTGCCACCTGACGTCTAAGAAACCATTATTATCATGACATTAACCTATAAAAATAGGCGTATCACGAGGCCCTTTCGTCTCGCGCGTTTCGGTGATGACGGTGAAAACCTCTGACACATGCAGCTCCCGGAGACGGTCACAGCTTGTCTGTAAGCGGATGCCGGGAGCAGACAAGCCCGTCAGGGCGCGTCAGCGGGTGTTGGCGGGTGTCGGGGCTGGCTTAACTATGCGGCATCAGAGCAGATTGTACTGAGAGTGCACCATATGCGGTGTGAAATACCGCACAGATGCGTAAGGAGAAAATACCGCATCAGGCGCCATTCGCCATTCAGGCTGCGCAACTGTTGGGAAGGGCGATCGGTGCGGGCCTCTTCGCTATTACGCCAGCTGGCGAAAGGGGGATGTGCTGCAAGGCGATTAAGTTGGGTAACGCCAGGGTTTTCCCAGTCACGACGTTGTAAAACGACGGCCAGTGAATTCTAATACGACTCACTATAGAAGCTGACCAGACAGTCGCCGCTTCGTCGTCGTCCTCTTCGGGGGAGACGGGCGGAGGGGAGGAAAGTCCGGGCTCCATAGGGCAGGGTGCCAGGTAACGCCTGGGGGGGAAACCCACGACCAGTGCAACAGAGAGCAAACCGCCGATGGCCCGCGCAAGCGGGATCAGGTAAGGGTGAAAGGGTGCGGTAAGAGCGCACCGCGCGGCTGGTAACAGTCCGTGGCACGGTAAACTCCACCCGGAGCAAGGCCAAATAGGGGTTCATAAGGTACGGCCCGTACTGAACCCGGGTAGGCTGCTTGAGCCAGTGAGCGATTGCTGGCCTAGATGAATGACTGTCCACGACAGAACCCGGCTTATCGGTCAGTTTCACCTGATTTACGTCATCCGGATCCTCTAGAGTCGACCTGCAGGCATGCAAGCTTGGCGTAATCATGGTCATAGCTGTTTCCTGTGTGAAATTGTTATCCGCTCACAATTCCACACAACATACGAGCCGGAAGCATAAAGTGTAAAGCCTGGGGTGCCTAATGAGTGAGCTAACTCACATTAATTGCGTTGCGCTCACTGCCCGCTTTCCAGTCGGGAAACCTGTCGTGCCAGCTGCATTAATGAATCGGCCAACGCGCGGGGAGAGGCGGTTTGCGTATTGGGCGCTCTTCCGCTTCCTCGCTCACTGACTCGCTGCGCTCGGTCGTTCGGCTGCGGCGAGCGGTATCAGCTCACTCAAAGGCGGTAATACGGTTATCCACAGAATCAGGGGATAACGCAGGAAAGAACATGTGAGCAAAAGGCCAGCAAAAGGCCAGGAACCGTAAAAAGGCCGCGTTGCTGGCGTTTTTCCATAGGCTCCGCCCCCCTGACGAGCATCACAAAAATCGACGCTCAAGTCAGAGGTGGCGAAACCCGACAGGACTATAAAGATACCAGGCGTTTCCCCCTGGAAGCTCCCTCGTGCGCTCTCCTGTTCCGACCCTGCCGCTTACCGGATACCTGTCCGCCTTTCTCCCTTCGGGAAGCGTGGCGCTTTCTCATAGCTCACGCTGTAGGTATCTC |

**Supplementary Table S4. Sequences of tRNAs and tRNA arrays used in this study**

| tRNA(or pre-tRNA) | Sequence |
| --- | --- |
| tRNA^Ala^GGC | GGGGCUAUAGCUCAGCUGGGAGAGCGCUUGCAUGGCAUGCAAGAGGUCAGCGGUUCGAUCCCGCUUAGCUCCACCA |
| tRNA^Arg^CCG | GCGCCCGUAGCUCAGCUGGAUAGAGCGCUGCCCUCCGGAGGCAGAGGUCUCAGGUUCGAAUCCUGUCGGGCGCGCCA |
| tRNA^Asp^GUC | GGAGCGGUAGUUCAGUCGGUUAGAAUACCUGCCUGUCACGCAGGGGGUCGCGGGUUCGAGUCCCGUCCGUUCCGCCA |
| tRNA^Cys^GCA | GGCGCGUUAACAAAGCGGUUAUGUAGCGGAUUGCAAAUCCGUCUAGUCCGGUUCGACUCCGGAACGCGCCUCCA |
| tRNA^Gly^GCG | GCGGGAAUAGCUCAGUUGGUAGAGCACGACCUUGCCAAGGUCGGGGUCGCGAGUUCGAGUCUCGUUUCCCGCUCCA |
| tRNA^Glu^CUC | GUCCCCUUCGUCUAGAGGCCCAGGACACCGCCCUCUCACGGCGGUAACAGGGGUUCGAAUCCCCUAGGGGACGCCA |
| tRNA^His^GUG | GGUGGCUAUAGCUCAGUUGGUAGAGCCCUGGAUUGUGAUUCCAGUUGUCGUGGGUUCGAAUCCCAUUAGCCACCCCA |
| tRNA^Leu^CAG | GCGAAGGUGGCGGAAUUGGUAGACGCGCUAGCUUCAGGUGUUAGUGUCCUUACGGACGUGGGGGUUCAAGUCCCCCCCCUCGCACCA |
| tRNA^Lys^CUU | GGGUCGUUAGCUCAGUUGGUAGAGCAGUUGACUCUUAAUCAAUUGGUCGCAGGUUCGAAUCCUGCACGACCCACCA |
| tRNA^mMet^CAU | GGCUACGUAGCUCAGUUGGUUAGAGCACAUCACUCAUAAUGAUGGGGUCACAGGUUCGAAUCCCGUCGUAGCCACCA |
| tRNA^Phe^GAA | GCCCGGAUAGCUCAGUCGGUAGAGCAGGGGAUUGAAAAUCCCCGUGUCCUUGGUUCGAUUCCGAGUCCGGGCACCA |
| tRNA^Ser^GGA | GGUGAGGUGUCCGAGUGGCUGAAGGAGCACGCCUGGAAAGUGUGUAUACGGCAACGUAUCGGGGGUUCGAAUCCCCCCCUCACCGCCA |
| tRNA^Thr^GGU | GCUGAUAUGGCUCAGUUGGUAGAGCGCACCCUUGGUAAGGGUGAGGUCCCCAGUUCGACUCUGGGUAUCAGCACCA |
| tRNA^Tyr^GUA | GGUGGGGUUCCCGAGCGGCCAAAGGGAGCAGACUGUAAAUCUGCCGUCACAGACUUCGAAGGUUCGAAUCCUUCCCCCACCACCA |
| tRNA^Val^GAC | GCGUCCGUAGCUCAGUUGGUUAGAGCACCACCUUGACAUGGUGGGGGUCGGUGGUUCGAGUCCACUCGGACGCACCA |
| tRNA^Asn^GUU(U\|A) | UCCUCUGUAGUUCAGUCGGUAGAACGGCGGACUGUUAAUCCGUAUGUCACUGGUUCGAGUCCAGUCAGAGGAGCCA |
| tRNA^Asn^GUU(G\|C)  *Mutated base | GCCUCUGUAGUUCAGUCGGUAGAACGGCGGACUGUUAAUCCGUAUGUCACUGGUUCGAGUCCAGUCAGAGGCGCCA |
| tRNA^Asn^GUU(G\|A) | GCCUCUGUAGUUCAGUCGGUAGAACGGCGGACUGUUAAUCCGUAUGUCACUGGUUCGAGUCCAGUCAGAGGAGCCA |
| tRNA^Gln^CUG(U\|A) | UGGGGUAUCGCCAAGCGGUAAGGCACCGGAUUCUGAUUCCGGCAUUCCGAGGUUCGAAUCCUCGUACCCCAGCCA |
| tRNA^Gln^CUG(G\|C) | GGGGGUAUCGCCAAGCGGUAAGGCACCGGAUUCUGAUUCCGGCAUUCCGAGGUUCGAAUCCUCGUACCCCCGCCA |
| tRNA^Gln^CUG(G\|A) | GGGGGUAUCGCCAAGCGGUAAGGCACCGGAUUCUGAUUCCGGCAUUCCGAGGUUCGAAUCCUCGUACCCCAGCCA |
| tRNA^Ile^GAU(A\|U) | AGGCUUGUAGCUCAGGUGGUUAGAGCGCACCCCUGAUAAGGGUGAGGUCGGUGGUUCAAGUCCACUCAGGCCUACCA |
| tRNA^Ile^GAU(G\|C) | GGGCUUGUAGCUCAGGUGGUUAGAGCGCACCCCUGAUAAGGGUGAGGUCGGUGGUUCAAGUCCACUCAGGCCCACCA |
| tRNA^Ile^GAU(G\|U) | GGGCUUGUAGCUCAGGUGGUUAGAGCGCACCCCUGAUAAGGGUGAGGUCGGUGGUUCAAGUCCACUCAGGCCUACCA |
| tRNA^fMet^CAU(C\|A) | CGCGGGGUGGAGCAGCCUGGUAGCUCGUCGGGCUCAUAACCCGAAGAUCGUCGGUUCAAAUCCGGCCCCCGCAACCA |
| tRNA^fMet^CAU(G\|A) | GGCGGGGUGGAGCAGCCUGGUAGCUCGUCGGGCUCAUAACCCGAAGAUCGUCGGUUCAAAUCCGGCCCCCGCAACCA |
| tRNA^fMet^CAU(A\|A) | AGCGGGGUGGAGCAGCCUGGUAGCUCGUCGGGCUCAUAACCCGAAGAUCGUCGGUUCAAAUCCGGCCCCCGCAACCA |
| tRNA^fMet^CAU(G\|U) | GGCGGGGUGGAGCAGCCUGGUAGCUCGUCGGGCUCAUAACCCGAAGAUCGUCGGUUCAAAUCCGGCCCCCGCUACCA |
| tRNA^fMet^CAU(A\|U) | AGCGGGGUGGAGCAGCCUGGUAGCUCGUCGGGCUCAUAACCCGAAGAUCGUCGGUUCAAAUCCGGCCCCCGCUACCA |
| tRNA^Pro^GGG(C\|G) | CGGCACGUAGCGCAGCCUGGUAGCGCACCGUCAUGGGGUGUCGGGGGUCGGAGGUUCAAAUCCUCUCGUGCCGACCA |
| tRNA^Pro^GGG(G\|C) | GGGCACGUAGCGCAGCCUGGUAGCGCACCGUCAUGGGGUGUCGGGGGUCGGAGGUUCAAAUCCUCUCGUGCCCACCA |
| tRNA^Pro^GGG(G\|G) | GGGCACGUAGCGCAGCCUGGUAGCGCACCGUCAUGGGGUGUCGGGGGUCGGAGGUUCAAAUCCUCUCGUGCCGACCA |
| tRNA^Trp^CCA(A\|U) | AGGGGCGUAGUUCAAUUGGUAGAGCACCGGUCUCCAAAACCGGGUGUUGGGAGUUCGAGUCUCUCCGCCCCUGCCA |
| tRNA^Trp^CCA(G\|C) | GGGGGCGUAGUUCAAUUGGUAGAGCACCGGUCUCCAAAACCGGGUGUUGGGAGUUCGAGUCUCUCCGCCCCCGCCA |
| tRNA^Trp^CCA(G\|U) | GGGGGCGUAGUUCAAUUGGUAGAGCACCGGUCUCCAAAACCGGGUGUUGGGAGUUCGAGUCUCUCCGCCCCUGCCA |
| tRNA^Leu^CGU  *anti codon | GCGAAGGUGGCGGAAUUGGUAGACGCGCUAGCUUCGUGUGUUAGUGUCCUUACGGACGUGGGGGUUCAAGUCCCCCCCCUCGCACCA |
| tRNA^Thr^CGU | GCUGAUAUGGCUCAGUUGGUAGAGCGCACCCUUCGUAAGGGUGAGGUCCCCAGUUCGACUCUGGGUAUCAGCACCA |
| tRNA^Gln^ with the leader sequence  *leader sequence | GGGAGACCACAACGGUUUCCCUCUAGAUGGGGUAUCGCCAAGCGGUAAGGCACCGGAUUCUGAUUCCGGCAUUCCGAGGUUCGAAUCCUCGUACCCCAGCCA |
| tRNA^Ser^-HDVR  *HDVR | GGUGAGGUGUCCGAGUGGCUGAAGGAGCACGCCUGGAAAGUGUGUAUACGGCAACGUAUCGGGGGUUCGAAUCCCCCCCUCACCGCCAGGGUCGGCAUGGCAUCUCCACCUCCUCGCGGUCCGACCUGGGCUACUUCGGUAGGCUAAGGGAGAAG |
| tRNAs^Ala_Ser^ | GGGGCUAUAGCUCAGCUGGGAGAGCGCUUGCAUGGCAUGCAAGAGGUCAGCGGUUCGAUCCCGCUUAGCUCCACCAGGUGAGGUGUCCGAGUGGCUGAAGGAGCACGCCUGGAAAGUGUGUAUACGGCAACGUAUCGGGGGUUCGAAUCCCCCCCUCACCGCCA |
| tRNA^Ala_Ser^-HDVR  *T7hyb10 | GGGGCUAUAGCUCAGCUGGGAGAGCGCUUGCAUGGCAUGCAAGAGGUCAGCGGUUCGAUCCCGCUUAGCUCCACCAGGUGAGGUGUCCGAGUGGCUGAAGGAGCACGCCUGGAAAGUGUGUAUACGGCAACGUAUCGGGGGUUCGAAUCCCCCCCUCACCGCCAGGGUCGGCAUGGCAUCUCCACCUCCUCGCGGUCCGACCUGGGCUACUUCGGUAGGCUAAGGGAGAAGAGAUAACAGAUACUUCGGUAUCUGUUAUCUGUUUUUUUUCAACAGAUAGCCGCGUUCGCGCGGCUAUCUGUUUUUUUU |
| Pre-tRNA^GARCQ^-HDVR | GCGGGAAUAGCUCAGUUGGUAGAGCACGACCUUGCCAAGGUCGGGGUCGCGAGUUCGAGUCUCGUUUCCCGCUCCAGGGGCUAUAGCUCAGCUGGGAGAGCGCUUGCAUGGCAUGCAAGAGGUCAGCGGUUCGAUCCCGCUUAGCUCCACCAGCGCCCGUAGCUCAGCUGGAUAGAGCGCUGCCCUCCGGAGGCAGAGGUCUCAGGUUCGAAUCCUGUCGGGCGCGCCAGGCGCGUUAACAAAGCGGUUAUGUAGCGGAUUGCAAAUCCGUCUAGUCCGGUUCGACUCCGGAACGCGCCUCCAUGGGGUAUCGCCAAGCGGUAAGGCACCGGAUUCUGAUUCCGGCAUUCCGAGGUUCGAAUCCUCGUACCCCAGCCAGGGUCGGCAUGGCAUCUCCACCUCCUCGCGGUCCGACCUGGGCUACUUCGGUAGGCUAAGGGAGAAGAGAUAACAGAUACUUCGGUAUCUGUUAUCUGUUUUUUUUCAACAGAUAGCCGCGUUCGCGCGGCUAUCUGUUUUUUUU |
| Pre-tRNA^DfMHKV^-HDVR | GGAGCGGUAGUUCAGUCGGUUAGAAUACCUGCCUGUCACGCAGGGGGUCGCGGGUUCGAGUCCCGUCCGUUCCGCCACGCGGGGUGGAGCAGCCUGGUAGCUCGUCGGGCUCAUAACCCGAAGAUCGUCGGUUCAAAUCCGGCCCCCGCAACCAGGUGGCUAUAGCUCAGUUGGUAGAGCCCUGGAUUGUGAUUCCAGUUGUCGUGGGUUCGAAUCCCAUUAGCCACCCCAGGGUCGUUAGCUCAGUUGGUAGAGCAGUUGACUCUUAAUCAAUUGGUCGCAGGUUCGAAUCCUGCACGACCCACCAGCGUCCGUAGCUCAGUUGGUUAGAGCACCACCUUGACAUGGUGGGGGUCGGUGGUUCGAGUCCACUCGGACGCACCAGGGUCGGCAUGGCAUCUCCACCUCCUCGCGGUCCGACCUGGGCUACUUCGGUAGGCUAAGGGAGAAGAGAUAACAGAUACUUCGGUAUCUGUUAUCUGUUUUUUUUCAACAGAUAGCCGCGUUCGCGCGGCUAUCUGUUUUUUUU |
| Pre-tRNA^SmMFT^-HDVR | GGUGAGGUGUCCGAGUGGCUGAAGGAGCACGCCUGGAAAGUGUGUAUACGGCAACGUAUCGGGGGUUCGAAUCCCCCCCUCACCGCCAGGCUACGUAGCUCAGUUGGUUAGAGCACAUCACUCAUAAUGAUGGGGUCACAGGUUCGAAUCCCGUCGUAGCCACCAGCCCGGAUAGCUCAGUCGGUAGAGCAGGGGAUUGAAAAUCCCCGUGUCCUUGGUUCGAUUCCGAGUCCGGGCACCAGCUGAUAUGGCUCAGUUGGUAGAGCGCACCCUUGGUAAGGGUGAGGUCCCCAGUUCGACUCUGGGUAUCAGCACCAGGGUCGGCAUGGCAUCUCCACCUCCUCGCGGUCCGACCUGGGCUACUUCGGUAGGCUAAGGGAGAAGAGAUAACAGAUACUUCGGUAUCUGUUAUCUGUUUUUUUUCAACAGAUAGCCGCGUUCGCGCGGCUAUCUGUUUUUUUU |
| Pre-tRNA^LWY^-HDVR | GCGAAGGUGGCGGAAUUGGUAGACGCGCUAGCUUCAGGUGUUAGUGUCCUUACGGACGUGGGGGUUCAAGUCCCCCCCCUCGCACCAAGGGGCGUAGUUCAAUUGGUAGAGCACCGGUCUCCAAAACCGGGUGUUGGGAGUUCGAGUCUCUCCGCCCCUGCCAGGUGGGGUUCCCGAGCGGCCAAAGGGAGCAGACUGUAAAUCUGCCGUCACAGACUUCGAAGGUUCGAAUCCUUCCCCCACCACCAGGGUCGGCAUGGCAUCUCCACCUCCUCGCGGUCCGACCUGGGCUACUUCGGUAGGCUAAGGGAGAAGAGAUAACAGAUACUUCGGUAUCUGUUAUCUGUUUUUUUUCAACAGAUAGCCGCGUUCGCGCGGCUAUCUGUUUUUUUU |
| Pre-tRNA^IPEN^-HDVR | GGGCUUGUAGCUCAGGUGGUUAGAGCGCACCCCUGAUAAGGGUGAGGUCGGUGGUUCAAGUCCACUCAGGCCUACCACGGCACGUAGCGCAGCCUGGUAGCGCACCGUCAUGGGGUGUCGGGGGUCGGAGGUUCAAAUCCUCUCGUGCCGACCAGUCCCCUUCGUCUAGAGGCCCAGGACACCGCCCUCUCACGGCGGUAACAGGGGUUCGAAUCCCCUAGGGGACGCCAUCCUCUGUAGUUCAGUCGGUAGAACGGCGGACUGUUAAUCCGUAUGUCACUGGUUCGAGUCCAGUCAGAGGAGCCAGGGUCGGCAUGGCAUCUCCACCUCCUCGCGGUCCGACCUGGGCUACUUCGGUAGGCUAAGGGAGAAGAGAUAACAGAUACUUCGGUAUCUGUUAUCUGUUUUUUUUCAACAGAUAGCCGCGUUCGCGCGGCUAUCUGUUUUUUUU |
| Pre-tRNA^PIEN^-HDVR | GGGCACGUAGCGCAGCCUGGUAGCGCACCGUCAUGGGGUGUCGGGGGUCGGAGGUUCAAAUCCUCUCGUGCCGACCAAGGCUUGUAGCUCAGGUGGUUAGAGCGCACCCCUGAUAAGGGUGAGGUCGGUGGUUCAAGUCCACUCAGGCCUACCAGUCCCCUUCGUCUAGAGGCCCAGGACACCGCCCUCUCACGGCGGUAACAGGGGUUCGAAUCCCCUAGGGGACGCCAUCCUCUGUAGUUCAGUCGGUAGAACGGCGGACUGUUAAUCCGUAUGUCACUGGUUCGAGUCCAGUCAGAGGAGCCAGGGUCGGCAUGGCAUCUCCACCUCCUCGCGGUCCGACCUGGGCUACUUCGGUAGGCUAAGGGAGAAGAGAUAACAGAUACUUCGGUAUCUGUUAUCUGUUUUUUUUCAACAGAUAGCCGCGUUCGCGCGGCUAUCUGUUUUUUUU |
| Pre-tRNA^EIPN^-HDVR | GUCCCCUUCGUCUAGAGGCCCAGGACACCGCCCUCUCACGGCGGUAACAGGGGUUCGAAUCCCCUAGGGGACGCCAAGGCUUGUAGCUCAGGUGGUUAGAGCGCACCCCUGAUAAGGGUGAGGUCGGUGGUUCAAGUCCACUCAGGCCUACCACGGCACGUAGCGCAGCCUGGUAGCGCACCGUCAUGGGGUGUCGGGGGUCGGAGGUUCAAAUCCUCUCGUGCCGACCAUCCUCUGUAGUUCAGUCGGUAGAACGGCGGACUGUUAAUCCGUAUGUCACUGGUUCGAGUCCAGUCAGAGGAGCCAGGGUCGGCAUGGCAUCUCCACCUCCUCGCGGUCCGACCUGGGCUACUUCGGUAGGCUAAGGGAGAAGAGAUAACAGAUACUUCGGUAUCUGUUAUCUGUUUUUUUUCAACAGAUAGCCGCGUUCGCGCGGCUAUCUGUUUUUUUU |
| Pre-tRNA^NIPE^-HDVR | GCCUCUGUAGUUCAGUCGGUAGAACGGCGGACUGUUAAUCCGUAUGUCACUGGUUCGAGUCCAGUCAGAGGAGCCAAGGCUUGUAGCUCAGGUGGUUAGAGCGCACCCCUGAUAAGGGUGAGGUCGGUGGUUCAAGUCCACUCAGGCCUACCACGGCACGUAGCGCAGCCUGGUAGCGCACCGUCAUGGGGUGUCGGGGGUCGGAGGUUCAAAUCCUCUCGUGCCGACCAGUCCCCUUCGUCUAGAGGCCCAGGACACCGCCCUCUCACGGCGGUAACAGGGGUUCGAAUCCCCUAGGGGACGCCAGGGUCGGCAUGGCAUCUCCACCUCCUCGCGGUCCGACCUGGGCUACUUCGGUAGGCUAAGGGAGAAGAGAUAACAGAUACUUCGGUAUCUGUUAUCUGUUUUUUUUCAACAGAUAGCCGCGUUCGCGCGGCUAUCUGUUUUUUUU |
| Pre-21tRNA^DSGLP^-HDVR *DSGLP:DfMHKVSmMFTGARCQLWYPIEN | GGAGCGGUAGUUCAGUCGGUUAGAAUACCUGCCUGUCACGCAGGGGGUCGCGGGUUCGAGUCCCGUCCGUUCCGCCACGCGGGGUGGAGCAGCCUGGUAGCUCGUCGGGCUCAUAACCCGAAGAUCGUCGGUUCAAAUCCGGCCCCCGCAACCAGGUGGCUAUAGCUCAGUUGGUAGAGCCCUGGAUUGUGAUUCCAGUUGUCGUGGGUUCGAAUCCCAUUAGCCACCCCAGGGUCGUUAGCUCAGUUGGUAGAGCAGUUGACUCUUAAUCAAUUGGUCGCAGGUUCGAAUCCUGCACGACCCACCAGCGUCCGUAGCUCAGUUGGUUAGAGCACCACCUUGACAUGGUGGGGGUCGGUGGUUCGAGUCCACUCGGACGCACCAGGUGAGGUGUCCGAGUGGCUGAAGGAGCACGCCUGGAAAGUGUGUAUACGGCAACGUAUCGGGGGUUCGAAUCCCCCCCUCACCGCCAGGCUACGUAGCUCAGUUGGUUAGAGCACAUCACUCAUAAUGAUGGGGUCACAGGUUCGAAUCCCGUCGUAGCCACCAGCCCGGAUAGCUCAGUCGGUAGAGCAGGGGAUUGAAAAUCCCCGUGUCCUUGGUUCGAUUCCGAGUCCGGGCACCAGCUGAUAUGGCUCAGUUGGUAGAGCGCACCCUUGGUAAGGGUGAGGUCCCCAGUUCGACUCUGGGUAUCAGCACCAGCGGGAAUAGCUCAGUUGGUAGAGCACGACCUUGCCAAGGUCGGGGUCGCGAGUUCGAGUCUCGUUUCCCGCUCCAGGGGCUAUAGCUCAGCUGGGAGAGCGCUUGCAUGGCAUGCAAGAGGUCAGCGGUUCGAUCCCGCUUAGCUCCACCAGCGCCCGUAGCUCAGCUGGAUAGAGCGCUGCCCUCCGGAGGCAGAGGUCUCAGGUUCGAAUCCUGUCGGGCGCGCCAGGCGCGUUAACAAAGCGGUUAUGUAGCGGAUUGCAAAUCCGUCUAGUCCGGUUCGACUCCGGAACGCGCCUCCAUGGGGUAUCGCCAAGCGGUAAGGCACCGGAUUCUGAUUCCGGCAUUCCGAGGUUCGAAUCCUCGUACCCCAGCCAGCGAAGGUGGCGGAAUUGGUAGACGCGCUAGCUUCAGGUGUUAGUGUCCUUACGGACGUGGGGGUUCAAGUCCCCCCCCUCGCACCAAGGGGCGUAGUUCAAUUGGUAGAGCACCGGUCUCCAAAACCGGGUGUUGGGAGUUCGAGUCUCUCCGCCCCUGCCAGGUGGGGUUCCCGAGCGGCCAAAGGGAGCAGACUGUAAAUCUGCCGUCACAGACUUCGAAGGUUCGAAUCCUUCCCCCACCACCAGGGCACGUAGCGCAGCCUGGUAGCGCACCGUCAUGGGGUGUCGGGGGUCGGAGGUUCAAAUCCUCUCGUGCCGACCAAGGCUUGUAGCUCAGGUGGUUAGAGCGCACCCCUGAUAAGGGUGAGGUCGGUGGUUCAAGUCCACUCAGGCCUACCAGUCCCCUUCGUCUAGAGGCCCAGGACACCGCCCUCUCACGGCGGUAACAGGGGUUCGAAUCCCCUAGGGGACGCCAUCCUCUGUAGUUCAGUCGGUAGAACGGCGGACUGUUAAUCCGUAUGUCACUGGUUCGAGUCCAGUCAGAGGAGCCAGGGUCGGCAUGGCAUCUCCACCUCCUCGCGGUCCGACCUGGGCUACUUCGGUAGGCUAAGGGAGAAGAGAUAACAGAUACUUCGGUAUCUGUUAUCUGUUUUUUUUCAACAGAUAGCCGCGUUCGCGCGGCUAUCUGUUUUUUUU |

**Supplementary Table S5. Composition of the PURE system**

| Component | Concentration | Component | Concentration |
| --- | --- | --- | --- |
| Initiation Factor 1 | 25 µM | Tryptophanyl-tRNA Synthetase | 28 nM |
| Initiation Factor 2 | 1.0 µM | Tyrosyl-tRNA Synthetase | 0.15 µM |
| Initiation Factor 3 | 4.9 µM | Valyl-tRNA Synthetase | 17 nM |
| Elongation Factor G | 1.1 µM | Methionyl-tRNA Formyltransferase | 0.59 µM |
| Elongation Factor Tu | 80 µM | Myokinase | 1.4 µM |
| Elongation Factor Ts | 3.3 µM | Creatine kinase | 0.25 µM |
| Release Factor 1 | 49 nM | Nucleoside diphosphate kinease | 16 nM |
| Release Factor 2 | 48 nM | Pyrophosphatase | 41 nM |
| Release Factor 3 | 0.17 µM | Trigger Factor | 1.0 µM |
| Ribosome Recycling Factor | 3.9 µM | E. coli DEAH type RNA helicase A | 63 nM |
| Alanyl-tRNA Synthetase | 0.73 µM | Ribosome | 1.0 µM |
| Arginyl-tRNA Synthetase | 31 nM | Tyrosine | 0.30 mM |
| Asparaginyl-tRNA Synthetase | 0.42 µM | Cysteine | 0.30 mM |
| Asparagyl-tRNA Synthetase | 0.12 µM | 18 other amino acids | 0.36 mM |
| Cysteinyl-tRNA Synthetase | 24 nM | ATP | 0.38* mM |
| Glutaminyl-tRNA Synthetase | 60 nM | GTP | 0.25* mM |
| Glutamyl-tRNA Synthetase | 0.23 µM | CTP | 0.13* mM |
| Glycyl-tRNA Synthetase | 86 nM | UTP | 0.13* mM |
| Histidyl-tRNA Synthetase | 85 nM | N-2-hydroxyethylpiperazine-N'-2-ethanesulfonic acid (pH7.6) | 0.10 M |
| Isoleucyl-tRNA Synthetase | 0.37 µM | Glutamic acid potassium salt | 70 mM |
| Leucyl-tRNA Synthetase | 41 nM | Spermidine | 0.375* mM |
| Lysyl-tRNA Synthetase | 0.12 µM | Creatine phosphate | 25 mM |
| Methionyl-tRNA Synthetase | 0.11 µM | Dithiothreitol | 6 mM |
| Phenylalanyl-tRNA Synthetase | 0.13 µM | 10-formyl-5,6,7,8-tetrahydro folic acid | 10 µg/mL |
| Prolyl-tRNA Synthetase | 0.17 µM | Yeast inorganic pyrophosphatase (NEB) | 0.2 units/mL |
| Seryl-tRNA Synthetase | 78 nM | RNase Plus Inhibitor (Promega) | 0.1 U/µL |
| Threonyl-tRNA Synthetase | 84 nM | T7 RNA polymerase (Takara) | 0.42* U/µL |

*For composition A (phi29)

**Supplementary Table S6. Optimized PURE system compositions for each experiment**

| Component | PURE system composition | | | | | |
| --- | --- | --- | --- | --- | --- | --- |
|  | A(phi29) | B(RNaseP) | C(5’-G) | D(HDVR) | E(RNaseP+HDVR) | F(21) |
| NTP (mM) | 0.88 | 3.1 | 2.0 | 3.1 | 3.1 | 7.4 |
| Mg(OAc)2 (mM) | 7.9 | 13 | 15 | 15 | 16 | 16 |
| Spermidine (mM) | 0.38 | 0.75 | 0.38 | 0.38 | 0.75 | 0.75 |
| T7 RNAP (U/µl) | 0.42 | 1.7 | 1.7 | 1.7 | 1.7 | 3.4 |

### Supplementary methods: preparation method of the PURE system

**Strains**

E. coli strain A19 was used for ribosome purification. For other protein components, an E. coli strain, BW25113ΔuidA(DE3)/pREP4, was used after introducing an expression plasmid for the target gene. The plasmids were previously reported (1). The strain was constructed from JW1609 in the KEIO collection (2) by removing the Km marker and then introducing λDE3 prophage, which expresses T7 RNA polymerase, and a plasmid (pREP4), which expresses lacI. The reason for using ΔuidA strain is because we previously used glucuronidase (uidA) as a reporter gene.

**Culture for ribosome purification**

For ribosome, fresh colonies of E. coli A19 strain were inoculated in 5 mL of LB medium and incubated at 37°C for 8 h. The culture was added to 100 mL LB medium and incubated at 37°C overnight. The culture was poured into a fermenter (Bioneer-500, B. E. MARUBISHI Co., Ltd., Japan) that contained 2.5 L of a prewarmed fermenter medium for ribosome and incubated at 37°C for 4-5 h with mixing at 900 rpm and an air flow (4) until OD600 reached 7.5. During incubation, the pH was maintained at approximately 7 using 6 N NaOH. The cells (approximately 60 g) were collected by centrifugation, quickly frozen in liquid nitrogen, and stored at -80°C.

The fermenter medium for ribosome was prepared as follows. Solution 1 was prepared by mixing yeast extract (17.5 g), NaCl (2.5 g), (NH_4_)_2_SO_4_ (2.5 g), CaCl_2_·2H_2_O (2.1 g), FeSO_4_·7H_2_O (1.25 g), K_2_SO_4_ (1.5 g), CoCl_2_·6H_2_O (12.5 mg), CuSO_4_·5H_2_O (75 mg), Na_2_MoO_4_·2H_2_O (12.5 mg), Mn(OAc)_2_·4H_2_O (60 mg), and ZnSO_4_·7H_2_O (163 mg) in 2200 mL tap water. Solution 2 was prepared by dissolving D(+)-glucose (25 g) and thiamine-HCl (25 mg) in 62.5 mL tap water. Solution 3 was prepared by dissolving KH_2_PO_4_ (4.25 g) and (NH_4_)_2_HPO_4_ (27.5 g) in distilled water (188 mL). Solution 4 was prepared by dissolving MgSO_4_·7H_2_O (6.25 g) in 62.5 mL tap water. After autoclaving (121°C for 20 min) each solution, all solutions were mixed in the fermenter.

**Culture for the translational proteins**

For other protein components, after introducing each expression plasmid into the E. coli strain, the colonies were inoculated into 5 mL of LB medium containing ampicillin (50 µg/mL) and kanamycin (50 µg/mL) and incubated at 37°C for 8 h. The culture was added to 100 mL of LB medium containing ampicillin and kanamycin at the same concentrations and incubated at 37°C overnight. The cells were collected by centrifugation and resuspended in 100 mL fresh LB medium. The cell suspension was poured into a fermenter (Bioneer-500, B. E. MARUBISHI Co., Ltd., Japan) containing 2 L of prewarmed fermenter medium and incubated at 37°C for 6 h with mixing at 900 rpm and air flow of 4. Next, 600 µL of 500 mg/ml ampicillin was added to the culture. The feeding medium 1 was attached to the fermenter, and the culture was further incubated at 37°C overnight with supplying the feeding medium 1 at a rate of a few drops per 10 s. During incubation, the pH was maintained at approximately 7 using 6 N NaOH. The next day, 1.125 mL of 500 mg/ml ampicillin and 1 M IPTG (2.5 mL) was added to the culture. The feeding medium 2 was attached to the fermenter, and the culture was further incubated at 37°C for 4 h with supplying the feeding medium 2 at a rate of a few drops per 4 s. If the Do value was greater than 10% during incubation, the feeding speed decreased. Then, the cells (approximately 300 g) were collected by centrifugation, quickly frozen in liquid nitrogen, and stored at -80°C.

The fermenter medium for translational proteins was prepared as follows: Solution 1 was prepared by mixing yeast extract (14 g), NaCl (2 g), (NH_4_)_2_SO_4_ (2 g), CaCl_2_ (1.28 g), FeSO_4_·7H_2_O (1 g), K_2_SO_4_ (1.2 g), CoCl_2_·6H_2_O (10 mg), CuSO_4_·5H_2_O (60 mg), Na_2_MoO_4_·2H_2_O (10 mg), Mn(OAc)_2_·4H_2_O (52 mg), and ZnSO_4_·7H_2_O (130 mg) in 1750 mL of tap water. Solution 2 was prepared by dissolving D(+)-glucose (20 g) and thiamine-HCl (20 mg) in 50 mL tap water. Solution 3 was prepared by dissolving KH_2_PO_4_ (3.4 g) and (NH_4_)_2_HPO_4_ (22 g) in distilled water (150 mL). Solution 4 was prepared by dissolving MgSO_4_ · 7H_2_O (5 g) in tap water (50 mL). After autoclaving (121°C for 20 min) each solution, all solutions were mixed in the fermenter with 0.4 mL of 500 mg/ml ampicillin and 2 mL of 50 mg/ml kanamycin.

The feeding medium 1 was prepared as follows. Solution 1 was prepared by dissolving yeast extract (20 g) in 100 mL of tap water. Solution 2 was prepared by dissolving D(+)-glucose (100 g) and thiamine-HCl (10 mg) in tap water (150 mL). After autoclaving (121°C for 20 min) each solution, the two solutions, 175 µL of 500 mg/ml ampicillin, and 250 µL of 50 mg/ml kanamycin were mixed just before use.

The feeding medium 2 was prepared as follows. Solution 1 was prepared by dissolving yeast extract (100 g) in 200 mL of tap water. Solution 2 was prepared by dissolving D(+)-glucose (80 g) and thiamine-HCl (10 mg) in tap water (50 mL). After autoclaving (121°C for 20 min) each solution, the two solutions, 350 µL of 500 mg/ml ampicillin, and 250 µL of 50 mg/ml kanamycin were mixed just before use.

Purification of ribosome

The frozen cells (~10 g) were mixed with two volumes (~20 mL) of S20 buffer. Three times the amount of 0.1 mm glass beads (~30 g) was added to the cell mixture and vigorously shaken with Multi-beads Shoker (Yasui Kikai, Japan) at 2500 rpm for 30 s for 3 cycles at 0°C to disrupt the cells. After removing cell debris and beads by centrifugation at 6 k×g for 20 min at 4°C, the supernatant was further centrifuged at 18 krpm for 30 h at 4°C with R10A2 rotor (Himac). The supernatant was centrifuged again at 18 krpm for 30 min at 4°C with R20A2 rotor (Himac). Ammonium sulfate (0.222 g per 1 mL of the supernatant) was dissolved in the supernatant with stirring at 4°C, stand for 20 min at at 4°C, and centrifuged at 18 krpm for 60 min at 4°C with R20A2 rotor (Himac). The supernatant was filtered through a 0.45 µm filter and applied to HiTrap Butyl FF column (5 mL ×4, Cytiva) at 2 mL/min using Akta go (Cytiva). The column was washed at 1 mL/min with 30 mL of butyl-column buffer A. Then, the column was washed at 2 mL/min with 60 mL of a solution (80% butyl-column buffer A and 20% butyl-column buffer B), followed by elution with a solution at 2 mL/min with 40 mL of a solution (50% butyl-column buffer A and 50% butyl-column buffer B). The fractions around the A260 peak were collected. The collected fractions (~10 mL) were placed on 30 mL of 30% sucrose buffer and centrifuged at 30 krpm for 17 h at 4°C with 45Ti rotor (Beckman). The precipitate was dissolved in 70S buffer and concentrated using an Amicon Ultra (Millipore, 100 kDa cut). The final solution was quickly frozen in liquid nitrogen, and stored at -80°C. The protein concentration was calculated using the A260 value.

S20 buffer contained 10 mM HEPES-KOH (pH7.6), 10 mM MgCl_2_, 10 mM KCl, and 1 mM DTT. Butyl-column buffer A contains 20 mM HEPES-KOH (pH7.6), 10 mM Mg(OAc)_2_, 1.5 M (NH_4_)_2_SO_4_, and 7 mM 2-mercaptoethanol. Butyl column buffer B contained 20 mM HEPES-KOH (pH7.6), 10 mM Mg(OAc)_2_, and 7 mM 2-mercaptoethanol. The 30% sucrose buffer contained 20 mM HEPES-KOH (pH7.6), 10 mM Mg(OAc)_2_, 30 mM (NH_4_)Cl, 30% sucrose, and 7 mM 2-mercaptoethanol. 70S buffer contained 20 mM HEPES-KOH (pH7.6), 6 mM Mg(OAc)_2_, 30 mM KCl, and 7 mM 2-mercaptoethanol.

**Purification of the translational proteins**

The frozen cells (20-40 g) were dissolved in three volumes (60-120 mL) of Ni-NTA buffer A. The same amount of glass beads 0.1 mm (20-40 g) was added to the cell suspension and vigorously shake with Multi-beads Shoker (Yasui Kikai, Japan) at 2500 rpm 30 sec for 3 cycles at 0°C to disrupt the cells. After removing cell debris and the beads by centrifugation at 8 k×g for 10 min at 4°C, the supernatant was further centrifuged at 40 krpm for 1 h at 4°C with 45Ti rotor (Beckman). The supernatant was diluted up to 100 mL with Ni-NTA buffer A and centrifuged at 40 krpm for 1 h at 4°C with 45Ti rotor again. After filtering with 0.45 µm filter, the supernatant was applied to HisTrap column (5 mL ×3, Cytiva) at 2.5 mL/min using Akta go (Cytiva). The column was washed at 2 mL/min with 60 mL of a solution (95% Ni-NTA buffer A and 5% Ni-NTA buffer B). The target protein was eluted at 2 mL/min with a linear gradient of 5% to 50% Ni-NTA buffer B for 30 min. Fractions containing the target proteins (typically ~12 mL) were collected and concentrated using an Amicon Ultra (Millipore). The pore size of Amicon Ultra was determined based on the size of the target protein. The protein solution was then applied to HiLoad® 16/600 Superdex 200 pg (Cytiva) using Akta Go and eluted with 150 mL of gel-filtration buffer. Fractions containing the target proteins (typically ~12 mL) were collected and concentrated using an Amicon Ultra (Millipore). The final solution was mixed with 2× glycerol stock buffer, quickly frozen in liquid nitrogen, and stored at -80°C. The protein concentration was calculated from the A280 value and the absorbance was estimated using ProtParam (3).

The Ni-NTA buffer A contained 50 mM HEPES-KOH (pH7.6), 10 mM MgCl_2_, 1 M NH_4_Cl, 15% glycerol, and 7 mM 2-mercaptoethanol. When purifying EF-Tu, 10 µM GDP was additionally supplied at 10 µM. The Ni-NTA buffer B contained 50 mM HEPES-KOH (pH7.6), 10 mM MgCl_2_, 100 mM KCl, 500 mM imidazole, 15% glycerol, and 7 mM 2-mercaptoethanol. When purifying EF-Tu, 10 µM GDP was additionally supplied at 10 µM. The gel filtration buffer contained 50 mM HEPES-KOH (pH7.6), 10 mM MgCl_2_, 100 mM KCl, and 7 mM 2-mercaptoethanol. The 2× glycerol stock buffer contained 50 mM HEPES-KOH (pH7.6), 10 mM MgCl_2_, 100 mM KCl, 60% glycerol, and 7 mM 2-mercaptoethanol.

**References**

1. Shimizu,Y., Inoue,A., Tomari,Y., Suzuki,T., Yokogawa,T., Nishikawa,K. and Ueda,T. (2001) Cell-free translation reconstituted with purified components. Nat Biotechnol, 19.

2. Baba T. et al. (2006) Mol Systems Biol, doi:10.1038/msb4100050.

3. Gasteiger E., Hoogland C., Gattiker A., Duvaud S., Wilkins M.R., Appel R.D., Bairoch A.; Protein Identification and Analysis Tools on the Expasy Server; (In) John M. Walker (ed): The Proteomics Protocols Handbook, Humana Press (2005). pp. 571-607
